# Identification of type III polyketide synthases from Ginger for dehydrogingerdione and curcumin biosynthesis by engineered *Escherichia coli*

**DOI:** 10.64898/2026.07.16.739013

**Authors:** Eliana L. Peña, Sun-Young Kang, François Gaascht, Claudia Schmidt-Dannert

**Affiliations:** Department of Biochemistry, Molecular Biology, & Biochemistry, University of Minnesota, Minneapolis, MN 55455, USA; BioTechnology Institute, University of Minnesota, St. Paul, MN 55108, USA

**Author notes:** Corresponding author: Prof. Claudia Schmidt-Dannert, 140 Gortner Laboratory, University of Minnesota, 1479 Gortner Avenue, St. Paul, MN 55108.

**Keywords:** Ginger, 6-dehydrogingerdione, polyketide synthase, curcumin, biosynthesis, biomanufacturing

## Abstract

Many valuable plant metabolites are synthesized by type III polyketide synthases (PKS) that have become targets for the engineering of microbial production systems of these compounds. The rhizomes of turmeric (*Curcuma longa*) and ginger (*Zingiber officinalis*) are highly regarded for medicinal and culinary purposes and are the sources of bioactive curcuminoid and gingeroid polyketides. Fast growing demand for these compounds has sparked effort to identify their biosynthetic pathways to facilitate their heterologous production. In turmeric, a collaborative diketide synthase (DCS) and PKS (CURS) pair synthesizes curcumin from feruloyl- and malonyl-CoA. Yet, *bona fide* genes for the biosynthesis of gingeroids in Ginger are not known. Here we report the identification of two DCS/PKS pairs in Ginger that have different activity profiles in *E. coli* engineered to provide feruloyl- and hexanoyl-CoA as substrates. We show that one PKS (ZoPKS2) makes 6-dehydrogingerdione (6-DHG) as its major product while the other PKS (ZoPKS1) is a curcumin synthase. We found that *Zo*PKS2 becomes an efficient curcumin synthase when hexanoyl-CoA is not available, making it a dual-function enzyme that can be used to easily switch heterologous productions towards either of these two valuable products. Precursor feeding studies show that the substrate promiscuity of the collaborative DCS/PKSs may be exploited to access different dehydrogingerdione derivatives, while structural models of the Ginger PKSs offer insights for future engineering of product profiles. We believe that this work will add to the type III PKS toolbox and enable the development of efficient production platforms for gingeroids.

## INTRODUCTION

Plants produce a plethora of secondary metabolites, including the large class of polyketides ^1^. These compounds have a long tradition of being used as medicines, pigments, food additives, pesticides and more recently, they have also attracted significant interest as platform chemicals for biomanufacturing ^2–3^. Unlike the large mega-synthases involved in microbial polyketide biosynthesis, plant polyketides are synthesized by ∼40 kDa homo-dimeric type III polyketide synthases (PKS) that catalyze the stepwise condensation of starter and extender acyl-CoA units into various scaffolds. Because of their simpler structure, plant PKSs have been extensively explored for the biosynthesis of valuable molecules by engineered microorganisms, including by our group ^2, 4–7^. In addition, dozens of functionally distinct type III PKSs have been characterized since the first structure of a chalcone synthase (CHS) was published in 1999 ^8^. These enzymes use substrates other than coumaroyl- and malonyl-CoA used by CHS and vary in the number of condensations and cyclization reactions they perform ^9^, providing biosynthetic routes to numerous natural and unnatural molecules via metabolic pathway and enzyme engineering.

Among the known plant polyketide pathways, the biosynthesis of curcumin stands out as it depends on the synergistic activities of two different type III PKSs identified in the rhizomes of *Curcuma longa* (a member of the ginger family). Here, a diketide synthase (DCS) first condenses feruloyl-CoA (starter) with malonyl-CoA (extender) to produce feruloyldiketide-CoA, which becomes the extender substrate for a second PKS, curcumin synthase (CURS1), that catalyzes the decarboxylative condensation with feruloyl-CoA to yield curcumin ^10–11^. Subsequently, two additional PKSs (CURS2 & 3) with slightly different substrate preferences were identified^12^. Downstream tailoring steps transform curcumin than into curcuminoids that together constitute a major fraction of the biological active compounds found in the yellow spice turmeric ^13^.

Interestingly, a distantly related PKS from rice (*Oryza sativa*) (CUS) was found to synthesize bisdemethoxycurcumin from coumaroyl- and malonyl-CoA via a β-keto acid intermediate that serves as extender for a second reaction ^14^. Because of a fast growing demand for curcumoids that cannot be met by harvesting rhizomes alone, microbial systems have been engineered for curcumin production. DCS/CURS or CUS expression in engineered *E. coli* yielded titers (100-500 mg L^-1^) that reached levels obtained for the chalcone naringenin ^13, 15–17^.

Much like turmeric, the rhizomes of *Zingiber officinale,* Ginger, are highly regarded for cooking and medicinal use. Many of ginger’s pharmaceutical properties are attributed to gingerols and their precursor, dehydrogingerdione ^18–21^. Yet, *bona fide* genes for the biosynthesis of these natural products in Ginger are not known. Recent studies correlated the upregulation of several type III PKS genes with high concentrations of dehydrogingerdione and gingerol compounds in the plants’ rhizomes ^22–23^, suggesting a biosynthetic route that is similar to curcumin except that the gingerdione PKS would use medium chain fatty acyl-CoA’s as starter unit instead of the feruloy-CoA used by CURS (**Figure 1**). Alternatively, a single enzyme could carry out this reaction as shown for CUS, which was expressed in *E. coli* with an engineered β-oxidation pathway to produce 3-oxoacyl CoA starter substrates for condensation with coumaryl-CoA to form dehydrogingerdiones ^24^. In fact, CUS has been extensively investigated for the *in vivo* or *in vitro* synthesis of curcuminoid and gingeroid derivatives from phenlypropionic acids and/or 3-oxo-fatty acids or their CoA analoges ^12, 14, 16, 24–25^.

**Figure 1:**
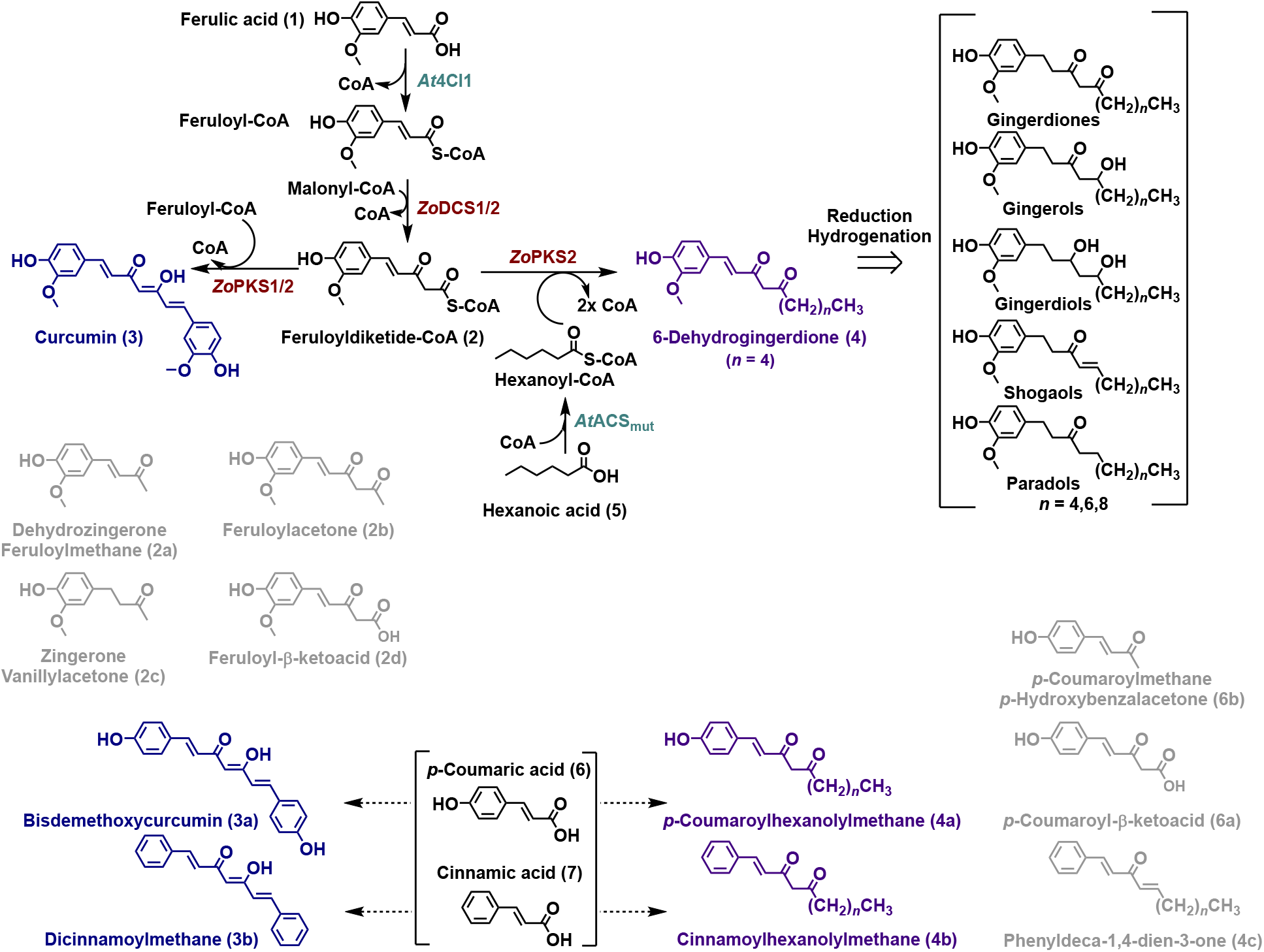
Designed biosynthetic pathway to 6-DHG and curcumin in *E. coli*. **(a)** Ginger diketide synthases (*Zo*DCS1 and 2) condense the starter substrate feruloyl-CoA with malonyl-CoA to yield feruloyldiketide-CoA (**2**), which becomes the extender substrate for a second set of ginger enzymes (*Zo*PKS1 and 2) that synthesize curcumin (**3**) or 6-DHG (6-DHG) (**4**) by decarboxylative condensation with feruloyl-CoA or hexanoyl-CoA as starter substrates. Only *Zo*PKS2 (GINS) synthesizes 6-DHG. CoA starter substrates are synthesized in *E. coli* cells by *Arabidopsis thaliana* CoA-ligases *At*4CL1 and *At*ACSmut from exogenously supplied ferulic acid (**1**) and hexanoic acid (**5**). Subsequent reduction and hydrogenation of 6-DHG and/or short chain fatty acyl-CoA substrates with varying chain lengths give rise to diverse gingeroid compounds. The feruloyldiketide intermediate (**2**), curcumin (**3**) and 6-DHG (**4**) are known to transform non-enzymatically to compounds such as **2a-c**. **(b)** Replacing ferulic acid with coumaric acid (**6**) or cinnamic acid (**7**) can lead to curcumin (**3a**, **3b**) and 6-DHG (**4a**, **4b**) derivatives via their corresponding coumaroyl- or cinnamoyldiketide-CoA intermediates (not shown). The diketide intermediates and products are known to degrade and transform into different compounds, with some relevant examples shown in gray (**2a-d**, **6a**, **6b**, **4c**).

Here we report the identification of two chromosomally clusterd DCS/PKS pairs (*Zo*DCS1,2 and *Zo*PKS1,2) from Ginger (*Zo*) with differet activity profiles when expressed in *E. coli* engineered to produce feruloyl-CoA and hexanoyl-CoA as substrates. Using a bi-directional-plasmid system desinged for the combinatorial co-expression of the DCS and PKS genes, *Zo*PKS2 synthesized 6-dehydrogingerdione (6-DHG) as its major product while *Zo*PKS1 produced curcumin when fed with ferulic and hexanoic acid. Suprisingly, *Zo*PKS2 becomes an efficient curcumin synthases when hexanoyl-CoA is not available, making it a dual-function enzyme that can be used to easily switch biosynthesis towards either of these two valuable natural products. In addition, precursor feeding studies show that the substrate promiscuity of the collaborative DCS/PKSs can be exploited to access different dehydrogingerdione derivatives, while structural models of the Ginger PKSs suggest opportunities for future engineering of product profiles. Together, these results lay the foundation for the microbial production of highly valuable gingeroids by applying the same metabolic engineering strategies previously reported for curcuminoids.

## RESULTS AND DISCUSSION

### Identification of 6-DHG biosynthetic gene candidates

Sequencing of the haplotype-resolved genome of the diploid *Zingiber officinalis* along with transcriptome and metabolomic analysis of its rhizomes was initially published by Li et. al. in Horticulture Research (a Nature publication) ^23^ without providing any supporting sequence data. Yet, the authors claimed the identification of candidate gingerol biosynthetic genes, including 17 type III PKS sequences of which four appear to be overexpressed in rhizome developmental stages associated with high 6,10-gingerdione and 6-gingerol content in these tissues. Subsequently, a different group sequenced the ginger genome and performed allelic gene expression studies for chromosome level genome assembly which was submitted as a BioProject to NCBI (original submission: PRJNA 647255, NCBI created RefSeq: PRJNA736965) ^26^.

We searched in the diploid ginger genome assembly deposited in the NCBI database for homologs of the characterized *Curcuma longa* DCS and PKS enzymes (CURS1-3), assuming that 6-DHG biosynthesis would also involve a collaborative DCS/PKS pair. In addition, we included the bisdemethoxycurcumin synthase CUS (*Oryza sativa*) and chalcone synthase CHS (*Arabidopsis thaliana*) in our analysis. Our search identified 29 putative DCS/PKS sequences of which 16 were annotated in the original BioProject assembly and 13 additional sequences annotated in NCBI’s RefSeq sequence (**Table S1)**. Phylogenetic analysis revealed three major clades composed of CHS, DCS and CURS-like sequences, with CUS as its own branch (**Figure 2a**). Each Ginger sequence (labeled: Zo) was represented by at least two chromosomal alleles and chromosome (Chr) 9 appeared to be a “hotspot” for their genomic location. Zo homologs were identified for each of the four characterized curcumin biosynthetic genes (labeled: Cl). Intriguingly, three Zo sequences clustered as a separate group from the three CURS clades.

**Figure 2:**
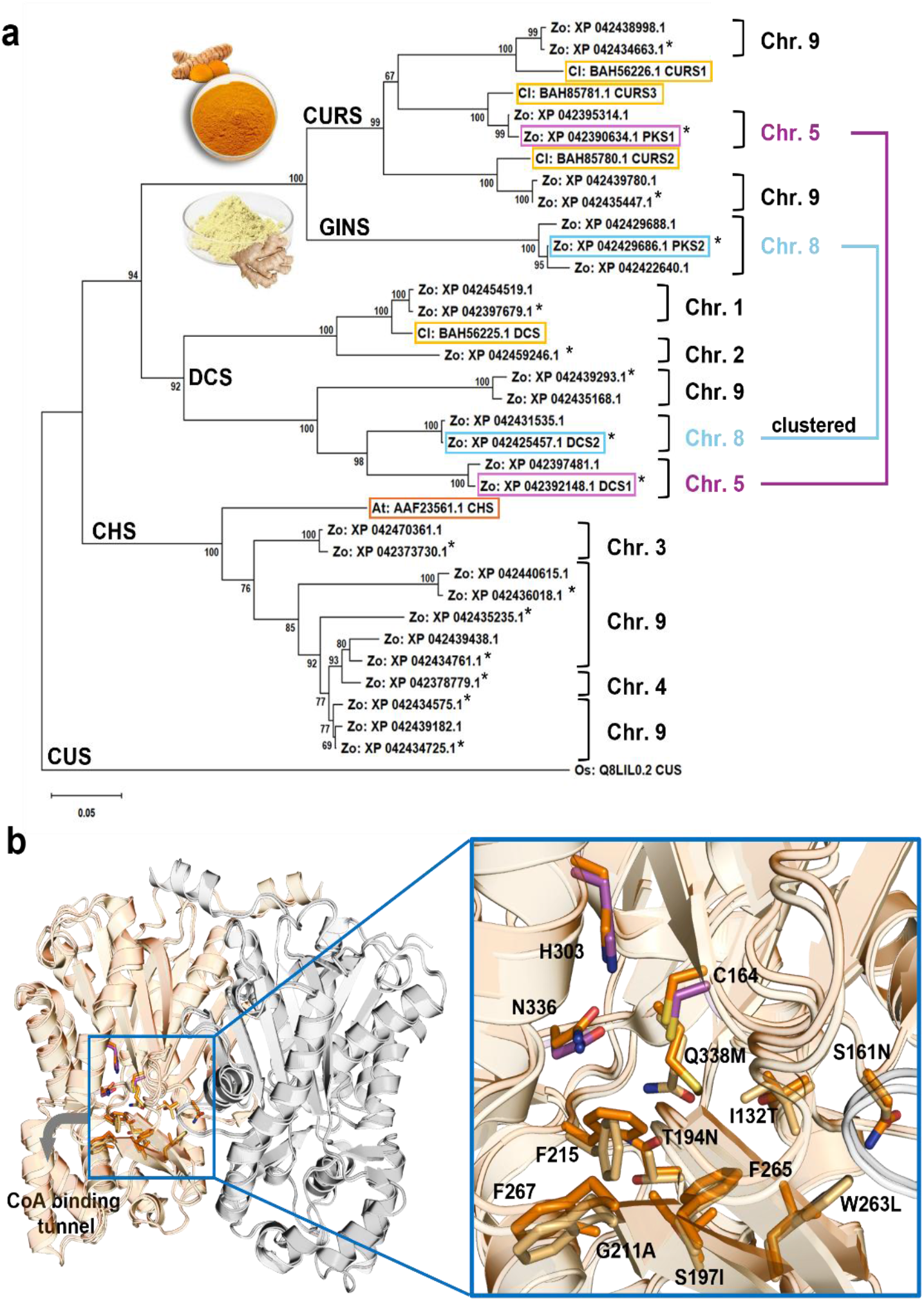
Identification of 6-DHG biosynthetic candidate genes. **(a)** Phylogenetic analysis of putative PKS and DCS genes from *Zingiber officinalis* (*Zo*) identified in the original NCBI Bioproject genome assembly (PRJNA 647255, 16 sequences labeled with a *) and in NCBI’s automatically generated RefSeq (PRJNA 73695, 13 additional sequences). Protein sequences were aligned with known curcumin biosynthetic proteins (CURS1-3, DCS) from *Curcuma longa* (Cl) and CUS from *Oryza sativa* (Os), and with chalcone synthase CHS from *Arabidopsis thaliana.* NCBI protein sequences accession numbers (**Table S1**) and chromosome locations are shown. Evolutionary history was inferred by the Neighbor-Joining method with a bootstrap test (500 replicates) (see Methods). Clustered *Zo*PKS1/DCS1 (Chr. 5) and *Zo*PKS2/DCS2 (Chr. 9) selected for characterization are highlighted. *Zo*PKS2 was identified as 6-DHG synthase (GINS). Images depict orange turmeric and yellow ginger spices extracted from their respective plant rhizomes. **(b)** Alignment of a *Zo*PKS2 homology model (dark orange) with the structure of CURS1 (PDB 3OV2) (light orange), which was used as template (see Methods). Shown are the homodimers with one monomer highlighted. Arrow depicts location of the CoA-binding tunnel for the extender substrate. Expansion shows residues lining the active site pocket, including the catalytic triad (purple, CURS1 residues) and the gatekeeper F residues (F215, F267, F265). Residues mutated in *Zo*PKS2 are indicated (e.g. W263L). See Supporting **Figure S1** for annotated protein sequence alignment and alignment of homology models of both *Zo*PKS1 and *Zo*PKS2 with the CURS1 structure.

To narrow down the number of putative 6-DHG candidate genes for functional characterization, we looked closely at their chromosomal locations, knowing that many plant natural products biosynthetic genes are organized in clusters ^27–28^, which could also apply to the co-location of the presumed 6-DHG DCS/PKS pair.

Indeed, two Zo CURS homologs (including their alleles) located on Chr 8 (Chr 8B: XP_042429686.1 (PKS2), XP_042429688.1, Chr 8A: XP_042422640.1) and Chr 5 (Chr 5A: XP_042390634.1(PKS1), Chr 5B: XP_042395314.1) were clustered with a DCS on Chr 8 (Chr 8A: XP_042425457.1 (DCS2), Chr 8B: XP_042431535.1) and Chr 5 (Chr 5A: XP_042392148.1 (DCS1), Chr 5B: XP_042397481.1) (**Figure 2a**). Notably, this included the three alleles in the separate CURS subclade. None of the other Zo CURS homologs on Chr 9 nor the Zo DCS homologs on Chr 1, 2 and 9 appeared to be part of a DCS/PKS cluster. From this analysis, we hypothesized that the gingeroid candidate genes are most likely located in one of the DCS/PKS pairs on Chr 5 and 8 and we selected one DCS and PKS allele from each cluster (Chr5: *Zo*DCS1/*Zo*PKS1; Chr8: *Zo*DCS2/*Zo*PKS2) for characterization as marked in **Figure 2a**. Curiously, we also noticed that in both DCS/PKS clusters, genes adjacent to the DCS are annotated as a chloroplastic, bifunctional diterpene synthase (predicted levopimaradiene synthase) which has no known role in gingeroid biosynthesis. However, both DCS/PKS pairs are predicted to be localized in the cytoplasm ^29–30^ where phenylpropanoid precursors like feruloyl-CoA are biosynthesized ^31–32^. Short-chain fatty acids for gingeroid biosynthesis are presumably in the plastids and CoA-activated in the cytoplasm as shown for cannabinoid biosynthesis ^33^.

Finally, to further narrow down the potential 6-DHG synthase candidates, we aligned the two selected *Zo*PKS protein sequences and their respective alleles with the three CURS proteins (**Figure S1a**). We also created homology models of *Zo*PKS1 and 2 based on the available CURS1 structure (PDB #3OV2) (**Figure 2b**)^11^. All sequences shared greater than 70% sequence identity, with each other and alleles being almost identical except for 1-2 residue differences. *Zo*PKS2 is the most divergent sequence (81.5% ZoPKS1 vs. 71.6 % ZoPKS2 sequence identity with CURS1), with changes in several key residues in the active site cavity (**Figure S1a**). Comparison of the *Zo*PKS2 active site model with that of CURS1 (**Figure 2b**), indicates that while the configuration of the catalytic triad (Cys164, His303, Asn336, using CURS1 numbering) involved in tethering the starter substrate (feruloyl- or hexanoyl-CoA) to Cys164 is preserved, several residues lining the active site pocket that are important for substrate binding and decarboxylative coupling of the diketide-CoA extender are different ^9, 34^ (**Figure 2b**). Notably, one of the “gatekeeper” phenylalanine residues (F215) that line a pocket of the CoA binding tunnel responsible for loading the feruloyldiketide substrate is rotated in the *Zo*PKS2 model. In contrast, the active site configuration of the ZoPKS1 model is almost identical to that of CURS1 (**Figure S1b**). Together these sequence and active site differences suggested that *Zo*PKS2 may have a different substrate profile and be able to accommodate longer acyl-CoA chains as starter substrates.

### Modular plasmid design for screening of *Zo*DCS/PKS function in *E. coli*

To test our hypothesized 6-DHG biosynthetic pathway, we designed a dual plasmid system for the combinatorial analysis of *Zo*PKS and *Zo*DCS functions together and individually in *E. coli* fed with different precursor substrates (**Figure 3**). For *in vivo* feruloyl-CoA synthesis, we chose 4-Coumaroyl-CoA Ligase 1 (4CL1) from *A. thaliana* which we have used previously used ^6–7^ and is known to accept diverse phenylpropanoic acids besides ferulic acid for subsequent substrate profiling experiments ^35^. For the *in vivo* synthesis of the hexanoyl-CoA substrate, we chose Acetyl-CoA Synthetase (ACSmut) from *A. thaliana* that has been engineered to accept hexanoic acid and other carboxylic acids as substrates ^36^.

**Figure 3:**
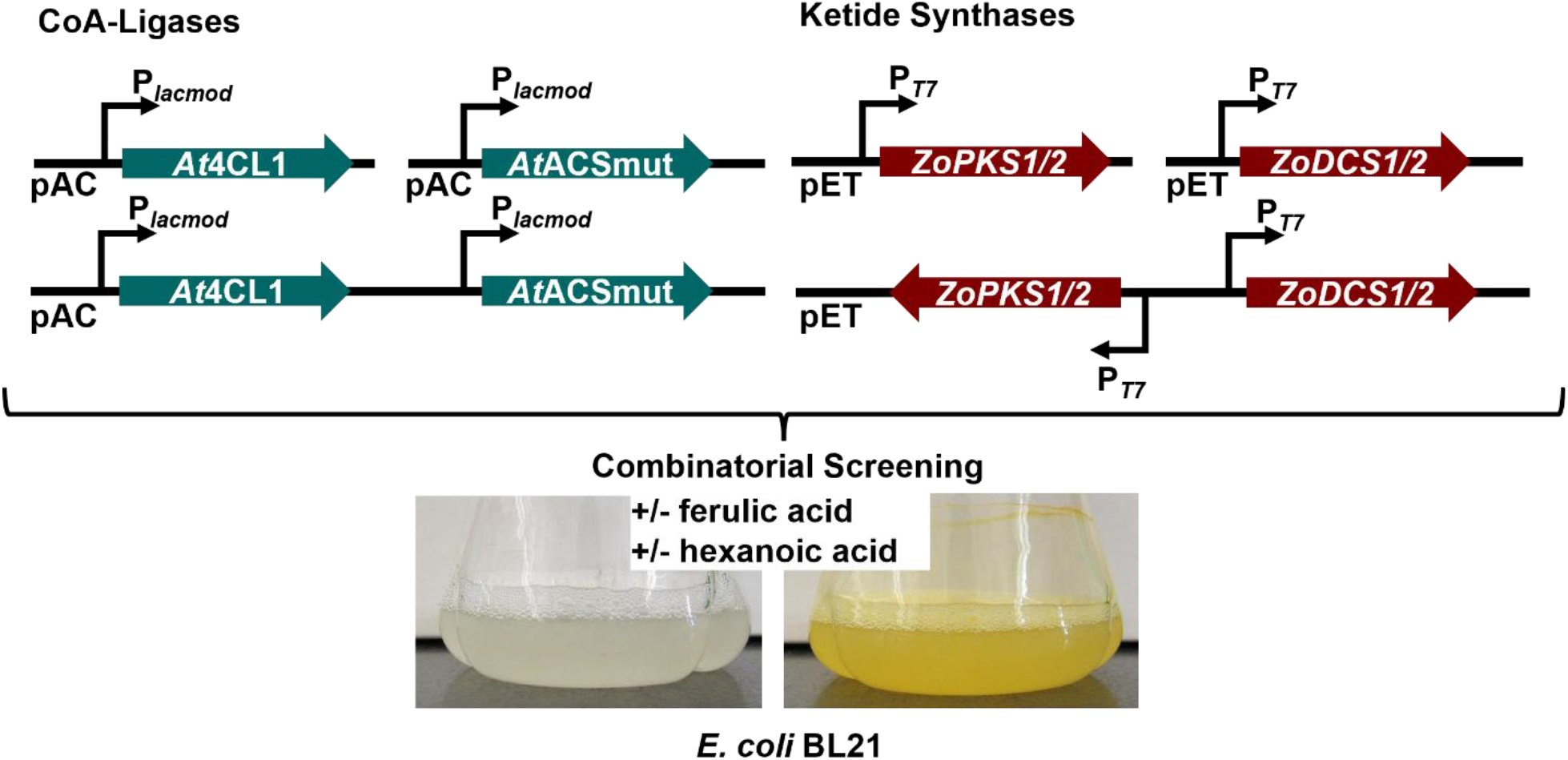
Modular plasmid design for combinatorial screening of 6-DHG biosynthesis. CoA-ligases (*At*4CL1, *At*ACSmut) for CoA-substrate biosynthesis were constitutively expressed (P_lacmod_) individually or together from a pAC plasmid. The four polyketide (PKS) and diketide (DCS) candidate genes were cloned individually or together in all four possible combinations for expression under the control of an inducible T7 promoter on a pET plasmid as shown. *E. coli* BL21 cells harboring different plasmid combinations, including empty plasmid controls, were induced and supplemented with or without ferulic and/or hexanoic acid substrate precursors. Functional PKS/DCS pairs are expected to yield yellow products for compounds with a λ_max_ > 400 nm like curcumin (λ_max_ = 420 nm) or uncolored or pale-yellow products for e.g. 6-dehydrogingerdione (6-DHG) (λ_max_ = 370 nm & shoulder absorption at > 400 nm).

We first attempted to adapt a multi-plasmid strategy that we previously used for stilbene biosynthesis ^6^ by placing the expression of the CoA-ligases and the ketide synthases under the control of a constitutive Lac_mod_ promoter and the use two compatible plasmid backbones for enzyme co-expression ^6–7, 37^. However, reproducibility issues in maintaining stable transformants required us to place ketide synthase expression under the control of an inducible T7 promoter and add a T7-terminator downstream of the genes to mitigate strain stability issues presumably due to product toxicity and/or resource depletion by recombinant gene overexpression in *E. coli*. Recombinant curcumin production in *E. coli* also reported the use of an inducible system ^38^. An N-terminal His-Tag was added to all recombinant proteins to aid expression of these plant proteins in *E. coli* and for future purification. Except for the previously cloned *At*4CL1 ^6^, all plant genes were codon-optimized for expression in *E. coli,* resulting in a final set of 13 plasmids for the co-expression of different combinations of CoA-ligases (pAC backbone, 4 plasmids) and DCS/PKS synthases (pET backbone, 9 plasmids) (**Table S3**). Conversion of exogenously fed hexanoic and ferulic acid substrates by the engineered *E. coli* cells expressing a functional PKS/DCS pair is then expected to yield products with distinct UV/Vis spectra due to their feruloyl moiety (**Figure 3**).

### Identification of *Zo*PKS2 as a 6-DHG synthase and *Zo*PKS1 as a curcumin synthase

For initial screening of *Zo*PKS function, we co-expressed in *E. coli* the different combinations of *Zo*PKS and *Zo*DCS enzymes with the two CoA-Ligases (*At*4CL1 and *At*ACSmut) and compared product formation to empty vectors (EVs) or CoA-ligases only transformants. After induction of *Zo*PKS/DCS expression and media supplementation with the two precursor substrates, we observed the formation of new products only in cultures expressing complete pathways consisting of the *Zo*PKS/DCS combinations and the two CoA-ligases, suggesting that their biosynthesis is dependent on CoA-activated precursors (**Figure 4, Figure S2**). SDS-PAGE analysis of protein expression at the end of the cultivation experiments confirmed overexpression of all recombinant proteins, with strong expression levels obtained for the ketide synthase genes under the control of the T7 promoter (**Figure S3a**).

**Figure 4:**
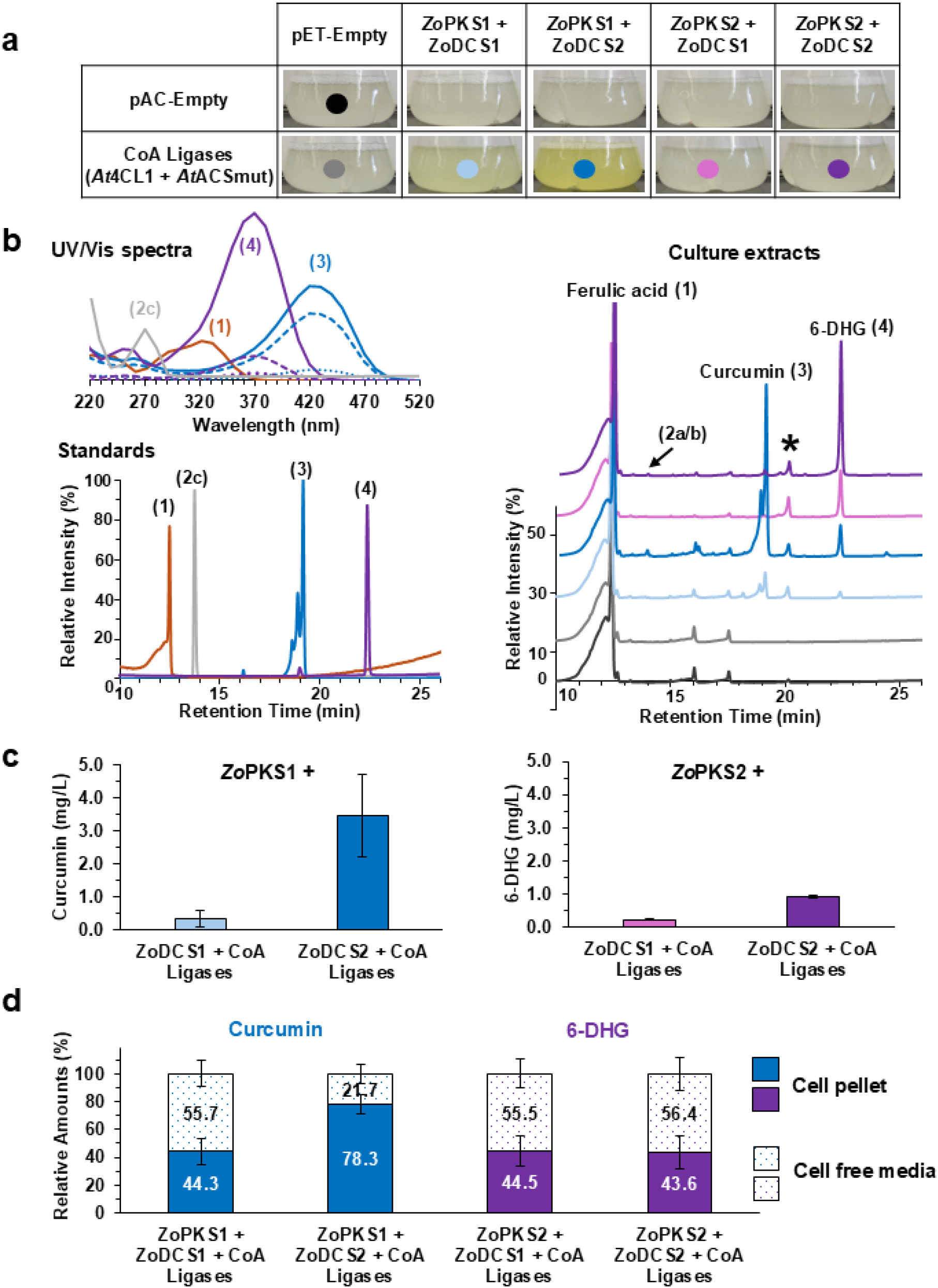
Conversion of ferulic and hexanoic acid by engineered *E. coli* co-expressing *Zo*PKS/DCS and CoA-ligases. **(a)** Recombinant *E. coli* BL21 cultures^#^ co-expressing Co-ligases (*At*4CL1 & *At*ACSmut) with *Zo*PKS1 and DCS1/2 accumulate a yellow product after feeding with ferulic and hexanoic acid. **(b)** HPLC analysis of total *E. coli* culture extracts monitored at 370 nm with additional chromatograms of controls shown in **Figure S2**. Analytical standards of ferulic acid (1), curcumin (3), 6-DHG (4) and zingerone (2c*, monitored at 290 nm) for comparison were run under identical conditions to identify compounds peaks based on retention times and UV/Vis spectra shown where solid lines represent standards, long and short dashed lines represent peaks from *Zo*PKS2 and *Zo*PKS1 cultures, respectively (see also **Figure S4**). Preparative HPLC followed by LC-MS/MS of fractionated peaks (**Figure S5**) was performed to further confirm compound identities by their mass spectra (**Table S2**). HPLC chromatograms are stacked for clarity and are scaled to the same relative intensity range of the bottom profile, with each profile’s 100% relative intensity set by its maximum absorbance peak. (**c**) Concentrations of curcumin (*Zo*PKS1) or 6-DHG (*Zo*PKS2) quantified from extracts of total *E. coli* cultures co-expressing *Zo*PKS1/2 in combination with *Zo*DCS1/2 and CoA-ligases, and (**d)** relative ratios of curcumin (*Zo*PKS1) and 6-DHG (*Zo*PKS2) extracted from cell pellets or cell-free culture supernatants from these cultures. ^#^All *E. coli* cultures were grown in modified M9 medium with 5 g L^-1^ glycerol at 25 °C and fed with 1 mM CoA-precursor substrates after 3 h of IPTG induction of *Zo*PKS/DCS expression. After 20 h of conversion, cultures were processed for analysis. Data were collected from three cultures and data points are the mean of three independent biological replicates. Representative HPLC profiles are shown. Compound/peak labels follow numbering in Figure 1. Color schemes for HPLC profiles correspond to those in panel **(a)**. Purple and blue colors represent either 6-DHG and curcumin products or *Zo*PKS2 and *Zo*PKS1 products, respectively.

*E. coli* cultures co-expressing *Zo*PKS1 with either *Zo*DCS1 or *Zo*DCS2 accumulated yellow products with the strongest color produced by the *Zo*DCS2 expressing strain (**Figure 4a**). HPLC separation of total culture extracts from the *Zo*PKS1 cultures (blue labels) with either of the *Zo*DCSs yielded three unique peaks in addition to unconverted ferulic acid (1). The UV/Vis spectrum (λ_max_ = 420 nm) and retention time of the major peak (3) matched that of the curcumin standard, which also had multiple peaks due to keto-enol tautomerization ^39^ (**Figure 4b**). The smaller peak (4) had a λ_max_ = 370 nm and retention time that matched the 6-DHG (4) standard.

A unique minor peak (2a/2b) eluted shortly after ferulic acid (1) and became significantly more prominent in subsequent *vivo* substrate profiling experiments (**Figure 5 & S6**) and will be discussed below. Other smaller peaks, including one peak with a retention time ∼20 mins, were also present in control cultures (empty vectors, CoA-Ligases only (**Figure 4b & S2**)) and were therefore not considered to be directly associated with the activities of the *Zo*PKS/DCS enzymes expressed in our recombinant *E. coli* strains. The emergence of a smaller peak (*) that eluted after curcumin (3) was strongly correlated with the expression of *At*ACSmut and feeding of hexanoic acid (see **Figure 5 & S6** below) and appeared to be a stress response metabolite that appeared to be a peptide because of its UV/Vis spectrum (λ_max_ = 270 nm) and high molecular weight (*m/z*: 931.13) (**Table S2 & Figure S5**).

**Figure 5:**
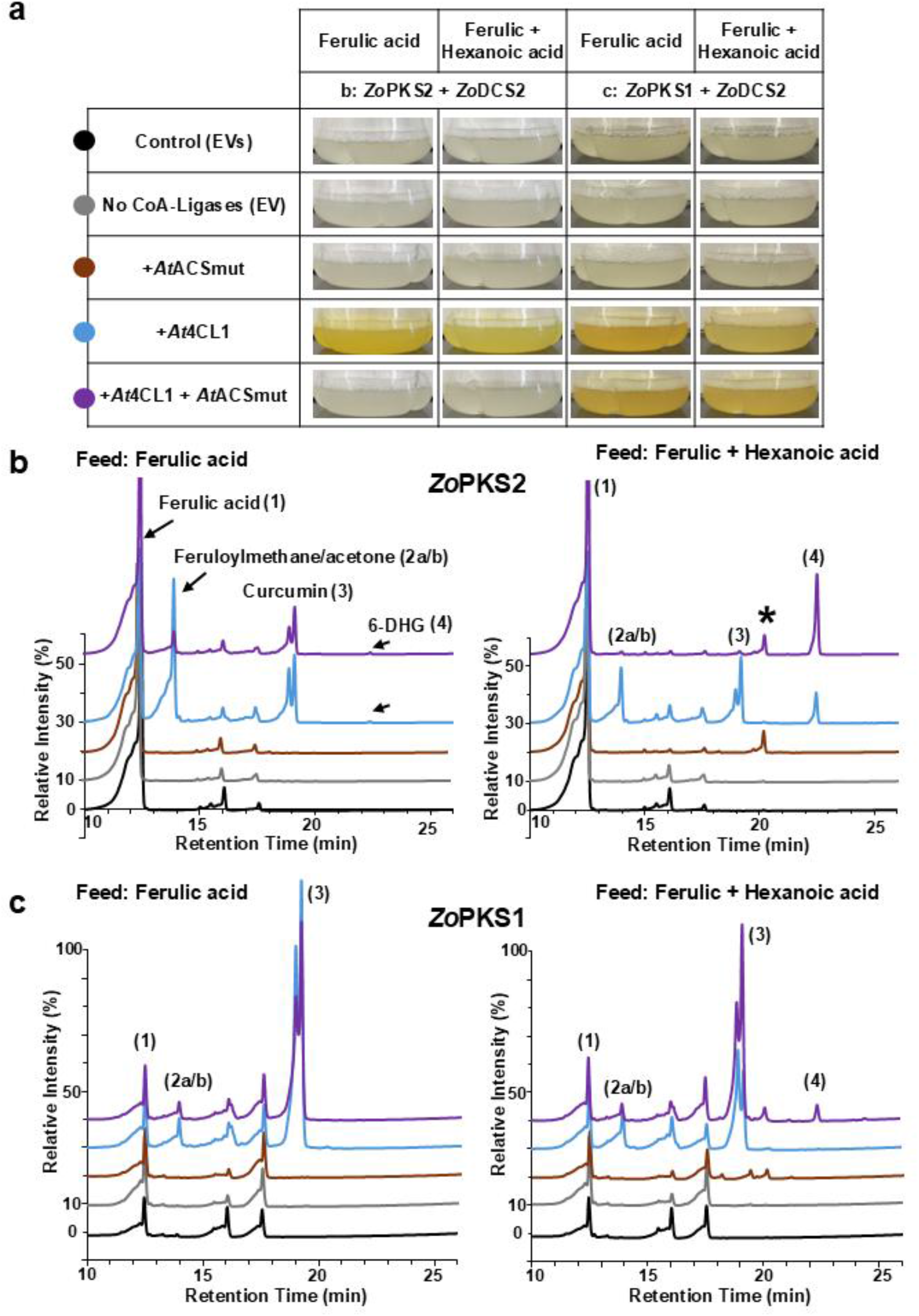
Profiling of starter substrate preference of engineered *E. coli* co-expressing *Zo*PKS/DCS and different CoA-ligase combinations. **(a)** Recombinant *E. coli* BL21^#^ cultures co-expressing *Zo*PKS1 or ZoPKS2 with *Zo*DCS2 and *At*4CL1 accumulated a yellow product after feeding with ferulic acid alone or both ferulic and hexanoic acid. Co-expression of both *At*4CL1 and *At*ACSmut with either *Zo*PKS1 or *Zo*PKS2 and *Zo*DCS2 only yielded a yellow product for ZoPKS1. No Feed and hexanoic acid only control cultures did not accumulate yellow products (**Figure S6**). (**b, c**) HPLC analysis of cell-free culture extracts monitored at 370 nm are shown for *Zo*PKS2 (**b**) and *Zo*PKS1 (**c**) cultures co-expressing the gene combinations shown in panel a. Colors of HPLC chromatograms correspond to different CoA ligase combinations and empty vectors (EVs) controls. Cultures were either fed only ferulic acid or ferulic and hexanoic acid. Additional chromatograms of “no feed” and hexanoic acid only fed control cultures are shown in **Figure S6**. HPLC chromatograms are stacked for clarity and are scaled to the same relative intensity range of the bottom profile as in Figure 4. ^#^All *E. coli* cultures were grown in modified M9 medium with 5 g L^-1^ glycerol at 25 °C and fed with 1 mM CoA-precursor substrates after 3 h of IPTG induction of *Zo*PKS/DCS expression. Controls consisted of empty vector strains (EVs) transformed with empty pAC and pET28a and No CoA Ligases (EV) strains were transformed with pET28a*-Zo*PKS/DCS and empty pAC. After 20 h of conversion, cultures were processed for analysis. Extracts and data were obtained from two independent cultures for each experimental condition. One HPLC profile for each condition is shown. Compound/peak labels follow numbering in Figure 1. Compound identities were confirmed with authentic standards, by UV/Vis and LC-MS analysis (Figure 5S, **Table S2**). Color schemes for HPLC profiles correspond to those in panel (**a**).

Unlike the *Zo*PKS1 expressing strains, cultures co-expressing *Zo*PKS2 with *Zo*DCS1 or 2 (magenta/violet labels) did not turn yellow (**Figure 4a**). HPLC analysis of their total culture extracts showed one major, unique peak (4) with a retention time and UV/Vis spectrum (λ_max_ = 370 nm) that matched those of the 6-DHG standard (**Figure 4b**). Interestingly, *Zo*PKS2 cultures also produced a small amount of curcumin (3) detected as a small peak in their chromatograms. LC-MS analysis of the fractionated peaks 3 and 4 from the *Zo*PKS1 and *Zo*PKS2 cultures and comparison to the MS spectra of the authentic standards confirmed them as curcumin (*m/z*: 367.12) and 6-DHG (*m/z*: 289.14) (**Table S2**).

Our initial cultivation experiments therefore suggested that *Zo*PKS2 prefers *in vivo* hexanoyl-CoA as a starter substrate to produce 6-DHG while *Zo*PKS1 like CURS1 favors feruloyl-CoA for curcumin synthesis but also synthesizes less efficiently 6-DHG. These results supported our initial hypothesis that *Zo*PKS2 has a different substrate profile compared to *Zo*PKS1 and the characterized CURS enzymes ^10, 12^ due to changes in its active site (**Figure 2**).

Additionally, we found that both *Zo*PKS’s function in *E. coli* with either of the *Zo*DCS enzymes regardless of whether they are clustered together in the genome. However, cultures with combinations of *Zo*PKS1/2 with *Zo*DCS1 produced significantly less polyketide products than those with *Zo*DCS2 (**Figure 4c**). Quantification of products from total culture extracts gave curcumin yields for *Zo*PKS1 cultures of 3.46 mg L^-1^ with *Zo*DCS2 and a 10-fold lower (0.34 mg L^-1^) with *Zo*DCS1. Similarly, 6-DHG production level for *Zo*PKS2 cultures were 0.94 mg L^-1^ with *Zo*DCS1 and 4-fold lower (0.23 mg L^-1^) with *Zo*DCS2 (**Figure 4c**). The reduced yields with *Zo*DCS1 may be explained by lower expression levels or solubility in *E. coli* compared to *Zo*DCS2. Indeed, SDS-PAGE analysis confirmed that *Zo*DCS1 is much less soluble than *Zo*DCS2 (**Figure S3c**). Because of the higher activity of *Zo*DCS2 in *E. coli*, we therefore used this enzyme in all subsequent experiments.

Considering that curcumin and 6-DHG differ in size and polarity which may impact their abilities to cross the *E. coli* cell membranes ^40^, we also analyzed their partitioning between cell pellets and cell-free culture supernatants in the higher yielding *Zo*DCS2 containing cultures to guide subsequent product analysis and quantifications. As expected, the less polar and smaller 6-DHG partitioned readily out of the cell with nearly equal quantities in both the media and cells, while a larger fraction of the curcumin remained in the cell pellet at the end of the 20 h conversion phase (**Figure 4d**). Nevertheless, both compounds were present in significant quantities in cell-free culture supernatants which were used for subsequent extractions to reduce the number of co-extracted, unspecific metabolites.

### *Zo*PKS2 efficiently uses feruloyl-CoA as starter when hexanoyl-CoA is absent

During our initial screening experiments, we observed that *Zo*PKS2 also produced a small amount of curcumin in addition to its major product 6-DHG, indicating that the enzyme prefers hexanoyl-CoA as starter substrate but also accepts feruloyl-CoA when both substrates are co-fed (**Figure 4b**). To explore *Zo*PKS2’s starter substrate promiscuity further as a potential strategy to use it as a dual-function enzyme in *E. coli* for the production of either of these two valuable natural products simply by providing access to different starter substrates, we changed hexanoyl-CoA availability via feeding or the absence of hexanoyl-CoA ligase *At*ACSmut. For comparison, we also conducted the same experiments with *Zo*PKS1 transformed with the same combinations of CoA ligases (*At*ACSmut, *At*4CL1) or none (**Figure 5 & S6** for additional controls). Cultures were fed ferulic acid, hexanoic acid, both or neither as in **Figure 4**.

Just by comparing the colors produced by the *E. coli* cultures after 20 h of conversion it was apparent that *Zo*PKS2/*Zo*DCS2 made the yellow curcumin only when provided with ferulic acid alone and expressing the *At*4CL1 CoA-ligase (**Figure 5a**). Under these conditions, co-expression of the hexanoyl-CoA ligase (*At*ACSmut) without hexanoic acid also led to the accumulation of the yellow curcumin. This matched the color profiles of the corresponding *Zo*PKS1/*Zo*DCS2 cultures, with one striking difference that indicated that *Zo*PKS1 is a curcumin synthase while *Zo*PKS2 is a 6-DHG synthase; when both ferulic and hexanoic acid are supplied and the strains express both CoA-ligases, *Zo*PKS2 cultures appeared to shift production to the colorless 6-DHG while *Zo*PKS1 cultures continued the synthesis of the yellow curcumin.

HPLC-analysis of the products at the end of the conversion phase confirmed that *Zo*PKS2 shifts from curcumin (3) to 6-DHG (4) production when hexanoyl- and feruloyl-CoA are both available via feeding and CoA-ligase expression (**Figure 5b**). This suggests that *Zo*PKS2 strongly prefers hexanoyl-CoA as starter substrate over feruloyl-CoA for condensation with the feruloyldiketide CoA provided by *Zo*DCS2. As expected, no products were detected when feruloyl-CoA was not available in the no-feed and hexanoic acid only control cultures (**Figure S6**) or the No-CoA-ligase and *At*ACSmut-only strains. Interestingly, we detected a small peak of 6-DHG (4) in *Zo*PKS2 cultures without hexanoic acid supplementation or *At*ACSmut expression, suggesting that the enzyme used endogenous hexanoyl-CoA from *E. coli* fatty acid metabolism. Intriguingly, the relative amount of this 6-DHG peak compared to curcumin increased when ferulic and hexanoic acid were fed but only *At*4CL1 was expressed for feruloyl-CoA synthesis, which suggested that under these conditions *E. coli* accumulated higher levels of hexanoyl-CoA from the supplemented hexanoic acid for *Zo*PKS2 to use for 6-DHG synthesis.

In contrast to *Zo*PKS2 and observed in our initial screening (**Figure 4**), *Zo*PKS1 strongly favored ferulic-CoA as a starter for the synthesis of curcumin (3) and made only a minor amount of 6-DHG (4) when hexanoyl-CoA is also available (**Figure 5c**). Like *Zo*PKS2, no PKS derived products were made by control culture that were unable to synthesize feruloyl-CoA. These results confirmed that *Zo*PKS2 is a *bona fide* dehydrogingerdione synthase while *Zo*PKS1 is a curcumin synthase. However, if only provided with feruloyl-CoA, *Zo*PKS2 will switch to curcumin synthesis.

In addition to the expected curcumin (3) and 6-DHG (4) products, we also observed the presence of a peak in both *Zo*PKS2 and *Zo*PKS1 cultures that eluted shortly after ferulic acid with an UV/Vis maximum of 330-340 nm (*m/z*: 191.07) matching that of feruloylmethane/dehydrozingerone (2a), a known curcumin degradation product ^41^. This peak appeared under conditions when *Zo*PKS1 and *Zo*PKS2 produced curcumin and particularly, in *Zo*PKS2 cultures that were fed ferulic acid only and expressed only *At*4CL1 (**Figure 5b**). The broad shoulder of this peak suggested the presence of additional curcumin degradation products such as feruloylacetone (2b) ^42^ for which a corresponding parent ion (*m/z*: 233.12) and base peak (*m/z*: 173.08) was detected by negative-ion MS (**Figure S5**). As in **Figure 4**, we also observed a peak at ∼20 min retention time in cultures overexpressing *At*ACSmut and fed with hexanoic acid, including in the hexanoic acid control but not in the No-feed cultures (**Figure S6**), indicating that this compound was unrelated to *Zo*PKS/DCS enzyme function and may be a stress response molecule.

### *Zo*PKS2 makes 6-DHG derivatives in engineered *E. col*i fed with coumaric or cinnamic acid

Type III PKS are known to exhibit some degree of substrate promiscuity, especially for their starter substrate ^5^. The three CURS enzymes from *C. longa* accept both feruloyl- and coumaroyl-CoA as substrates *in vitro* to produce with varying efficiencies bismethoxy- and dimethoxycurcumin in addition to curcumin ^10–12^, which reflects the mixtures isolated from turmeric rhizomes and also indicates that the *C. longa* DCS converts these substrates also into the diketide extender CoA substrates. The unusual CUS enzyme has been extensively investigated for the *in vivo* and *in vitro* synthesis of diverse curcuminoids ^14, 16, 24–25, 43^ because of its ability to produce a diketide-CoA intermediate, that upon hydrolysis into the β-keto acid can become the extender for another condensation reaction. CUS was also shown to catalyze *in vitro* the condensation of fatty acyl chains to phenylpropanoids when provided with an activated β-keto fatty acid substrate for the condensation reaction. For *in vivo* dehydrogingeroid biosynthesis by CUS though, *E. coli* had to be engineered with a β-oxidation pathway to provide the necessary β-keto acyl-CoA substrates ^24^.

Because *Zo*PKS2 paired with *Zo*DCS2 could be used in simplified microbial production platform for the precursor directed synthesis of natural and unnatural gingeroids from inexpensive carboxylic acids, we explored as a first proof-of-concept the enzyme’s promiscuity with coumaroyl- and cinnamoyl-CoA – substrates known to be accepted by the CURS enzymes ^10, 12^. Like the native DCS enzymes from turmeric, we expected that *Zo*DCS2 would accept these additional starter substrates to produce the corresponding diketide-CoA extender substrates. Indeed, upon feeding *E. coli* strains containing the complete *Zo*PKS2/DCS2/*At*4CL1/*At*ACSmut pathway with hexanoic acid and either of these two phenylpropanoic acids, both cultures produced new colored products (**Figure 6a**). As a comparison, we did the same feeding experiments with the completed *Zo*PKS1 pathway and did not observe the synthesis of new colored products besides the yellow curcumin product in the ferulic acid control culture.

**Figure 6:**
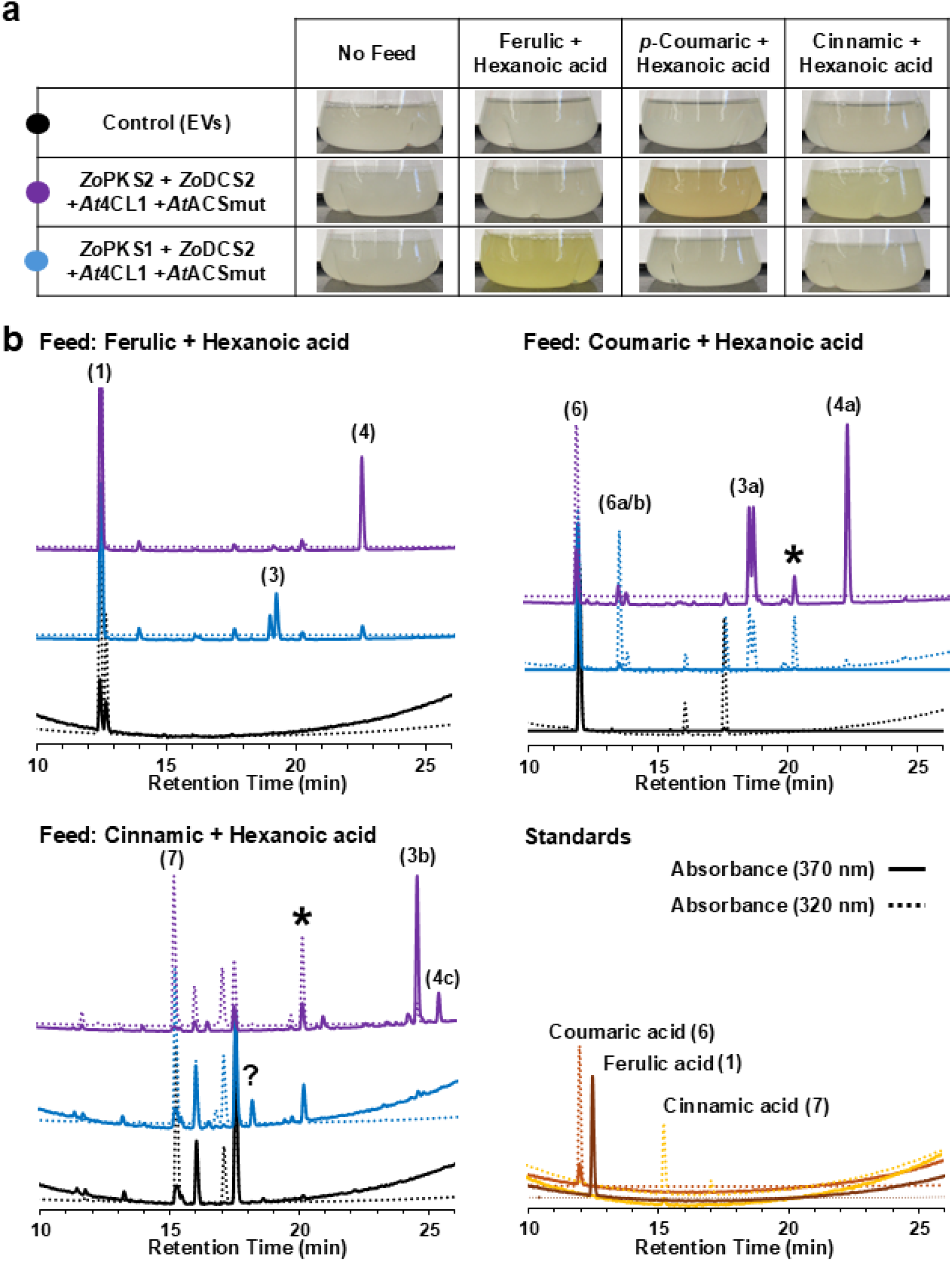
Exploring the conversion of *p*-coumaric acid and cinnamic acid into curcumin and 6-DHG derivatives. **(a)** Recombinant *E. coli* BL21 cultures^#^ co-expressing *Zo*PKS2 or *Zo*PKS1 with *Zo*DCS2, *At*4CL1 and *At*ACSmut were fed with phenylpropanoic acids (ferulic, *p*-coumaric, cinnamic acid) and hexanoic acid. The accumulation of yellow and orange products was visible in *Zo*PKS2 cultures fed with *p*-coumaric and cinnamic acid, while ZoPKS1 cultures produced, as observed before, a bright yellow product with ferulic acid. **(b)** HPLC analysis of products from cell free media extracts monitored at 370 nm (solid traces) and 320 nm (dotted traces, to better monitor cinnamic and coumaric acid derived compounds) are shown for the three feeding conditions. HPLC profile colors correspond to the *Zo*PKS2 and *Zo*PKS1 pathway expressing strains and empty vector (EV) control strain shown in panel a. HPLC traces are stacked for clarity, and importantly, each chromatogram is scaled to 60% of its highest (100%) intensity peak to visualize less abundant peaks. Peak label numbers correspond to compounds in Figure 1 and have been identified by a combination of UV/Vis (**Figure S4**), MS spectra (**Table S2**) and retention times (standards). Peaks labeled with (*) correspond to unspecific metabolites observed under previous conditions (Figure 4 **& 5**), including in control cultures. (?) denotes unidentified product peak. ^#^All *E. coli* cultures were grown in modified M9 medium with 5 g L^-1^ glycerol at 25 °C and fed with 1 mM phenylpropanoic and hexanoic acid substrates after 3 h of IPTG induction of PKS/DCS expression. After 20 h of conversion, cultures were processed for analysis. Extracts were obtained from three independent cultures for each experimental condition. One representative HPLC profile for each condition is shown.

HPLC analysis (**Figure 6b**) confirmed the production of new compounds with *p*-coumaric acid by *Zo*PKS2 as the gingeroid *p*-coumaroylhexanoylmethane (4a) (λ _max_ = 360 nm) and curcuminoid bisdemethoxycurcumin (3a) (λ _max_ = 418-420 nm) which were responsible for the color of the cultures. UV/Vis spectra and negative ion MS data (4a *m/z*: 259.13, 3b *m/z*: 307.10) (**Table S2 & Figure 5S**) for these products agree with those reported for CUS and CURS enzymes ^24–25^. In comparison, *Zo*PKS1 cultures produced very small amounts of the curcuminoid (3a) and an almost undetectable amount of the gingeroid (4a), which were only made visible upon scaling the relative intensity of the HPLC traces recorded at 320 nm. Both *Zo*PKS1 and *Zo*PKS2 also accumulate a peak (6a/b) that appears to be a degradation product (or a mixture) of either the *p*-coumaroyl diketide intermediate or bisdemethoxycurcumin (λ _max_ = 320 nm) like the feruloylmethane/acetone (2a/b) peak observed for the ferulic aid fed cultures in **Figure 5**. The negative ion MS data indicated the presence of *p*-coumaroyl-ß-keto-acid (6a) (*m/z*: 205.05) and *p*-hydroxybenzalacetone (*p*-coumaroylmethane) (6b) (*m/z*: 161.04) (**Table S2**). Feeding of cinnamic and hexanoic acid yielded two new compound peaks (3b, 4c) only with *Zo*PKS2 that were identified as dicinnamoylmethane (3b) (λ _max_ = 390 nm, *m/z:* 276.16) and a derivative of the gingeroid cinnamoylhexanoylmthane (4b) (labeled 4c). The red-shifted UV/Vis spectrum (λ _max_ = 390 nm) and negative ESI fragment (*m/z* = 227.20) (**Table S2 & Figure 5S**) of this peak suggest it to be phenyldeca-1,4-dien-3-one (4c) which has an extended conjugated systems due to the loss of the keto group of 4b. *Zo*PKS1 cultures did not seem to make any curcuminoid or gingeroid products with cinnamic acid. One new compound peak appeared with UV/Vis and MS properties (λ _max_ = 270 nm, *m/z*: 213.96) (**Table S2, Figure S5**) that did not match any expected enzyme products or derivatives thereof and is labeled as unknown (?) in **Figure 6**.

These preliminary feeding studies showed that *Zo*PKS2 does accept alternative phenylpropanoic acid substrates to produce gingeroids but appeared to be less efficient in catalyzing the condensation with hexanoyl-CoA as cultures accumulated much larger ratios of curcuminoids compared to the ferulic acid fed ones which almost exclusively produced 6-DHG (**Figure 6b**).

### Identification of bottlenecks and opportunities for future metabolic engineering

While the objective of this study was primarily the identification of a 6-DHG PKS to enable the recombinant production of gingeroids, we were surprised about the comparatively low polyketide yields considering that in previous studies we could readily reach 25-100 mg L^-1^ of flavonoid production in *E. coli* using the same media conditions and without significant optimization ^6-7^. Suspecting that hexanoic acid and expression of hexanoyl-CoA ligase (*At*ACSmut) negatively impacted *E. coli* cells, and thus 6-DHG production levels, we decided to quantify polyketide production in the presence and absence of *At*ACSmut and/or hexanoic acid feeding for *Zo*PKS2/DCS2 and *Zo*PKS1/DCS2 strains that also expressed *At*4CL1 (**Figure 7a**). Without the expression of *At*ACSmut and feeding of only ferulic acid, *Zo*PKS2 strains produced 14.5 mg L^-1^ curcumin compared to 3 mg L^-1^ made by *Zo*PKS1 cultures. This showed that *Zo*PKS2 under these conditions is a more efficient curcumin synthase than *Zo*PKS1 in *E. coli*, which may be due to higher soluble expression of *Zo*PKS2 (**Figure S3**).

**Figure 7:**
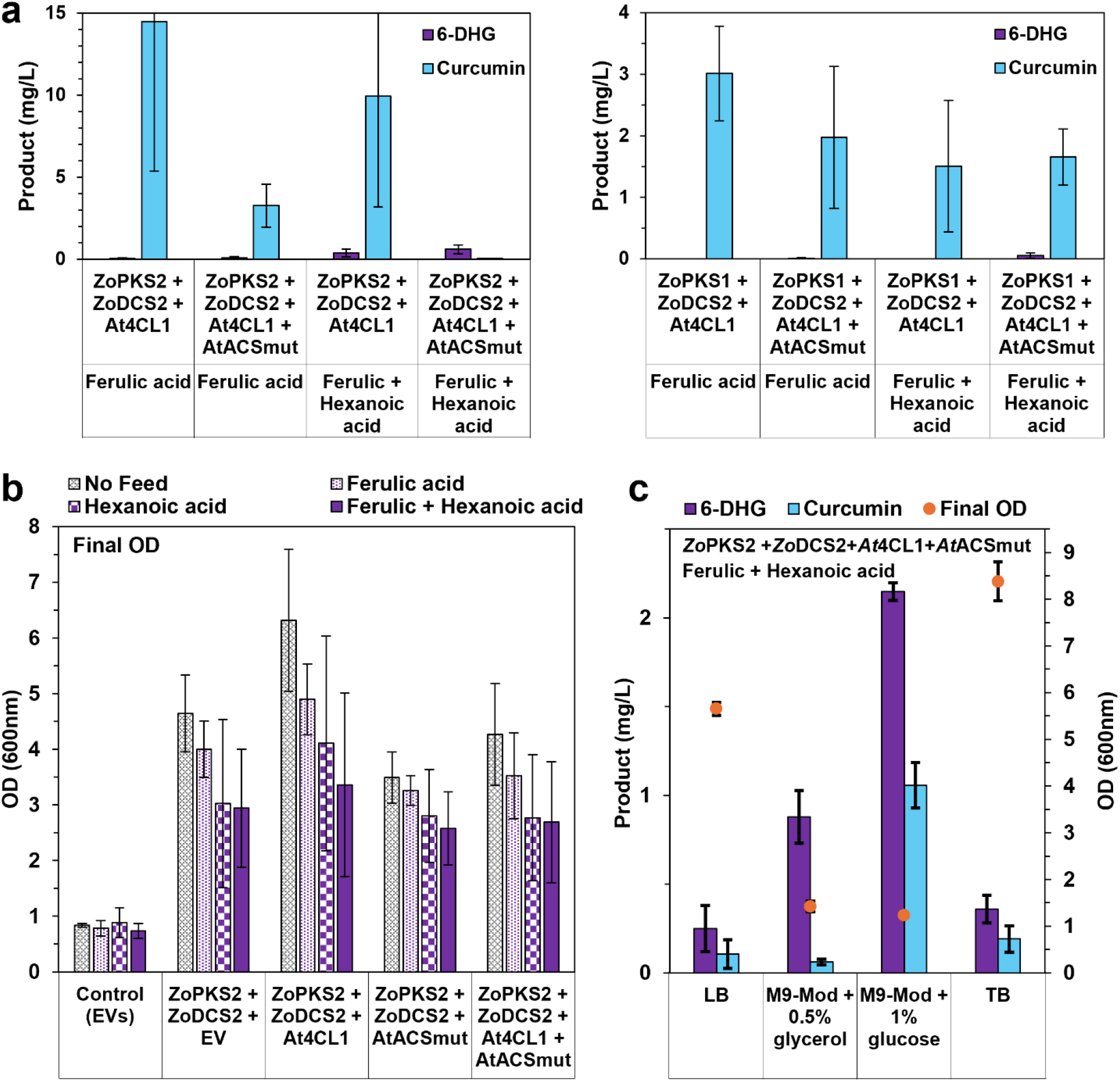
Comparing curcumin and 6-DHG production and cell growth across recombinant *E. coli* strains and media conditions. **(a)** Curcumin and 6-DHG yields are compared for *Zo*PKS2 and *Zo*PKS1 *E. coli* strains expressing different combinations of CoA ligases and fed either ferulic acid alone or both ferulic and hexanoic acid. Products are quantified from cell free media extracts from three cultures and data are shown as mean values ± SD and error bars represent the standard deviations of three, independent biological replicates. **(b)** Final optical densities (OD_600_) reached of recombinant cultures (from Figure 5a) expressing different *Zo*PKS2/DCS2 and CoA ligases combinations or empty vectors (EVs) and fed with different substrates or none (see **Figure S7** for comparison with *Zo*PKS1/DCS1 data). Data show the average of two replicated cultures per condition and the error-bars show the variability between the two cultures. **(c)** Curcumin and 6-DHG product yields and final OD reached by *E. coli* strains expressing the complete *Zo*PKS2 pathway and grown in different media supplemented with 1 mM ferulic and hexanoic acid for 20 h of conversion in Figures 4 **& 5**. Products are quantified from cell free media extracts from three cultures and data are shown as mean values ± SD and error bars represent the standard deviations of three, independent biological replicates.

Remarkable though was the almost complete switch of *Zo*PKS2 strains to 6-DHG biosynthesis when supplied with hexanoyl-CoA via hexanoic acid feeding and *At*ACSmut expression, reaching levels that were comparable to curcumin yields obtained with *Zo*PKS1 (**Figure 7a**). These data again confirmed the results in **Figure 4 & 5** and demonstrated the strong affinity of *Zo*PKS2 for hexanoyl-CoA vs feruloyl-CoA as a starter – even when provided even with small quantities via endogenous hexanoyl-CoA pools boosted in *E. coli* by only hexanoic acid feeding and no *At*ASCmut expression or only *At*ASCmut expression and no feeding. Future *in vitro* experiments with purified *Zo*PKS enzymes and comparison with kinetic parameters of the CURS enzymes ^12^ are expected to identify mechanistic reasons for this selectivity and guide protein design studies.

During our production experiments, we noticed though that hexanoic acid feeding and/or *At*ACSmut expression reduced polyketide production levels not only in *Zo*PKS2 but also in *Zo*PKS1 strains by adversely impacting *E. coli* cell growth. As shown in **Figure 7b** & **S7**, hexanoic acid significantly impaired growth regardless of whether it is used as a substrate, including by *Zo*PKS1 strains. Further, co-feeding of ferulic acid and/or expression of *At*ASCmut had compounding effects on cell growth, and consequently polyketide yields. This is not surprising as short chain fatty acids and changes in acyl-CoA pools in *E. coli* are well known to be inhibitory for cell growth and are targeted for optimization in microbial production processes ^44–46^. For unknown reasons, our empty vector (EVs) control cultures (harboring empty and sequence verified pAC_mod_ and pET28a plasmids) consistently grew to much lower ODs compared to the recombinant *Zo*PKS strains, regardless of feeding conditions (**Figure 7b & S7**).

In a final attempt to increase the yield of 6-DHG production by *E. coli* cells expressing the full *Zo*PKS2 pathway, we compared hexanoic and ferulic acid conversion in different media, including Luria Bertani (LB), Terrific Broth (TB) and our modified M9 medium supplemented with glycerol (5 g L^-1^) (used in all previous experiments) or glucose (10 g L^-1^) (**Figure 7c**). Strains cultivated in the two nutrient rich media (LB and TB) produced the lowest 6-DHG yields of less than 0.5 mg L^-1^ (LB: 0.25 ± 0.13 mg L^-1^, TB: 0.36 ± 0.08 mg L^-1^) despite growing to the highest cell densities.

In contrast, the modified M9 media grown strains produced ∼1-2 mg L^-1^ of 6-DHG (M9-mod + glycerol: 0.88 ± 0.15 mg L^-1^, M9-mod + glucose: 2.15 ± 0.05 mg L^-1^) at much lower cell densities. Notable was the effect of media composition on the relative ratios of 6-DHG and curcumin accumulated by the *Zo*PKS2 cultures; compared to our modified M9 medium with glycerol, the other media increased the curcumin ratio. For example, replacement of glycerol with glucose in the modified M9 medium increased yields of 6-DHG (2.15 ± 0.05 mg L^-1^) 2-fold and curcumin (1.06 ± 0.13 mg L^-1^) 17-fold.

In summary, these findings indicate that there is significant room for future optimization of cultivation conditions for 6-DHG production, including testing different carbon sources, precursor substate feeding regimens and concentrations, testing different *E. coli* strains and applying metabolic engineering strategies to direct precursor substrate biosynthesis in *E. coli.* Many of the same strategies used to increase flavonoid and curcumin production by metabolically engineered microorganisms, may be applied to increase gingeroid yields as well ^13, 15–17^. The requirement for fatty acyl-CoA substrates poses additional challenges, including addressing growth inhibition ^45–47^, but also offers opportunities for metabolic engineering of hexanoyl-CoA and other medium-chain fatty acyl-CoA pools^48–50^ to access gingeroids with different acyl-chain length (e.g. 6/8/10-gingeroids that are known to be made by *Z. officinales* ^51^) and non-natural structurs. In this work, we chose the commonly used *At*4CL1 for its broad activity towards phenylpropanoic acids ^6-7, 35^, but replacement of *At*4CL1 with a dedicated feruloyl-CoA synthetase could easily increase production of 6-DHG ^52^. Likewise, different acyl-CoA ligases, including a different hexanoyl-CoA ligase ^33^ could be test for increasing yields along with the use of a robust microbial chassis such as *Pseudomonas putida* ^53^ that can better tolerate stress conditions.

## CONCLUSIONS

In this work, we describe the biosynthesis of 6-dehydrogingerdione (6-DHG) in engineered *E. coli* by a *bona fide* gingerdione polyketide synthase (PKS) identified in *Zingiber officinalis*. This enzyme (*Zo*PKS2) along with a close homolog (*Zo*PKS1) characterized as curcumin synthase, was identified in the ginger genome through homology-based searches, phylogenetic analysis and structural modeling. These analysis revealed chromosomal clustering of ZoPKS2 (and ZoPKS1) with a diketide synthase (DCS) and identified changes in its active side residues compared to curcumin synthase ZoPKS1 and closely related curcumin synthases (CURS) from *Curcuma longa*. We demonstrated that *Zo*PKS2 produces almost exclusively 6-DHG in recombinant *E. coli* when provided with hexanoyl-CoA as a starter substrate. When only feruloyl-CoA is available, *Zo*PKS2 will use it as a starter instead to efficiently produce curcumin in *E. coli.* In contrast, we found that ZoPKS1 is a curcumin synthase that produces only very small amounts of 6-DHG in the presence of hexanoyl-CoA. While yields for 6-DHG (∼ 2 mg L^-1^) were low, preliminary production experiments identified bottlenecks and opportunities for optimization of cultivation conditions and metabolic engineering strategies to increase yields and access 6-DHG compounds with different acyl-chain lengths and structures.

The switch of *Zo*PKS2 from a 6-DHG to an efficient curcumin synthase in the absence of an acyl-CoA starter substrate makes it an ideal candidate for the development of a microbial production platform that can produce either of these valuable bioactive natural products just by altering precursor feeding. Future mechanistic studies with purified enzymes are expected to provide insights into the substrate selectivity of *Zo*PKS2, which together with structural studies will inform protein design efforts to produce new bioactive gingeroids with different phenylpropanoic and/or carboxylic acid derived moieties. Finally, our engineered 6-DHG producing *E. coli* strain lends itself to be a convenient platform for the discovery and screening of additional oxidoreductases to produce e.g. gingerols, shogaols or paradols with known potent bioactivities.

## METHODS

### Chemicals and materials

All chemicals were purchased from Sigma-Aldrich, Inc. (St. Louis, MO, USA) unless otherwise noted. This includes curcumin (SIAL-C1386-5G), zingerone (88787-50MG), sodium hexanoate (hexanoic acid) (153745-2.5g), cinnamic acid (C80857-5g), and kanamycin (60615-5g). Analytical standard of 6-dehydrogingerdione (AST-E88643) was purchased from Neta Scientific (Marlton, NJ, USA). Coumaric acid (102576) was purchased from MP Biomedicals (Solon, OH, USA). Ferulic acid (ICN10168580), formic acid (270480250), chloramphenicol (502388044), methanol (A452SK4), acetonitrile (A998-4), and Tris-base (B152-5), CaCl_2_ (AC349610250), ammonium persulfate (AC327081000), and EDTA (S311-3) were purchased from Thermo Fischer Scientific (Carlsbad, CA, USA). Glycerol (G22025-0.5) and ammonium chloride (A20424-500) were purchased from Research Products International (Mounts prospect, IL, USA). PCRBio Verifi Mix used for colony PCR, Gen Clone 30 mm/13 mm diameter x 0.22 µm pore size syringe filters (25-240/25-239) used for extract filtration, and yeast extract (20-102) were purchased from Genessee Scientific (San Diego, CA, USA) and all other molecular biology kits and enzymes were obtained from New England Biolabs (Ipswhich, MA, USA). Oligonucleotides were ordered from Integrated DNA Technologies, Inc. (Coralville, IA, USA). Codon optimized genes (see below) were synthesized by Twist Bioscience (San Francisco, CA, USA). For overnight culturing, BD DIFCO LB-Broth (240230) was purchased from Becton-Dickinson (Franklin Lakes, NJ). For SDS-PAGE analysis of proteins, Coomassie brilliant blue was from Sigma-Aldrich and all other reagents (TEMED, Precision Plus Protein^TM^ prestained protein standard (161-0373), 30% acrylamide/bis-acrylamide solution (37.5:1) (161-0159)) were purchased from Bio-Rad (Hercules, CA, USA).

### Sequence identification, analysis and homology modeling

Putative PKS and DCS candidate genes from *Zingiber officinalis* were identified by performing a BLAST search of the originally as NCBI Bioproject submitted genome assembly (PRJNA 647255) ^26^ and NCBIs automatically created RefSeq (PRJNA736965). Searches were performed with the characterized curcumin biosynthetic genes from *Curcuma longa* CURS1 (BAH56226.1), CURS2 (BAH85780.1) CURS3 (BAH85781.1) and DCS (BAH56225.1) as well as with the bismethoxycurcumin synthase from *Oryza sativa* CUS (Q8LIL0.2) and chalcone synthase from *Arabidospsis thaliana* CHS (AAF23561.1). Sequence alignment and phylogenetic analysis of the resulting sequences (**Table S1**) were done with MEGA 12 ^54^. Alignments were performed with ClustalW (default settings) and evolutionary history inferred by the Neighbor-Joining method with a bootstrap test (500 replicates, percentage of replicate trees with associated taxa clustered together are shown). Evolutionary distances were calculated using the Poisson-correction method in MEGA 12.

Homology models of *Zo*PKS1 and *Zo*PKS2 were created with SWISS-MODEL ^55^ using the curcumin synthase homodimer (CURS1) as the template (PDB 3OV2). The resulting models (*Zo*PKS1: 81.5% sequence identity with CURS1, GMQE=0.95; *Zo*PKS2: 71.61% sequence identity with CURS1, GMQE=0.90) were visualized and aligned to the CURS1 template structure in PyMOL (Schrodinger, LLC. 2010. The PyMOL Molecular Graphics System, Version 3.1.6.1). Predictions of subcellular plant cell localization for *Zo*PKS1/2 and *Zo*DCS1/2 were performed with DeepLoc 2.1 ^29^ and LOCALIZER ^30^.

### Molecular cloning and plasmid construction

All molecular cloning was done in *E. coli* Top10 cells cultivated in SOC (20 g L-^1^ Tryptone, 5 g L^-1^ yeast extract, 10 mM NaCl, 10 mM MgCl_2_, 2.5 mM KCl, 10 mM MgSO_4_, 20 mM glucose) at 37 °C with appropriate antibiotics added: kanamycin (50 µg mL^-1^) for pET28a plasmids, chloramphenicol (30 µg mL^-1^) for pAC_mod_ plasmids. The TEDA method ^56^ was used for Gibson assembly of DNA fragments with forward and reverse primers for amplification of gene inserts and vector backbones. The codon optimized *Zo*DCS1, *Zo*DCS2, *Zo*PKS1, *Zo*PKS genes were first individually cloned into pET28a-His_6_. To create the *Zo*DCS/*Zo*PKS co-expression plasmid combinations, the P_T7_-RBS-His_6_-*Zo*PKS1/2 expression cassettes were amplified and assembled on the complementary strands of the pET28a-His_6_-*Zo*DCS1/2 plasmids with a 200-nucleotide spacer in between the two promoters. The codon optimized hexanoic acid CoA ligase *At*ACSmut (NP_001031974.2) ^36^ and *At*4CL1 was also first cloned into pET28a-His_6_ to add an N-terminal His_6_-tag to the enzyme. This sequence was then placed 160-nucleotides downstream of the His_6_-*At*4CL1 expression cassette cloned into our previously described pAC_mod_ expression plasmid ^6-7^. A list of all plasmids and strains, primer sequences, amino acid and nucleotide sequences of genes is provided in **Tables S3-S6.** Expression of the recombinant proteins in *E. coli* BL21(DE3) was checked by 12% SDS-PAGE as described below.

### Production experiments

For production studies, chemically competent (CaCl_2_) *E. coli* BL21 (DE3) cells were first co-transformed with the desired pAC_mod_ and pET28a plasmid combinations. Single colonies of the transformed cells were picked from selective LB agar plates to inoculate overnight cultures that were grown at 30 °C, 180 rpm in selective LB medium (50 µg mL^-1^ kanamycin, 30 µg mL^-1^ chloramphenicol).

Larger production cultures were started by inoculating (1:100) 1 mL overnight cultures into baffled flasks containing 100 mL modified M9 media (M9-Mod) (M9 media (6 g L^-1^ Na_2_HPO_4_, 3 g L^-1^ KH_2_PO_4_, 1 g L^-1^ NH_4_Cl, 0.5 g L^-1^ NaCl, 1 mM thiamine-HCl, 1 mM MgSO_4_, 0.1 mM CaCl_2_) containing 1.25 g L^-1^ yeast extract) supplemented with 5 g L^-1^ glycerol. Cultures were grown for 2 h (37 °C, 180 rpm) to an OD_600_ = 0.3-0.5, chilled on ice for ∼ 15 min to lower the temperature to 25 °C prior to inducing the T7-promoter controlled *Zo*PKS/DCS expression by adding 1 mM IPTG to the cultures. Induced cultures were then grown for an additional 3 h (25 °C, 180 rpm) to allow for protein expression prior to feeding with 1 mM (final concentration) of the desired precursor substrates from stock solutions of 200 mM ferulic acid (substituted for cinnamic or coumaric acid for phenylpropanoic acid CoA promiscuity experiments) in 0.3 M NaOH and 200 mM sodium hexanoate in MilliQ-H_2_O. Cultivation then continued (25 °C, 180 rpm) until analysis of product formation after 20 h post precursor feeding (25 h total cultivation time). All cultivations were performed in triplicate with biological replicates unless otherwise noted. Cell growth was followed by OD_600_ measurements with samples taken at the cultivation start (0 h), time of IPTG induction (2 h), addition of precursor substrates (5 h) and end of cultivation (25 h). Optical densities (OD) were measured spectrophotometrically at 600 nm in 96-well plates (Varioskan Lux, Thermo Fisher Scientific) and measured values were corrected to a 1 cm path length. At the end of the experiments, samples were removed for product analysis as described below and during the initial screening experiments, for analyses of recombinant enzyme expression by SDS-PAGE as described below. For comparison of curcumin and 6-DHG production in different media conditions, *Zo*PKS2/*Zo*DCS2/*At*4CL1/*At*ACSmut transformed *E. coli* BL21 was also grown in Luria Bertani (LB; 10 g L^-1^ tryptone, 5 g L^-1^ yeast extract, 5 g L^-1^ NaCl) and Terrific Broth (TB; 12 g L^-1^ tryptone, 24 g L^-1^ yeast extract, 0.4% glycerol (v/v), 2.3 g L^-1^ KH_2_PO_4_, 12.5 g L^-1^ K_2_HPO_4_) and modified M9-medium supplemented either with 5 g L^-1^ glycerol or 10 g L^-1^ glucose. Production experiments, including induction and feeding with ferulic and hexanoic acid, were performed as described above.

### SDS-PAGE analysis of protein expression

Total, insoluble and soluble protein expression of the individual enzymes (**Figure S3b-c**) was analysed by transforming *E. coli* BL21 cells with pET28 (*Zo*PKS/1/2, *Zo*DCS1/2) or pAC_mod_ (*At*ACSmut, *A*t4CL1) plasmids expressing the individual proteins or none (empty vector (EV) Ctrl). Cells were grown in 100 mL modified M9 medium with 5 g L^-1^ glycerol and 1.25 g L^-1^ yeast extract (37 °C, 180 rpm). pET28a transformed cells were induced with 1 mM IPTG at an OD_600_ = 0.3-0.5 and allowed to express for 3 h before harvesting of all cultures by centrifugation (10,000 x *g*, 10 min, 4 °C). pAC_mod_ transformed cells were not induced but treated with 1 mM IPTG for reproducibility and grown for the same total duration until harvesting. Cells were resuspended to a final OD_600_ = 10.0 with 0.1 M Na phosphate buffer pH 8.0 and sonicated for 10 min (1s ON, 2 s OFF at 40% amplitude, Branson Sonifier). The total protein (T) sample was prepared by mixing 500 µL of the crude lysate with 100 µL of 6x SDS-PAGE. A second 500 µL aliquot of the crude lysate was spun down (10,000 x *g*, 10 min, RT) and supernatant containing the soluble proteins (S) was removed and mixed with 100 µL of 6x-SDS-PAGE sample buffer. The remaining insoluble proteins in the pellet (P) were resuspended in 500 µL of 0.1 Na phosphate buffer pH 8.0 and then mixed with 100 µL of 6x-SDS-PAGE sample buffer. All SDS-PAGE samples were boiled at >99 °C for 20 min and chilled on ice for 2 min. Gel analysis was performed by loading 5 µL of sample or protein standard onto 12% SDS-PAGE gels and ran at 120 V for 90 min. The gels were stained with Coomassie Blue according to the manufacturers’ instructions. Analysis of protein expression by recombinant *E. coli* strains co-expressing different combinations of *Zo*PKS, *Zo*DCS and CoA ligases (**Figure S3**) were performed with IPTG induced and uninduced cultures grown for another 20 h following induction and feeding with 1 mM ferulic and hexanoic acid as in the production experiments. Cultures were grown in 100 mL modified M9 medium with 5 g L^-1^ glycerol and 1.25 g L^-1^ yeast extract at 25 °C, 180 rpm. Cells were harvested and total protein expression was analyzed as described above and compared to empty vector (EV) control cultures.

### Product extraction from recombinant cultures

For the initial product screening experiments (**Figure 4**) and profiling of feruloyl-CoA vs hexanoyl-CoA starter substrate preference (**Figure 5**), products were extracted from complete cultures (total culture), cell free media supernatant (cell free media) and spun down cell pellets (cell pellet). For the total culture extracts, 10 mL of *E. col*i culture was extracted twice with equal volume (10 mL) of ethyl acetate. The aqueous and organic phases were allowed to separate (∼1 h) before collecting the organic phases and drying under N_2_ and resuspension in 2 mL methanol. For the cell free media and cell pellet extractions, 10 mL of culture was centrifuged (10,000 x *g*, 10 min, 4 °C) to separate cells from media. The collected media supernatant was extracted as described for the total cell culture. The remaining cell pellet was resuspended by vigorous vortexing in 2 mL ice-cold methanol and allowed to chill on ice for 1 h before filtering through a 30 mm diameter x 0.22 µm pore PVDF syringe filter to remove cell debris. For phenylpropanoid-CoA promiscuity experiments (**Figure 6**) and comparing curcumin and 6-DHG production levels by recombinant *E. coli* strains or in different media (**Figure 7**), 1 mL of culture samples were used for complete culture (total), cell-free media and cell pellet extractions using the protocol described above with a modification of extraction volume to 1 mL (equal to the sample volume) of ethyl acetate for the complete culture (total) and cell-free media, and 200 µL of ice-cold 100% MeOH for the cell-pellet. The cell pellet was subjected to filtration using a 15 mm x 0.22 µm pore sized PVDF syringe filter for removal of the cell debris. All extraction samples were stored at −20 °C until further use.

### Analytical HPLC

For analysis of all culture extracts and concentrated fractions isolated by preparative HPLC (see below), 10-50 µL of extract were separated on an Agilent Eclipse XDB-C18 column (150 mm x 4.6 mm, 2.5 mL column volume) with a Agilent 1100 series HPLC equipped with an Agilent 1364 DAD detector for monitoring absorbances at 370 nm for the detection of curcumin and 6-DHG products from most experiments, and in addition at 320 nm for the detection of products from the phenylpropanoic acid feeding experiments (**Figure 6**) (curcumin λ_max_= 420-430 nm; 6-DHG λ_max_= ∼370 nm, ferulic acid λ_max_= 310-320 nm, coumaric acid λ_max_= ∼310 nm, cinnamic acid λ_max_= ∼270 nm). The detector was configured to acquire UV/Vis spectra from 200-600 nm. Separations were done at a flow rate of 1 mL min^-1^ and column temperature of 30 °C with mobile phases A (HPLC grade water with 0.01% trifluoro acetic acid (TFA)) and B (100% acetonitrile) using the following method: 5 min 5% B, 20 min gradient of 5%-95% B, 5 min of 95% B, 1 min gradient of 95% to 5% B, and then 5 min 5% B.

### Preparative HPLC for compound isolation

Compounds selected for subsequent identification by LC-MS/MS were purified from extracts by preparative HPLC on the same Agilent 1100 series HPLC with a Microsorb-MV, 100 Å, C18 reverse-phase column (250 mm x 4.6 mm) (Agilent, R0086200C5), column temperature at room temperature. Cell free media extracts from 10 mL cultures were dried under N_2_ gas as described above and then resuspended to a final concentration of 10 mg mL^-1^ in methanol. Peaks were isolated through preparative HPLC with an injection volume of 100 µl and separation using a flow rate of 1 mL min ^-1^ with mobile phases A (HPLC grade water and 0.1 % formic acid) and B (acetonitrile and 0.1% formic acid). The separation method was as follows: 0–5 min loading/wash with 5% B, 40 min gradient 5% to 100% B, 5 min wash with 100% B, 1 min solvent change from 100% to 5% B, and then 5 min re-equilibration with 5% B. Fractions were collected for the first 5 min (5 mL, 5% B) for the loading/wash phase of run, every 2 min (2 mL) over the course of the gradient (40 min total, 5% to 95% B), and then for a second wash (6 min, 100% B) and re-equilibration (6 min, 5 % B). Fractions of interest were identified, pooled based on their UV/Vis absorbance, extracted twice with equal volumes of ethyl acetate (2 mL-10 mL depending on peak size), dried under N_2_ gas, and resuspended in 100 µL of MeOH and stored in −20 °C. Each purified sample was checked to contain the desired compound peak by analytical HPLC as described above (see **Figure S5** for representative workflow examples).

### LC-MS

LC-MS/MS analysis of were done using a Thermo Fisher Scientific LTQ Orbitrap Velos mass spectrophotometer connected to an UltiMate 3000 UHPLC. Negative-ion electrospray ionization (ESI) was used for ionization. Separations were done with an Agilent Eclipse XDB-C18 column (150 mm x 4.6 mm, 2.5 mL column volume) at a flow rate of 1 mL min^-1^ with the column temperature set at room temperature and an injection volume of 10 µL. Mobile phases use were: mobile phase A: LC-MS grade water with 0.1% formic acid and mobile phase B: LC-MS grade acetonitrile with 0.1% formic acid. The separation method was as follows: 0-20 min gradient of 5% to 100% mobile phase B, 5 min of 100% mobile phase B, 1 min of solvent change from 100% to 5% mobile phase B, and then 5 min re-equilibration with 5% mobile phase B. The ion source parameters were as follows: negative-mode ESI, spray voltage at −4.00 kV, source heater temperature at 400 °C, sheath gas flow (N_2_) at 80 arbitrary units, auxiliary gas flow at 5 arbitrary units, and a capillary temperature at 300 °C. Mass analyzer was set to 7 scan events per cycle (one full MS1 scan followed by top 5 fragmentation plus one targeted fragmentation). Full MS1 scan range was between 100-1200 m/z with a resolution set to 30,000 in profile data form. Data dependent tandem mass spectrometry (MS/MS) product ions were generated using HCD normalized collision energy conditions of 25.0 with isolation width set to 2.0 and minimal signal required of 5,000 with resolution set to 7,500. MS2 fragmentation data was performed on the five most intense ions in each MS1 scan. For targeted fragmentation, isolation width was set at 1.5 m/z; HCD normalized collision energy was set at 30; resolution was set at 15.000. Maximum injection time: 10 ms for MS1 and 100 ms for MS2; AGC target: 1E6 for MS1 and 5E6 for MS2. Orbitrap was used for both the MS1 and MS2.

Data analysis was performed using the Thermo Fisher Scientific Xcalibur Qual Browser by filtering range of MS values to the nearest +/- 1.0 u (i.e. 289.00 – 290.00) for [M-H]^-^ m/z values for the following compounds as follows: curcumin ([M-H]^-^ = 367.12), 6-dehydrogingerdione (6-DHG) ([M-H]^-^ = 289.14), zingerone ([M-H]^-^ = 193.09), feruloylacetone ([M-H]^-^ = 233.08), feruloyl-ß-ketoacid ([M-H]^-^ = 237.08), bismethoxycurcumin ([M-H]^-^ = 307.10), dicinnamoylmethane ([M-H]^-^ = 276.12), *p*-coumaroylhexanoylmethane ([M-H]^-^ =259.13), cinnamoylhexanoylmethane (non-reduced [M-H]^-^ = 243.15, reduced [M-H]^-^ = 227.15), *p-* coumaroyl-ß-ketoacid ([M-H]^-^ = 205.05). Parent mass values of the unknown stress peaks were determined based on their retention time. All experimental LC-MS parent ions and fragmentations were compared to 6-DHG, curcumin, and zingerone analytical standards. Peaks of the LC-MS analysis were compared to peaks from the analytical HPLC performed above and by performing the analytical HPLC separation with the modified LC-MS mobile phases.

### Statistical analysis and reproducibility

Quantification of curcumin and 6-DHG products was performed by fitting a linear regression model of standard curves generated with authentic reference compounds (R-squared values ≥ 0.99). Peak areas were calculated using the Agilent OpenLab CDS program. Calculations and graphing of data and HPLC chromatograms were performed by exporting data as CVS files for use with Microsoft Excel 365 with following calculations. UV/Vis Maximum was calculated from the max value (MAX function) of the absorbance vector as input for the LOOKUP function from the corresponding Wavelength vector to return the max wavelength value corresponding with the maximum absorbance value. Relative Intensity of the chromatogram was calculated as the percentage ratio of each individual absorbance value divided by the maximum intensity value from each chromatogram. Variations of the Relative Intensity calculation were performed for scalability and noted in the figure legends. Mean and standard deviations for product concentrations and OD_600_ values were calculated using Average and STDEV functions in Microsoft Excel 365. Unless otherwise noted, production experiments were performed with three biological replicates (i.e. separately inoculated cultures).

## Data availability

All data generated for this work are presented in this manuscript and its Supporting Information. Plasmids made in this work will be made available subject to an MTA that can be requested by contacting the corresponding author Prof. Schmidt-Dannert, who will respond to requests within a week.

## SUPPORTING INFORMATION

Supporting information provides: Sequence and homology model alignments, additional HPLC profiles for control cultures, SDS-PAGE analysis of protein expression, representative HPLC chromatograms for compound isolation, U/Vis and MS spectral data, culture growth data, list of plasmids and strains, nucleotide and/or amino acid sequences of primers and proteins used.

## AUTHOR CONTRIBUTIONS

E.L.P. contributed to the experimental design, created the genetic constructs, performed the production experiments, isolated and identified products, analyzed data and wrote the manuscript. S-Y.K. contributed the experimental design, construction of plasmid, identification of compounds, data analysis and writing of the manuscript. F.J.G. contributed to the construction of the bi-directional plasmid, analysis and identification of products, experimental design and writing of the manuscript. C.S.-D. conceived, conceptualized, and directed the overall project, analyzed the data, performed the bioinformatics analysis, and wrote the manuscript together with E.L.P. and F.J.G and S.-Y.K.

## ACKNOWLEDGMENTS

We would like to thank Dr. Yudi Rusman and Prof. Christine Salomon, Center of Drug Discovery, University of Minnesota, for consultation on the optimization compound extractions and analysis. LC-MS analysis was done using shared mass spectrometry resources by the University of Minnesota’s Masonic Cancer Center that is supported in part by NIH P30 CA77598. This research was sponsored by the BioTechnology Institute at the University of Minnesota and funds provided to C.S.D. through the Kirkwood Chair Endowment. E.L.P was supported by a predoctoral National Institutes of Health traineeship (2T32GM008347).

