## Supporting Information for "Identification of type III polyketide synthases from Ginger for dehydrogingerdione and curcumin biosynthesis by engineered *Escherichia coli*"

*Corresponding author

Prof. Claudia Schmidt-Dannert

140 Gortner Laboratory, University of Minnesota, 1479 Gortner Avenue, St. Paul, MN 55108

### SUPPLEMENTARY FIGURES

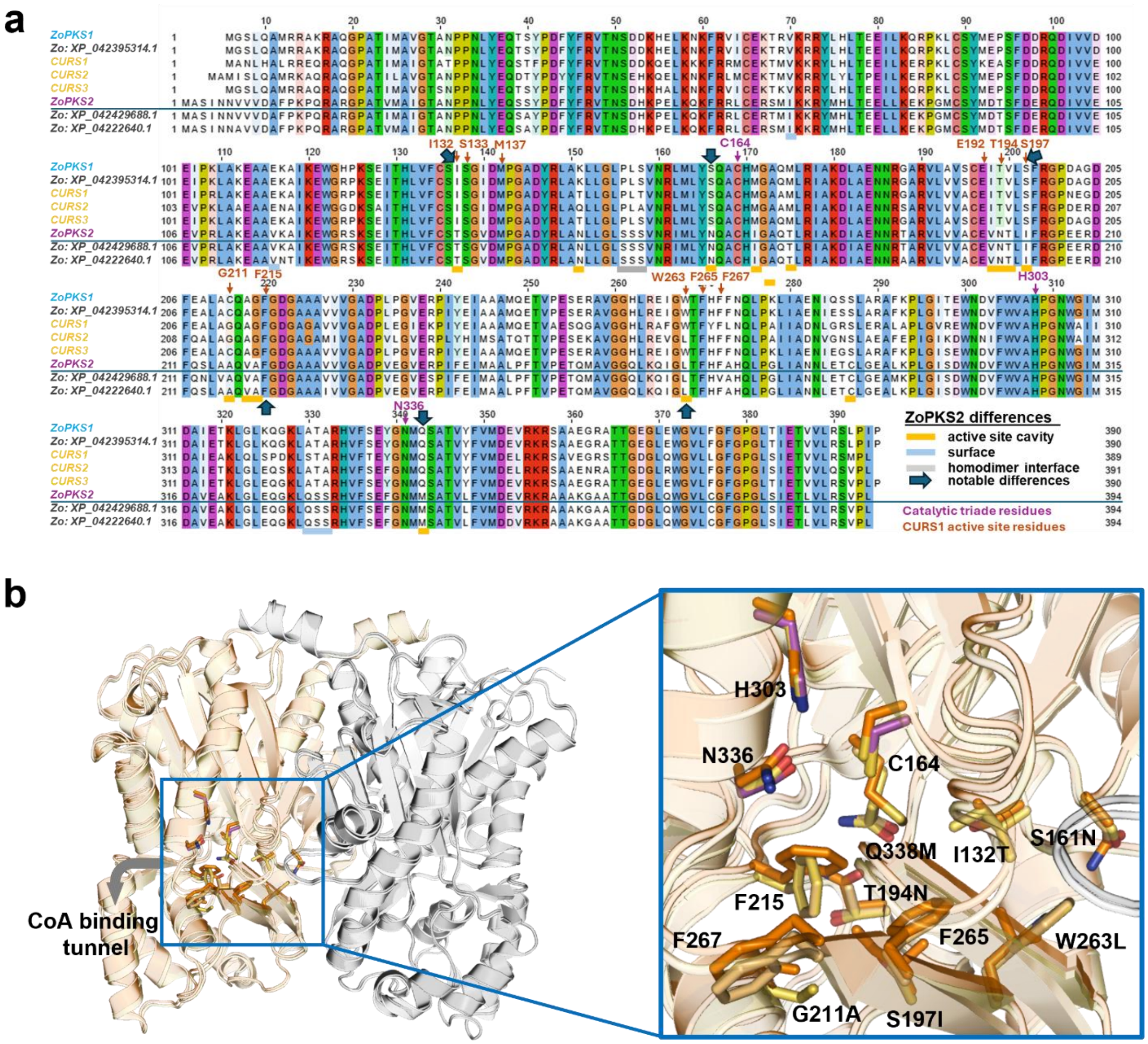

**Figure S1: Sequence alignments and alignments of *Zo*PKS1 and *Zo*PKS2 structural models with CURS1.**

**(a)** Protein sequence alignment of *Zo*PKS1 and 2 and their respective alleles with the known curcumin biosynthetic proteins CURS1-3 from *Curcuma longa*. Highlighted are residues that are notably different in *Zo*PKS, including in the active site. Purple and orange colors denote catalytic triad and CURS1 active site residues, respectively. Accession numbers and sequences are listed in **Table S1**. **(b)** Alignment of both *Zo*PKS1 (yellow backbone) and *Zo*PKS2 (orange backbone) homology models with the structure of CURS1 (PDB 2OV2) (yellow-orange backbone) as in **Figure 2**. The expansion shows residues lining the active site, including the catalytic triad (purple, CURS1 residues).

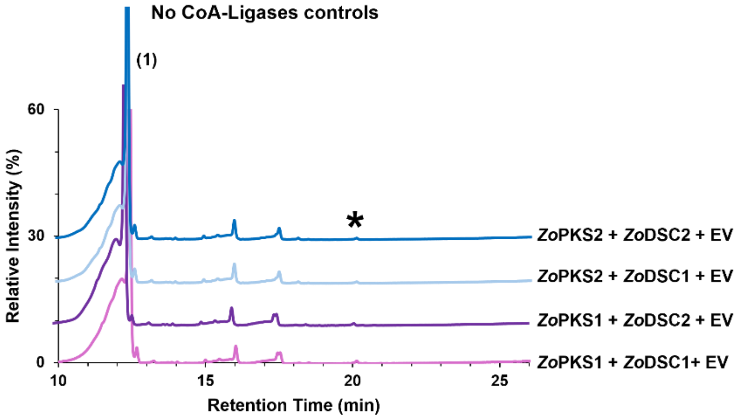

**Figure S2: HPLC profiles of extracts from control cultures from Figure 4.**

Recombinant *E. coli* BL21 cultures # co-expressing *Zo*PKS/DSC combinations and harboring pAC-empty plasmids without CoA ligases (see **Figure 4a**) were fed with hexanoic and ferulic acid, and after 20 h of conversion, total cultures were extracted and compounds analyzed by HPLC. HPLC traces are stacked for clarity, and importantly, each chromatogram is scaled to 60% of its highest (100%) intensity peak to visualize less abundant peak. #All E. coli cultures were grown in modified M9 medium with 5 g $L^{-1}$ glycerol at 25 ºC and fed with 1 mM CoA-precursor substrates after 3 h of IPTG induction of *Zo*PKS/DCS expression. Data were collected from three cultures and data points are the mean of three independent biological replicates. Representative HPLC profiles are shown.

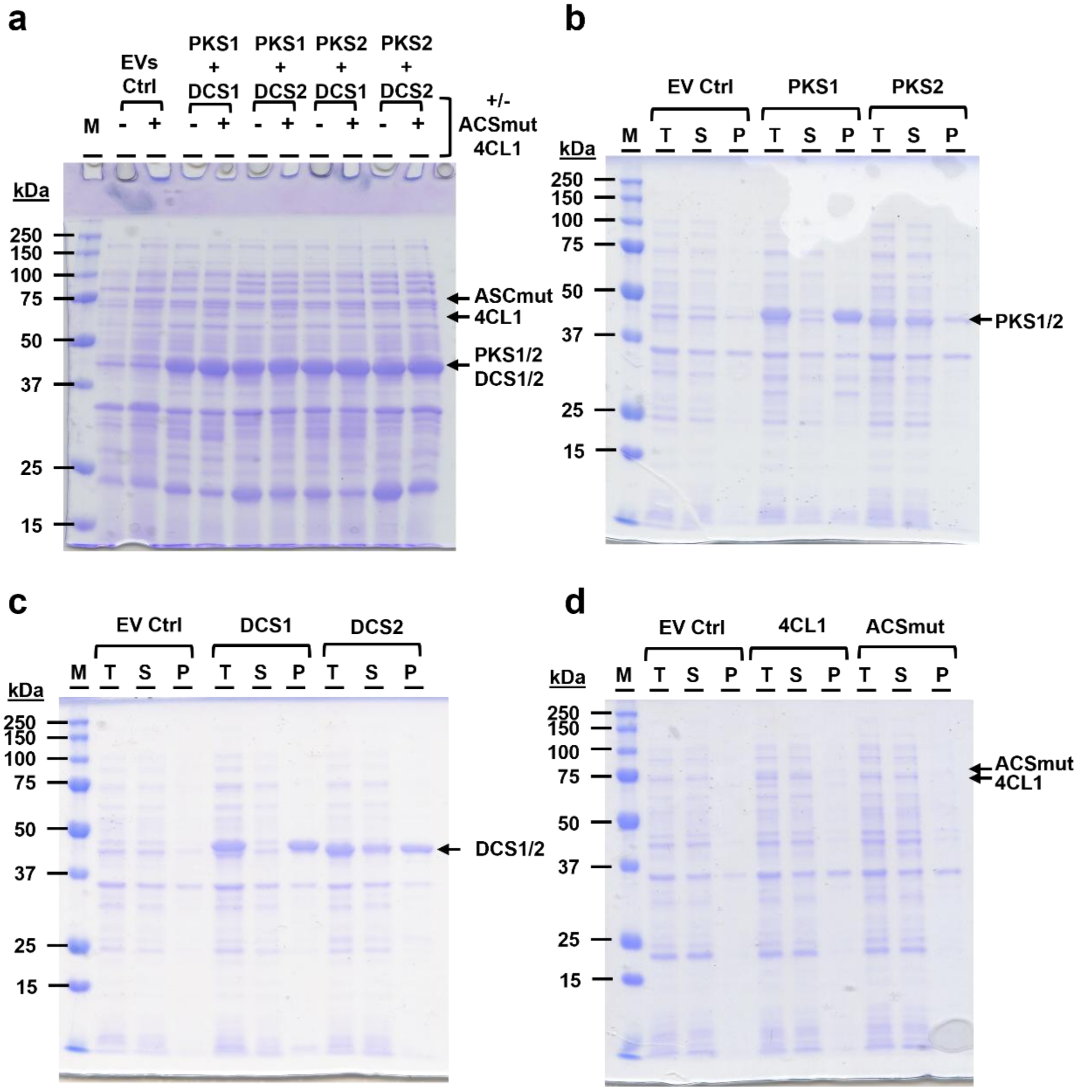

MWs: His$_6$-*At*ACSmut (75 kDa), His$_6$-*At*4CL1 (65 kDa), His$_6$-*Zo*PKS1/2 & His$_6$-*Zo*DCS1/2 (45 kDa)

**Figure S3: Analysis of recombinant protein expression and solubility.**

**(a)** SDS-PAGE gel analysis of protein expression by recombinant *E. coli* strains co-expressing different combinations of *Zo*PKS and *Zo*DCS enzymes (IPTG inducible, pET28 plasmid) with (+) or without (-) CoA ligases (*At*ACSmut, *At*4CL1, constitutive, pAC$_{mod}$ plasmid) 20 h after IPTG induction and feeding with 1 mM ferulic acid and 1 mM hexanoic acid. Cultures were grown in modified M9 medium with 5 g L$^{-1}$ glycerol at 25 ºC. Cells were spun down and used for total protein expression analysis and compared to an empty vector (EVs) control culture (Ctrl). **(b-c)** Soluble protein expression of individual enzymes was analysed by transforming *E. coli* cells with pET28a (*Zo*PKS/1/2, *Zo*DCS1/2) or pAC$_{mod}$ (*At*ACSmut, *A*t4CL1) plasmids (**Table S3**) expressing the individual proteins or none (EV Ctrl). Cells were grown in the same production medium as in (a) at 37 ºC. Transformed cells were induced with 1 mM IPTG at an OD$_{600}$ of 0.3-0.5 and allowed to express for 3 h before harvesting of all cultures. Harvested cells were washed, normalized and lysed (T: total protein), centrifuged to fractionate cell pellet (P: pellet, insoluble protein) and supernatant (S: soluble protein) for separation by SDS-PAGE.

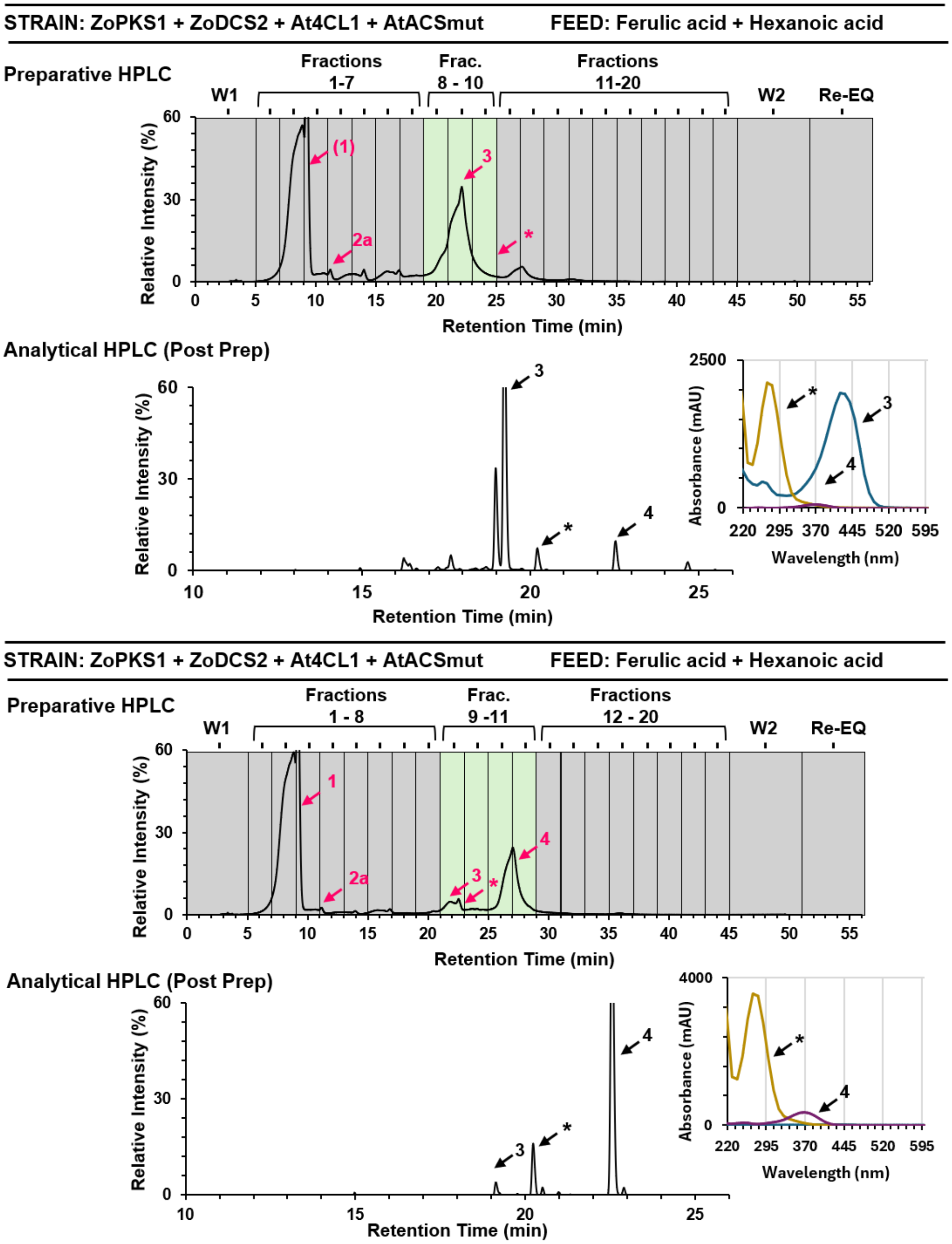

**Figure S4. Chromatogram examples of preparative and analytical HPLC of compound peaks subjected to LC-MS/MS analysis.**

Two representative examples (top bar shows strains and substrate feeding) are shown to illustrate the workflow used for the isolation of compound peaks from recombinant culture extracts. Preparative HPCL from total culture extracts from recombinant *E. coli* strains was performed as described in the Materials and Methods. Compound peaks are labeled according to numbering in **Figure 1** and were identified and selected based on their absorbance spectra and retention times of standard compounds. Fractions (2 mL) (highlighted in green as examples) representing peaks of interest were collected, pooled, dried under $N_2$ gas, resuspended in MeOH, and stored at -20 °C until further use. Collection of the desired compound peaks was confirmed by analytical HPLC of the purified samples and comparing retention times and UV/Vis spectra to the original HPLC product profiles of culture extracts and standards (see Methods).

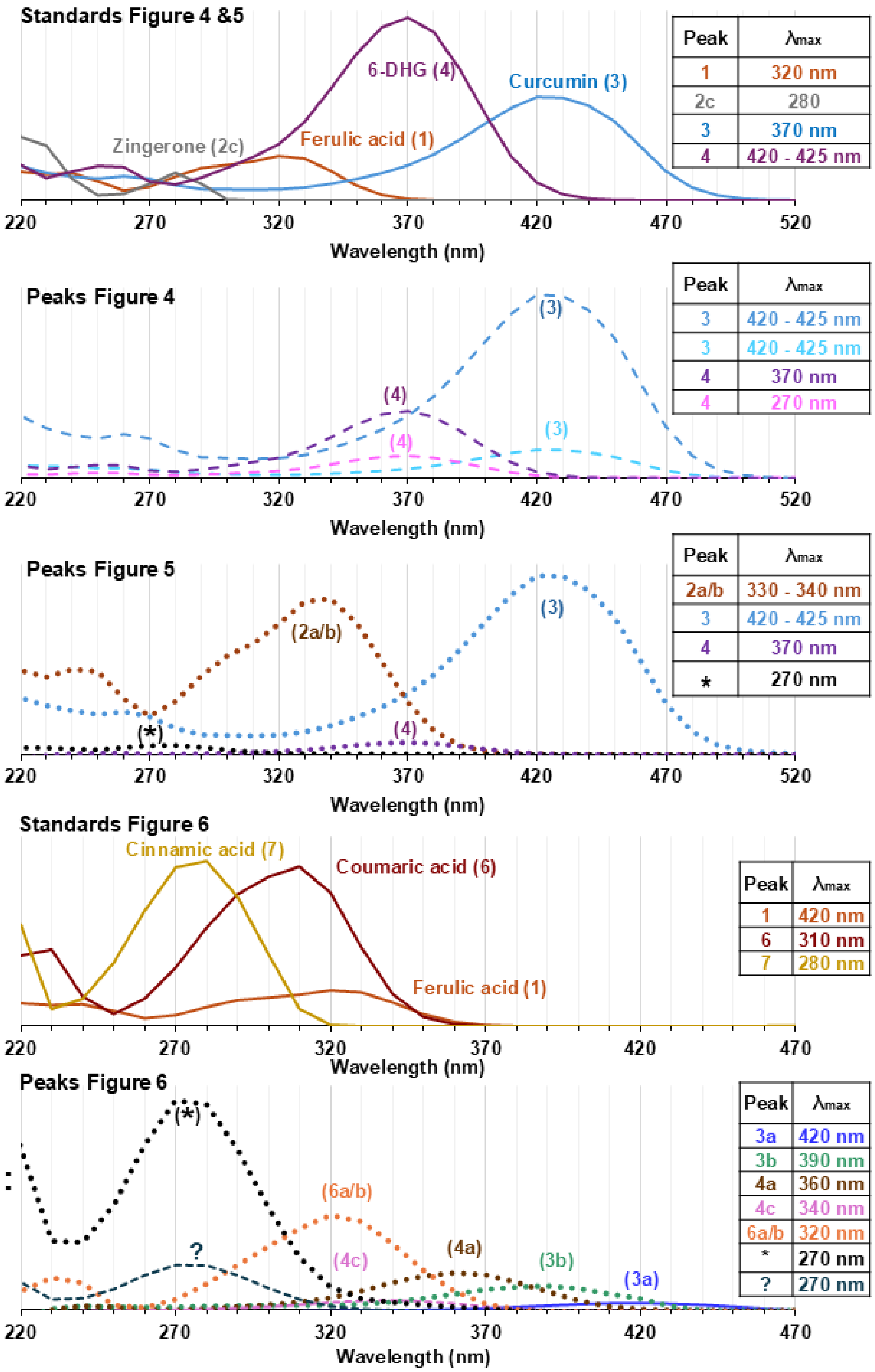

**Figure S5: HPLC-DAD UV/Vis spectra of standard compounds and major peaks from culture extracts.**

UV/Vis spectra and measured absorption maxima of authentic standards and representative peaks from **Figures 4-6** are shown. Peaks are labeled using the compound numbers of **Figure 1**, which are used in all Figures.

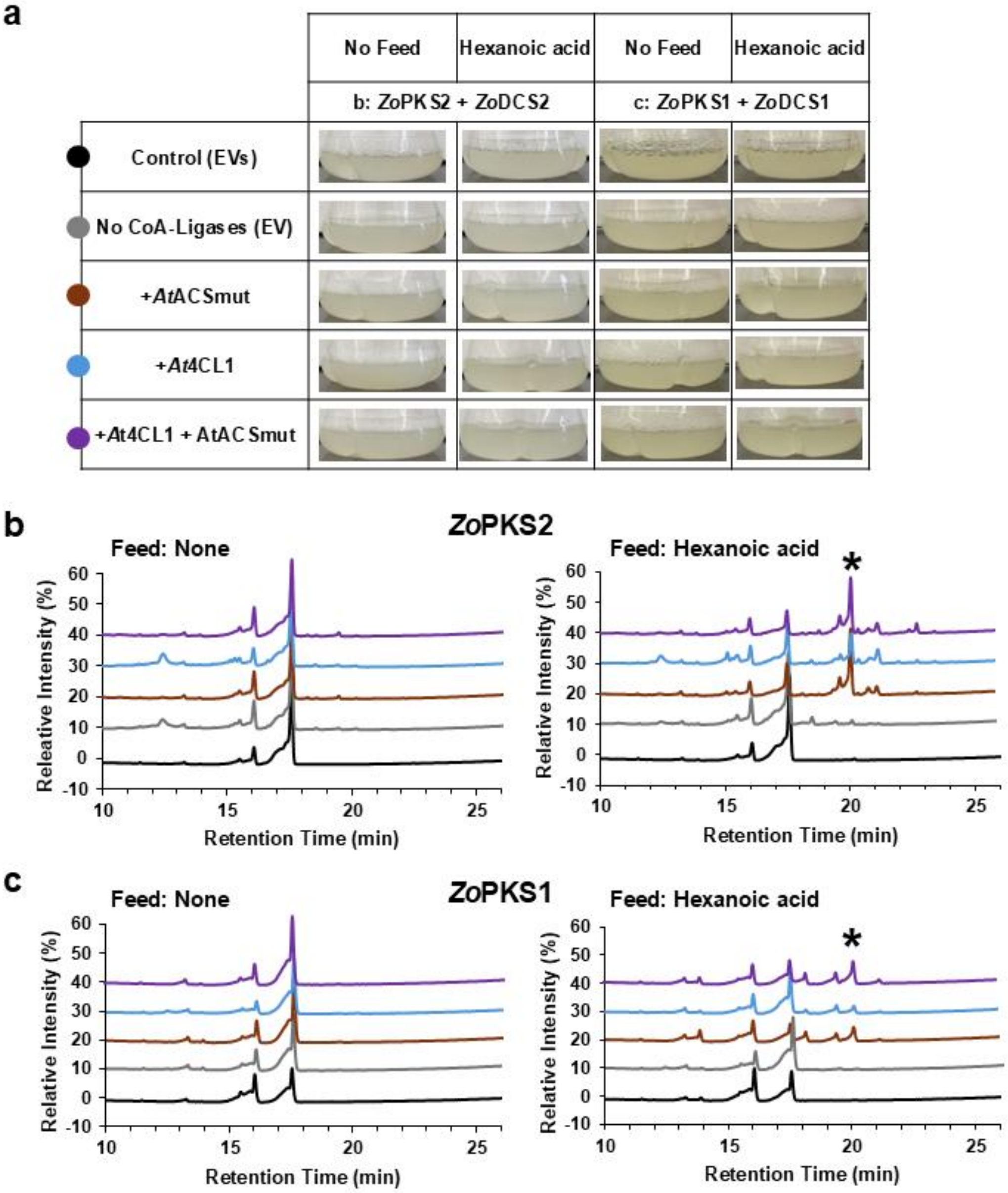

**Figure S6: Cultures and HPLC profiles of culture media extracts from control cultures in Figure 5.**

**(a)** Recombinant *E. coli* BL21[#] cultures harboring *Zo*PKS1 or *Zo*PKS2 with *Zo*DCS2 and either *At*4CL1, *At*ACSmut, both, or empty vector (EV) producing no significant color products different than that of the double empty vector control when fed hexanoic acid as compared to the No Feed control. **(b, c)** HPLC analysis of cell free extracts monitored at 370 nm are shown for *Zo*PKS2 **(b)** or *Zo*PKS1 **(c)** co-expressing the gene combinations shown in panel a. Colors of the chromatogram trace correspond to different CoA ligase combinations and empty vector(s) (EV(s) controls. [#]All *E. coli* cultures were grown in modified M9 medium with 5 g $L^{-1}$ glycerol at 25 ºC and fed with 1 mM precursor substrates after 3 h of IPTG induction of *Zo*PKS/DCS expression. After 20 h of conversion, cultures were processed for analysis. Extracts and data were obtained from two independent cultures for each experimental condition. One HPLC profile for each condition is shown. Compound/peak labels follow numbering in **Figure 1**.

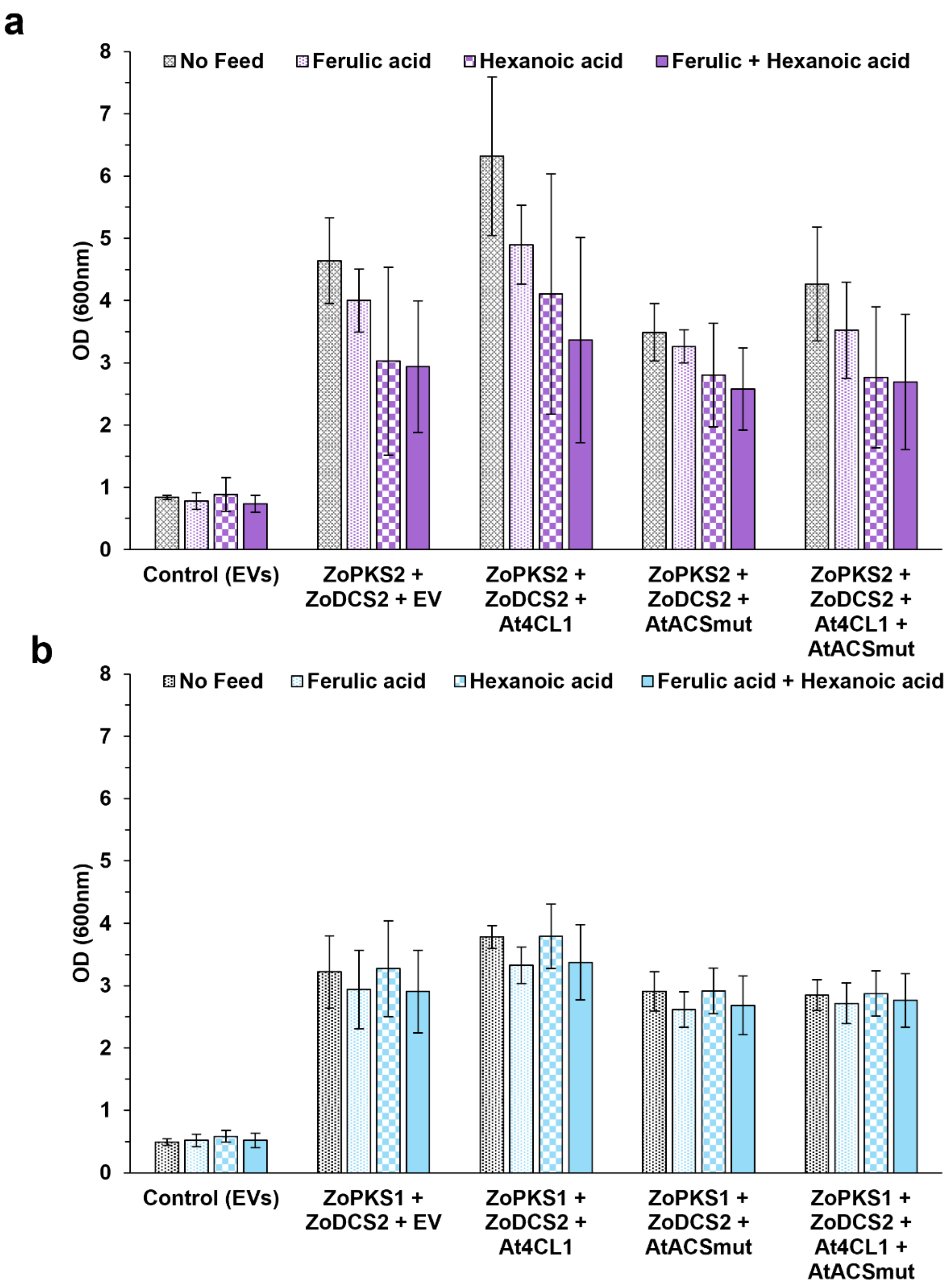

**Figure S7: Comparison of optical densities reached by recombinant strains after 20 h of conversion.**

Final optical densities ($OD_{600}$) reached of recombinant cultures (from **Figure 5a**) expressing *Zo*PKS2 (**a**) or *Zo*PKS1 (**b**) in combinations with *Zo*DCS2 and CoA-Ligases and fed with different substrates or none. Control empty vectors (EVs) cultures are co-transformed with empty $pAC_{mod}$ and pET28a. Graphs shown are an average of two replicated cultures per condition and the error-bars show variability between the two cultures.

### SUPPLEMENTARY TABLES

**Table S1: List of accession numbers of type III PKS protein sequence used for sequence alignments and phylogenetic analysis.**

| Sequence Identifier | Sequence |
|---|---|
| Zo: XP_042470361.1 chalcone synthase 2 like [*Zingiber officinale*] | MAKLVVSEIRRSQRAEGAAAVLAIGTANPPNVVYQADYPDYYFRITRSEHLTELKEKFKRMCDKSMIRKRHMYLTEEILKENPKMCAYMEPSLDARQDIVVVEVPKMGKEAAAKAIKEWGQPKSKITHLVFCTTSGVDMPGADYQLTKLLGLRPSVNRFMMYQQGCFAGGTVLRLAKDLAENNRGARVLVVCSEITAVTFRGPSDSHLDSMVGQALFADGAAAIIVGADPDPATERPLFELVSASQTILPDSEGAIDGHLREVGLTFHLLKDVPGLISKNIEKSLAEAFKPLGISDWNSLFWIAHPGGPAILDQVEAKLALDKNKMKATRDVLSEYGNMSSACVLFILDEMRRRSAEEGKATTGEGLEWGVLFGFGPGLTVETVVLHSVPIAAAH |
| Zo:_XP_042459246.1 phenylpropanoylacetylCoA synthaselike [*Zingiber officinale*] | METTVFRTTQSAEGPATILAIGTANPANVVDQSAYPDFYFRVTDCEHLQDLKAKFRRICERAAIRKRHLYLTEDILRKNPSLLAPMAPSFDARQAIVVAAIPELAKEAAEKAIKEWGRPKSDITHLVFCSASGVDMPGSDLQLLKMLGLPMSVNRVMLYNVGCHAGGTALRVAKDLAENNRGARVLAVCSEVTVLSYRGPDAAHIESLFVQALFGDGAAALVVGSDPIDGVERSIFEVASASQVMLPESAEAVGGHLREIGLTFHLKSQLPAIIASNIEQSLVAACAPLGLTDWNQLFWVVHPGGRAILDQVEARLGLQKERLAATRHVLSEFGNMQSATVLFILDEMRKRSAAEGRATTGEGDEWGVLLGFGPGLSIETVVLRSVPT |
| Zo: XP_042454519.1 phenylpropanoylacetylCoA synthase like [*Zingiber officinale*] | MEVNGYRIIHSADGPATILAIGTANPTNVVDQNAYPDFYFRVTNSEHLQELKAKFRRICEKAAIRKRHLYLTEEILRENPSLLAPMAPSFDARQQIVVEAVPKLAKEAAEKAIKEWGRPKSDITHLVFCSASGIDMPGSDLQLLKLLGLPLCVNRVMLYNVGCHAGGTALRVAKDLAENNRGARVLAVCSEVTVLSYRGPHPAHIESLFVQALFGDGAAALVVGSDPVDGVERPIFEIASASQVMLPESEEAVGGHLREIGLTFHLKSQLPSIIASNIEQSLTTACSPLGLSDWNQLFWTVHPGGRAILDQVEARLGLEKDRLAATRHVLSEYGNMQSATVLFILDEMRKRSAADGHATTGEGLDWGVLLAFGPGLSIETVVLHSCKLN |
| Zo: XP_042440615.1 chalcone synthase 2 like [*Zingiber officinale*] | MATLQEIRWAQRAEGRAAVLAIGTATPANVVYQADYPDQYFRMTKSEHLTELKEKFKRICEKTMIRKRYTHLTEEMLLENPNMCAYMEPSLDERQDIVAVEVPKLGKEAAAKAIEEWGQPKFKITHLVFCTTSGVDMPGADYQITKLLGLCPSVNRFMLYQQGCFAGGTALRLAKDLAENNRGARVLVVCSEITALAFRGPSESHQESLVAQALFGDGAGAVIVGADPNPETERPLFELVSASQTILPDSEGAIVYGRLGEAGFMVHLVKDVPALISKNIEKSLVEAFAPLGIDDWNTIFWIVHPGGSAILDQVEAKLALGKEKMAASRQVLSEYGNMSSPSVLFILDEMRRRSAEDGKATTGDGFHWGVLFGFGPGFTVETVVMHSMPIDY |
| Zo: XP_042439780.1 curcumin synthase 2 like [*Zingiber officinale*] | MAMISLQAMRKAQRAQGPATILSVGTANPPNLYEQTTYPDYYFRVTNSEDKQELKSKFRRMCEKTMVKRRYLYLTPEILKERPKLCSYMEPSFDDRQDIVVEEIPKLAAEAAEKAIKEWGGEKSAITHLVFCSISGIDMPGADYRLAKLLGLPLTVNRLMLYSQACHMGAAMLRIAKDIAENNRSARVLVVACEITVLSFRGPDERDFQALAGQAGFGDGAAAVVVGADPVPGVERPLYHIMSATQTTVPESEKAVGGHLREVGLTFHSFNQLPAIIADNVGSSLAEAFEPIGIKDWNDIFWVAHPGNWAIMNAIETKLGLEQSKLATARHVFSEFGNMQSATVYFVMDELRKRSAAENRETTGDGLRWGVLFGFGPGISIETVVLQSVPL |
| Zo:_XP_042439438.1 chalcone synthase 2 like [*Zingiber officinale*] | MAKVQEIRRSQRAEGPAAVLAIGTATPANVVYQADYADYYFRMTKSEHLTELKEKFKRMCDKSMIRKRYMHLTEEMLSENPNMCAYMEPSLDERQDIAVVEVPKLGKEAAAKAIKEWGQPKSKITHLIFCTTSGVDMPGADYQITKLLGLRPSVNRFMIYKQGCFAGGTVLRLAKDLAENNRGARVLAVCSEIMALKFRGPSESHLDNLVGQALFGDGAGAVIVGADPDLETERPLFELVSASQTILPDSEGAIGGHLREVGLIFHLLKDVPGLISKNIEKSLVEAFAPLGIDDWNSIFWIAHPGGPAILDQVEAKLALEKEKMTATRQVLSEYGNMSSACVLFILDEMRRKSAEEGKATTGEGFNWGVLLSFGPGITVETVVLHSKQINY |

| | |
|---|---|
| Zo:_XP_042439293.1 phenylpropanoylacetylCoA synthase like [*Zingiber officinale*] | MATIDASGKAADGPATILAIGTANPSNVVDQMQYPEYYFRITNSQYKTDLMHK<br>FQRVCEKSMIGKRHMCLTEEILKENPELCAYMAPSFDVRQKIVLAEVPRLAAE<br>AADKAIKEWGQPASDITHLVFSSSAGVDLLGVDCRLLQLLGLSPRVRRVMLYN<br>IGCHAGGTALRVAKDLAENNKGARVMVVCSELNVMFFRGPDDHHFENLIAQA<br>LFGDGAAVVIVGADPKEAERPIYELASAAQVMLPESEEMVAGHLREIGLTFHL<br>GSKLPAVVGANIQRCLEVSFAPMGVSNWNDLFWIVHPGGRAIVDQVEMSAGL<br>GAGKLAATRHVLREYGNMQSASVLFIMDEMRKRSAAEGCTTTGQGCEWGVL<br>FGFGPGLTVETVVLHALPI |
| Zo:_XP_042439182.1 chalcone synthase 2 like [*Zingiber officinale*] | MDKVQEIRRSQRAEGPAAVLAIGTATPANVVYQADYADYYFRMTKSEHLTEL<br>KEKFKRMCDKSMIRKRYMHLTEEMLRENPNMCAYMEPSLDERQDIVVVEVP<br>KLGKEAAAKAIKEWGQPKSKITHLIFCTTSGVDMPGADYQITKLLGLRPSVNR<br>FMMYQQGCFAGGTVLRLAKDLAENNRGARVLVVCSEITAVTFRGPSESHLDSL<br>VGQALFGDGAGAVIVGADPDLETELPLFELVSASQTILPDSEGAIDGHLREVGL<br>TFHLLKDVPGLISKNIEKSLVEAFAPLGIDDWNSIFWIAHPGGPAILDQVEAKLA<br>LEKEKMAASRQVLSEYGNMSSACVLFILDEMRRKSAEEGKATTGEGFNWGV<br>LFGFGPGLTVETVVLHSKPINY |
| Zo:_XP_042438998.1 curcumin synthase 1 [*Zingiber officinale*] | MASLHALRREQRAQGPATIMAIGTATPPNLYEQSTFPDFYFRVTNSDDKQELKE<br>KFRRICNKTMVKKRYLYLTEEILKERPKLCSYKEPSFDDRQDIVVEEIPKLAKE<br>AAEKAIKEWGRPKSEITHLVFCSISGIDMPGADYRLATLLGLPLTVNRLMIYSQ<br>ACHMGAAMLRIAKDLAENNRGARVLVVACEITVLSFRGPNERDFEALAGQAG<br>FADGAAAVVVGADPLERVEKPIYEIAAAMQETVAESQEAVGGHLRAFGWTFY<br>FLNQLPAIISNNIGKSLERALVPLGVREWNDVFWVAHPGNWAIMDAIEAKLQL<br>TPDKLSTARHVFSEYGNMQSATVYFVMDELRKRSAVEGRSTTGDGLQWGVLF<br>GFGPGLSIETVVLRSMPL |
| Zo:_XP_042436018.1 chalcone synthase 2 like [*Zingiber officinale*] | MATLQEIRWAQRTEGRAAVLAIGTATPANVVYQADYPDQYFRMTKSEHLTELK<br>EKFKRICEKTMIRKRYTHLTEEMLLENPNMCAYMEPSLDERQDIVAVEVPKLG<br>KEAAAKAIEEWGQPKFKITHLVFCTTSGVDMPGADYQITKLLGLCPSVNRFML<br>YQQGCFAGGTALRLAKDLAENNRGARVLVVCSEITAVAFRGPSESHQESLVAQ<br>ALFGDGAGAVIVGADPNPETERPLFELVSASQTILPDSEGAVYGRLGEAGFMV<br>HLVKDVPALISKNIEKSLVEAFAPLGIDDWNTIFWIVHPGGSAILDQVEAKLAL<br>GKEKMAASRQVLSEYGNMSSPSVLFILDEMRRRSAEDGKATTGDGFHWGVL<br>FGFGPGFTVETVVLHSMPINY |
| Zo:_XP_042435447.1 curcumin synthase 2 [*Zingiber officinale*] | MAMISLQAMRKAQRAQGPATILSVGTANPPNLYEQTTYPDYYFRVTNSEDKQ<br>ELKSKFRRMCEKTMVKRRYLYLTPEILKERPKLCSYMEPSFDDRQDIVVEEIPK<br>LAAEAAEKAIKEWGGEKSAITHLVFCSISGIDMPGADYRLAKLLGLPLTVNRL<br>MLYSQACHMGAAMLRIAKDIAENNRSARVLVVACEITVLSFRGPDERDFQAL<br>AGQAGFGDGAAAVVVGADPVPGVERPLYHIMSATQTTVPESEKAVGGHLREV<br>GLTFHSFNQLPAIIADNVGSSLAEAFEPIGIKDWNDIFWVAHPGNWAIMNAIET<br>KLGLEQSKLATARHVFSEFGNMQSATVYFVMDELRKRSAAENRATTGDGLRW<br>GVLFGFGPGISIETVVLQSVPL |
| Zo:_XP_042435235.1 chalcone synthase 2 like [*Zingiber officinale*] | MAKVQEIRRSQRAEGPAAILAIGTATPANVVYQADYADNYFRMTKSEHLTELK<br>EKFKRMCDKSMIRKRYMHLTEEMLRENPNMCAYMEPSLDERQEIVVVEVPKL<br>GKEAAAKAIKEWGQPKSKITHLIFCTTTSVDMPVADYQITKLLGLRPSVNRYM<br>MYQQGCFAAGTVLRLAKDLAENNRGARVLVVCSEITALTFRGPSESHLDNLVG<br>QALFADGAGAVIVGADPDLETERPLFELVSASQTILPDSEGAIHGDLREAGLIIH<br>VLKDVPELISKNIEKSLVDAFAPLGVDDWNSIFWIAHPGGPAILDQIEAKLGLE<br>KEKMAAARQVLSEYGNMVSACVLFILDEMRRRSAEEGKATTGDGFNWGVLF<br>GFGPGLTVETVVLHSKPINY |
| Zo:_XP_042435168.1 phenylpropanoylacetylCoA synthase like [*Zingiber officinale*] | MATIDASGKAADGPATILAIGTANPSNVVDQMQYPEYYFRITNSQYKTDLMHK<br>FRRVCEKSMIGKRHMCLTEEILKENPELCAYMAPSFDVRQKIVLAEVPRLAEE<br>AADKAIKEWGQPASDITHLVFSSSAGVDLLGVDCRLLQLLGLSPRVRRVMLYN<br>IGCHAGGTALRVAKDLAENNKGARVMVVCSELNVMFFRGPDDHHFENLIAQA<br>LFGDGAAAVIVGADPKEAERPIYELASAAQVMLPESEEMVAGHLREIGLTFHL<br>GSKLPAVVGANIQRCLEVSFAPMGVSNWNDLFWIVHPGGRAIVDQVEMSAGL<br>GAGKLAATRHVLREYGNMQSASVLFIMDEMRKRSATEGCTTTGEGCEWGVL<br>FGFGPGLTVETVVLHALPN |

| | |
|---|---|
| Zo:_XP_042434761.1 chalcone synthase 2 like [*Zingiber officinale*] | MAKVQEIRRSQRAEGPAAVLAIGTATPANVVYQADYADYYFRMTKSEHLTEL<br>KEKFKRMCDKSMIRKRYMHLTEEMLSENPNMCAYMEPSLDERQDIAVVEVPK<br>LGKEAAAKAIKEWGQPKSKITHLIFCTTSGVDMPGADYQITKLLGLRPSVNRF<br>MIYQQGCFAGGTVLRLAKDLAENNRGARVLVVCSEITAVKFRGPSESHLDNLL<br>GQALFGDGAGAVIVGADPDLETERPLFELVSASQTILPDSEGAIGGHLREVGLI<br>FHLLKDVPGLISKNIEKSLVEAFAPLGIDDWNSIFWIVHPGGPAILDQVEAKLAL<br>EKEKMTATWQVLSEYGNMSSACVLFILDEMRRKSAEEGKATTGEGFNWGVL<br>FGFGPGLTVETVVLHSKPINY |
| Zo:_XP_042434725.1 chalcone synthase 2 like [*Zingiber officinale*] | MDKVQEIRRSQRAEGPAAVLAIGTATPANVVYQADYADYYFRMTKSEHLTEL<br>KEKFKRMCDKSMIRKRYMHLTEEMLRENPNMCAYMEPSLDERQDIVVVEVP<br>KLGKEAAAKAIKEWGQPKSKITHLIFCTTSGVDMPGADYQITKLLGLRPSVNR<br>FMMYQQGCFAGGTVLRLAKDLAENNRGARVLVVCSEITAVTFRGPSESHLDSL<br>VGQALFGDGAGAVIVGADPDLETERPLFELVSASQTILPDSEGAIDGHLREVGL<br>TFHLLKDVPGLISKNIEKSLVEAFAPLGIDDWNSIFWIAHPGGPAILDQVEAKLA<br>LEKEKMAATRQVLSEYGNMSSACVLFILDEMRRKSAEEGKATTGEGFNWGV<br>LFGFGPGLTVETVVLHSKPINY |
| Zo:_XP_042434663.1 curcumin synthase 1 [*Zingiber officinale*] | MASLHALRREQRAQGPATIMAIGTATPPNLYEQSTFPDFYFRVTNSDDKQELKE<br>KFRRMCNKSMVKKRYLYLTEEILKERPKLCSYKEPSFDDRQDIVVEEIPKLAKE<br>AAEKAIKEWGRPKSEITHLVFCSISGIDMPGADYRLATLLGLPLTVNRLMIYSQ<br>ACHMGAAMLRIAKDLAENNRGARVLVVACEITVLSFRGPNERDFEALAGQAG<br>FADGAAAVVVGADPLERVEKPIYEIAAAMQETVAESQEAVGGHLRAFGWTFY<br>FLNQLPAIISNNIGKSLERALVPLGVREWNDVFWVAHPGNWAIMDAIEAKLQL<br>TPDKLSTARHVFSEYGNMQSATVYFVMDELRKRSAVEGRSTTGDGLQWGVLF<br>GFGPGLSIETVVLRSMPL |
| Zo:_XP_042434575.1 chalcone synthase 2 [*Zingiber officinale*] | MAKVQEIRRSQRAEGPATVLAIGTATPANVVYQADYADYYFRMTKSEHLTELK<br>EKFKRMCDKSMIRKRYMHLTEEMLSENPNMCAYMEPSLDERQDIVVVEVPKL<br>GKEAAAKAIKEWGQPKSKITHLIFCTTSGVDMPGADYQITKLLGLRPSVNRFM<br>MYQQGCFAGGTVLRLAKDLAENNRGARVLVVCSEITAVTFRGPSESHLDSLVG<br>QALFGDGAGAVIVGADPDLETERPLFELVSASQTILPDSEGAIDGHLREVGLTF<br>HLLKDVPGLISKNIEKSLVEAFAPLGIDDWNSIFWIAHPGGPAILDQVEAKLALE<br>KDKMAATRQVLSEYGNMSSACVLFILDEMRRKSAEEGKATTGEGFNWGVLF<br>GFGPGLTVETVVLHSKPINY |
| Zo:_XP_042431535.1 phenylpropanoylacetylCoA synthase like [*Zingiber officinale*] | MACTEAFRRAPPADGPATVLAIGTANPSHFVDQMQYPDYYFRVTNAEDKTEL<br>KQKFKRICEKSTIRKRHMCLTEEILKENPSLCAYMAPSFDARQGIVLEEVPRLA<br>KEAADKAIKEWGRPVSDVTHLVFCSAAGVDLPGVDYRLIQLLGLPARVRRVM<br>LYNVGCHAGGTALRVAKDLAENNKGARVLVVCSELNVMFFRGPDDHHFENLI<br>GQALFGDGAAALIVGADPEEAERAIYEVASATQVMLPESEEMVGGHLREIGLT<br>FHLASKLPAVVGGNIERCLEAAFGPQAGVADWNELFWIVHPGGRAIIDQVEAR<br>AGLTAEKLAVTRHVLREYGNMQSASVLFIMDEMRKRSAAEGCATTGQGCQW<br>GVLFGFGPGLTVETVVLRSVPIKLIN |
| Zo:_XP_042429688.1 curcumin synthase 3 like [*Zingiber officinale*] | MASINNVVVDAFPKPQRARGPATIMAIGTANPPNLYEQSAYPDFYFRVTNSDH<br>KPELKQKFRRLCERTMIKKRYMHLTEELLKEKPGMCSYMDTSFDERQDIVVE<br>EVPRLAKEAAVKAIKEWGRSKSEITHLVFCSTSGVDMPGADYRLANLLGLSSS<br>VNRIMLYNQACHIGAQTLRIAKDIAENNRTARVLVVACEVNTLIFRGPEERDFQ<br>NLVAQVAFGDGAAAVVVGADPVEGVESPIFEIMAALPFTVPETQMAVGGQLK<br>QIGLTFHVAHQLPGLIANNLETCLGEAMKPLGISDWNDVFWVAHPGNWGIMD<br>AVEAKLGLEQGKLQSSRHVFSEFGNMMSATVLFVMDEVRKRAAAKGAATTG<br>DGLQWGVLCGFGPGLSIETLVLRSVPL |
| Zo:_XP_042429686.1_PKS2 curcumin synthase 3 like [*Zingiber officinale*] | MASINNVVVDAFPKPQRARGPATVMAIGTANPPNLYEQSSYPDFYFRVTNSDH<br>KPELKQKFRRLCERSMIKKRYMHLTEELLKEKPGMCSYMDTSFDERQDIVVE<br>EVPRLAKEAAVKAIKEWGRSKSEITHLVFCSTSGVDMPGADYRLANLLGLSSS<br>VNRIMLYNQACHIGAQTLRIAKDIAENNRTARVLVVACEVNTLIFRGPEERDFQ<br>SLAAQVAFGDGAAAVVVGADPVEGVERPIFEIMAALPFTVPETQMAVGGQLK<br>QIGLTFHFAHQLPGLIANNLETCLGEALKPLGISDWNDVFWVAHPGNWGIMD<br>AVEAKLGLEQGKLQSSRHVFSEFGNMMSATVLFVMDEVRKRAAAKGAATTG<br>DGLQWGVLCGFGPGLSIETLVLRSVPL |

| | |
|---|---|
| Zo:_XP_042425457.1_DCS2 phenylpropanoylacetylCoA synthase like [*Zingiber officinale*] | MASTEAFRRAPPADGPATVLAIGTANPSHFVDQMQYPDYYFRVTNAEDKTEL<br>KQKFKRICEKSTIRKRHMCLTEEILKENPSLCAYMAPSFDARQGIVLEEVPRLA<br>KEAADKAIKEWGRPVSDVTHLVFCSAAGVDLPGVDYRLIQLLGLPARVRRVM<br>LYNVGCHAGGTALRVAKDLAENNKGARVLVVCSELNVMFFRGPDDHHFENLI<br>GQALFGDGAAALIVGADPEEAERAIYEVASATQVMLPESEEMVGGHLREIGLT<br>FHLASKLPAVVGGNIERCLEAAFGPQAGVADWNELFWIVHPGGRAIIDQVEAR<br>AGLTAEKLAVTRHVLREYGNMQSASVLFIMDEMRKRSAAEGCATTGQGCQW<br>GVLFGFGPGLTVETVVLRSVPIKLIN |
| Zo:_XP_042422640.1 curcumin synthase 3 like [*Zingiber officinale*] | MASINNAVVDAFPKPQRARGPATVMAIGTANPPNLYEQSAYPDFYFRVTNSDH<br>KPELKQKFRRLCERSMIKKRYMHLTEELLKEKPGMCSYMDTSFDERQDIVVE<br>EVPRLAKEAAVNTIKEWGRSKSEITHLVFCSTSGVDMPGADYRLANLLGLSSS<br>VNRIMLYNQACHIGAQTLRIAKDIAENNRAARVLVVACEVNTLIFRGPEERDFQ<br>SLAAQVAFGDGAAAIVVGADPVEGVERPIFEIMAALPFTVPETQMAVGGQLK<br>QIGLTFHFAHQLPGLIANNLETCLGEALKPLGISDWNDVFWVAHPGNWGIMD<br>AVEAKLGLEQGKLQSSRHVFSEFGNMMSATVLFVMDDVRKRAAAKGAATTG<br>DGLQWGVLCGFGPGLSIETLVLRSVPL |
| Zo:_XP_042397679.1 phenylpropanoylacetylCoA synthase [*Zingiber officinale*] | MEVNGYRIIHSADGPATILAIGTANPTNVVDQNAYPDFYFRVTNSEHLQELKAK<br>FRRICEKAAIRKRHLYLTEEILREHPSLLAPMAPSFDARQQIVVEAVPKLAKEA<br>AEKAIKEWGRPKSDITHLVFCSASGIDMPGSDLQLLKLLGLPLCVNRVMLYNV<br>GCHAGGTALRVAKDLAENNRGARVLAVCSEVTVLSYRGPHPAHIESLFVQALF<br>GDGAAALVVGSDPVDGVERPIFEIASASQVMLPESEEAVGGHLREIGLTFHLKS<br>QLPSIIASNIEQSLTTACSPLGLSDWNQLFWTVHPGGRAILDQVEARLGLEKDR<br>LAATRHVLSEYGNMQSATVLFILDEMRKRSAADGHATTGEGLDWGVLLGFGP<br>GLSIETVVLHSCKLN |
| Zo:_XP_042397481.1 phenylpropanoylacetylCoA synthase like [*Zingiber officinale*] | MAAVMEAFSRTPPADGAANVLAIGTANPSHFVDQMQYPEYYFRITDAEGKTE<br>LQQKFKRICEKSMIRKRHMCLTEEVLRENPCLCGYMTPSFDARQRIVVEEVPR<br>LAKEAADKAINEWGRPVTDITHLVFCSAAGVDLHGADYRLLQLLGLPLHVRR<br>VMLYNVGCHAGGTALRVAKDLAENNKGARVLVVCSELNVMFFRGPGDDHIE<br>NLIGQALFGDGAAAVIVGADTDETERPIYEVASATQVMLPESEEMVGGHLREI<br>GLTFHLASRLPAVVGENIERCLESAFGVEAGDWNELFWIVHPGGRAIIDQVEA<br>RVRLTPEKLAATRHVLREYGNMQSASVLFIMDEMRKLSAAEGCATTGQGCQ<br>WGVLFGFGPGLTIETVVLRSVPRQIN |
| Zo:_XP_042395314.1 curcumin synthase 3 [*Zingiber officinale*] | MGSLQAMRRAKRAQGPATIMAVGTANPPNLYEQTSYPDFYFRVTNSDDKHEL<br>KNKFRVICEKTRVKRRYLHLTEEILKQRPKLCSYMEPSFDDRQDIVVDEIPKLA<br>KEAAEKAIKEWGRPKSEITHLVFCSISGIDMPGADYRLAKLLGLPLSVNRLML<br>YSQACHMGAQMLRIAKDLAENNRGARVLVVSCEITVLSFRGPDAGDFEALAC<br>QAGFGDGAAAVVVGADPLPGVERPIYEIAAAMQETVPESERAVGGHLREIGW<br>TFHFFNQLPKLIAENIESSLARAFKPLGITEWNDVFWVAHPGNWGIMDAIETKL<br>GLKQGKLATARHVFSEYGNMQSATVYFVMDEVRKRSAAEGRATTGEGLEWG<br>VLFGFGPGLTIETVVLRSLPIP |
| Zo:_XP_042392148.1_DCS1 phenylpropanoylacetylCoA synthase like [*Zingiber officinale*] | MAAVMEAFSRTPPADGAANVLAIGTANPSHFVDQMQYPEYYFRITDAEGKTE<br>LQQKFKRICEKSMIRKRHMCLTEEVLRENPCLCGYMTPSFDARQRIVVEEVPR<br>LAKEAADKAIKEWGHPVTDITHLVFCSAAGVDLPGADYSLLQLLGLPLHVRR<br>VMLYNVGCHAGGTALRVAKDLAENNKGARVLVVCSELNVMFFRGPGDDHIE<br>NLIGQALFGDGAAAVIVGADTDETERPIYEVASATQVMLPESEEMVGGHLREI<br>GLTFHLASRLPAVVGENIERCLESAFGVEAGDWNELFWIVHPGGRAIIDQVEA<br>RVRLRPEKLAATRHVLREYGNMQSASVLFIMDEMRKLSAAEGCATTGQGCQ<br>WGVLFGFGPGLTIETVVLRSVPRQIN |
| Zo:_XP_042390634.1_PKS1 curcumin synthase 3 like [*Zingiber officinale*] | MGSLQAMRRAKRAQGPATIMAVGTANPPNLYEQTSYPDFYFRVTNSDDKHEL<br>KNKFRVICEKTRVKRRYLHLTEEILKQRPKLCSYMEPSFDDRQDIVVDEIPKLA<br>KEAAEKAIKEWGHPKSEITHLVFCSISGIDMPGADYRLAKLLGLPLSVNRLML<br>YSQACHMGAQMLRIAKDLAENNRGARVLAVSCEITVLSFRGPDAGDFEALAC<br>QAGFGDGAAAVVVGADPLPGVERPIYEIAAAMQETVPESERAVGGHLREIGW<br>TFHFFNQLPKLIAENIQSSLARAFKPLGITEWNDVFWVAHPGNWGIMDAIETK<br>LGLKQGKLATARHVFSEYGNMQSATVYFVMDEVRKRSAAEGRATTGEGLEW<br>GVLFGFGPGLTIETVVLRSLPIP |

| | |
|---|---|
| Zo:_XP_042378779.1<br>chalcone synthase 2 like<br>[*Zingiber officinale*] | MAKVQEIRRSQRAEGPAAVLAIGTATPANVVYQADYADYYFRMTKSEHLTEL<br>KAKFKRMCDKSMIRKRYMHLTEEMLSENPNMCAYMEPSLDERQDIAVVEVP<br>KLGKEAAAKAIKEWGQPESKITHLIFCTTSGVDMPGADYQITKLLGLRPSVNR<br>FMMYQQGCFAGGTVLRLAKDLAENNRGARVLVVCSEITAVTFRGPSESRLDN<br>LVGQALFGDGAGAVIVGADPDLETERPLFELVSASQTILPDSEGAIGGHLREVG<br>LTFHLLKDVPGLISKNIEKSLVEAFAPLGIDDWNSIFWIAHPGGPAILDQFEAKL<br>ALEKEKMTATRQVLSEYGNMSSACVLFILDEMRRKSAEEGKATTGEGFNWGV<br>LFGFGPGITVETVVLHSKPINY |
| Zo:_XP_042373730.1<br>chalcone synthase 2 like<br>[*Zingiber officinale*] | MAKLVVSEIRRSQRAEGAAAVLAIGTANPPNVVYQADYPDYYFRITRSEHLTE<br>LKEKFKRMCDKSMIRKRHMYLTEEILKENPKMCAYMEPSLDARQDIVVLEVP<br>KMGKEAAAKAIKEWGQPKSKITHLVFCTTSGVDMPGADYQLTKLLGLRPSVN<br>RFMMYQQGCFAGGTVLRLAKDLAENNRGARVLVVCSEITAVTFRGPSDSHLD<br>SMVGQALFADGAGAIIVGADPDPATERPLFELVSASQTILPDSEGAIDGHLREV<br>GLTFHLLKDVPGLISKNIEKSLAEAFKPLGISDWNSLFWIAHPGGPAILDQVEA<br>KLALDKNKMKATRDVLSEYGNMSSACVLFILDEMRRRSVEEGKATTGEGLE<br>WGVLFGFGPGLTVETVVLHSVPIAAAH |
| Os:_Q8LIL0.2_CUS<br>[*Oryza sativa*] | MAPTTTMGSALYPLGEMRRSQRADGLAAVLAIGTANPPNCVTQEEFPDFYFRV<br>TNSDHLTALKDKFKRICQEMGVQRRYLHHTEEMLSAHPEFVDRDAPSLDARL<br>DIAADAVPELAAEAAKKAIAEWGRPAADITHLVVTTNSGAHVPGVDFRLVPLL<br>GLRPSVRRTMLHLNGCFAGCAALRLAKDLAENSRGARVLVVAAELTLMYFTG<br>PDEGCFRTLLVQGLFGDGAAAVIVGADADDVERPLFEIVSAAQTIIPESDHALN<br>MRFTERRLDGVLGRQVPGLIGDNVERCLLDMFGPLLGGDGGGGWNDLFWAV<br>HPGSSTIMDQVDAALGLEPGKLAASRRVLSDYGNMSGATVIFALDELRRQRKE<br>AAAAGEWPELGVMMAFGPGMTVDAMLLHATSHVN |
| Cl:_BAH85781.1_CURS3<br>curcumin synthase<br>[*Curcuma longa*] | MGSLQAMRRAQRAQGPATIMAVGTSNPPNLYEQTSYPDFYFRVTNSDHKHAL<br>KNKFRVICEKTKVKRRYLHLTEEILKQRPKLCSYMEPSFDDRQDIVVEEIPKLA<br>KEAAEKAIKEWGRPKSEITHLVFCSISGIDMPGADYRLATLLGLPLSVNRLMLY<br>SQACHMGAQMLRIAKDLAENNRGARVLAVSCEITVLSFRGPDAGDFEALACQ<br>AGFGDGAAAVVVGADPLPGVERPIYEIAAAMQETVPESERAVGGHLREIGWT<br>FHFFNQLPKLIAENIEGSLARAFKPLGISEWNDVFWVAHPGNWGIMDAIETKL<br>GLEQGKLATARHVFSEYGNMQSATVYFVMDEVRKRSAAEGRATTGEGLEWG<br>VLFGFGPGLTIETVVLRSVPLP |
| Cl:_BAH85780.1_CURS2<br>curcumin synthase<br>[*Curcuma longa*] | MAMISLQAMRKAQRAQGPATILAVGTANPPNLYEQDTYPDYYFRVTNSEHKQ<br>ELKNKFRLMCEKTMVKRRYLYLTPEILKERPKLCSYMEPSFDDRQDIVVEEVP<br>KLAAEAAENAIKEWGGDKSAITHLVFCSISGIDMPGADYRLAQLLGLPLAVNR<br>LMLYSQACHMGAAMLRIAKDIAENNRSARVLVVACEITVLSFRGPDERDFQAL<br>AGQAGFGDGAGAMIVGADPVLGVERPLYHIMSATQTTVPESEKAVGGHLREV<br>GLTFHFFNQLPAIIADNVGNSLAEAFEPIGIKDWNNIFWVAHPGNWAIMDAIET<br>KLGLEQSKLATARHVFSEFGNMQSATVYFVMDELRKRSAAENRATTGDGLRW<br>GVLFGFGPGISIETVVLQSVPL |
| Cl:_BAH56226.1_CURS1<br>curcumin synthase<br>[*Curcuma longa*] | MANLHALRREQRAQGPATIMAIGTATPPNLYEQSTFPDFYFRVTNSDDKQELK<br>KKFRRMCEKTMVKKRYLHLTEEILKERPKLCSYKEASFDDRQDIVVEEIPRLA<br>KEAAEKAIKEWGRPKSEITHLVFCSISGIDMPGADYRLATLLGLPLTVNRLMIY<br>SQACHMGAAMLRIAKDLAENNRGARVLVVACEITVLSFRGPNEGDFEALAGQ<br>AGFGDGAGAVVVGADPLEGIEKPIYEIAAAMQETVAESQGAVGGHLRAFGWT<br>FYFLNQLPAIIADNLGRSLERALAPLGVREWNDVFWVAHPGNWAIIDAIEAKL<br>QLSPDKLSTARHVFTEYGNMQSATVYFVMDELRKRSAVEGRSTTGDGLQWG<br>VLLGFGPGLSIETVVLRSMPL |
| Cl:_BAH56225.1_DCS<br>diketide CoA synthase<br>[*Curcuma longa*] | MEANGYRITHSADGPATILAIGTANPTNVVDQNAYPDFYFRVTNSEYLQELKA<br>KFRRICEKAAIRKRHLYLTEEILRENPSLLAPMAPSFDARQAIVVEAVPKLAKE<br>AAEKAIKEWGRPKSDITHLVFCSASGIDMPGSDLQLLKLLGLPPSVNRVMLYN<br>VGCHAGGTALRVAKDLAENNRGARVLAVCSEVTVLSYRGPHPAHIESLFVQAL<br>FGDGAAALVVGSDPVDGVERPIFEIASASQVMLPESAEAVGGHLREIGLTFHLK<br>SQLPSIIASNIEQSLTTACSPLGLSDWNQLFWAVHPGGRAILDQVEARLGLEKD<br>RLAATRHVLSEYGNMQSATVLFILDEMRNRSAAEGHATTGEGLDWGVLLGFG<br>PGLSIETVVLHSCRLN |

| | |
|---|---|
| At:_AAF23561.1_CHS<br>[*Arabidopsis thaliana*] | MVMAGASSLDEIRQAQRADGPAGILAIGTANPENHVLQAEYPDYYFRITNSEH<br>MTDLKEKFKRMCDKSTIRKRHMHLTEEFLKENPHMCAYMAPSLDTRQDIVVV<br>EVPKLGKEAAVKAIKEWGQPKSKITHVVFCTTSGVDMPGADYQLTKLLGLRP<br>SVKRLMMYQQGCFAGGTVLRIAKDLAENNRGARVLVVCSEITAVTFRGPSDT<br>HLDSLVGQALFSDGAAALIVGSDPDTSVGEKPIFEMVSAAQTILPDSDGAIDGH<br>LREVGLTFHLLKDVPGLISKNIVKSLDEAFKPLGISDWNSLFWIAHPGGPAILD<br>QVEIKLGLKEEKMRATRHVLSEYGNMSSACVLFILDEMRRKSAKDGVATTGE<br>GLEWGVLFGFGPGLTVETVVLHSVPL |

**Table S2: Representative mass spectra of major compound peaks identified by HPLC analysis of recombinant *E. coli* culture extracts shown in Figures 4-6.**

| Figure | Peak | Exact Mass | Observed $[M-H]^-$ | Compound |
|---|---|---|---|---|
| 4 & 5 | 3 | 368.13 | 367.12 | Curcumin (standard) |

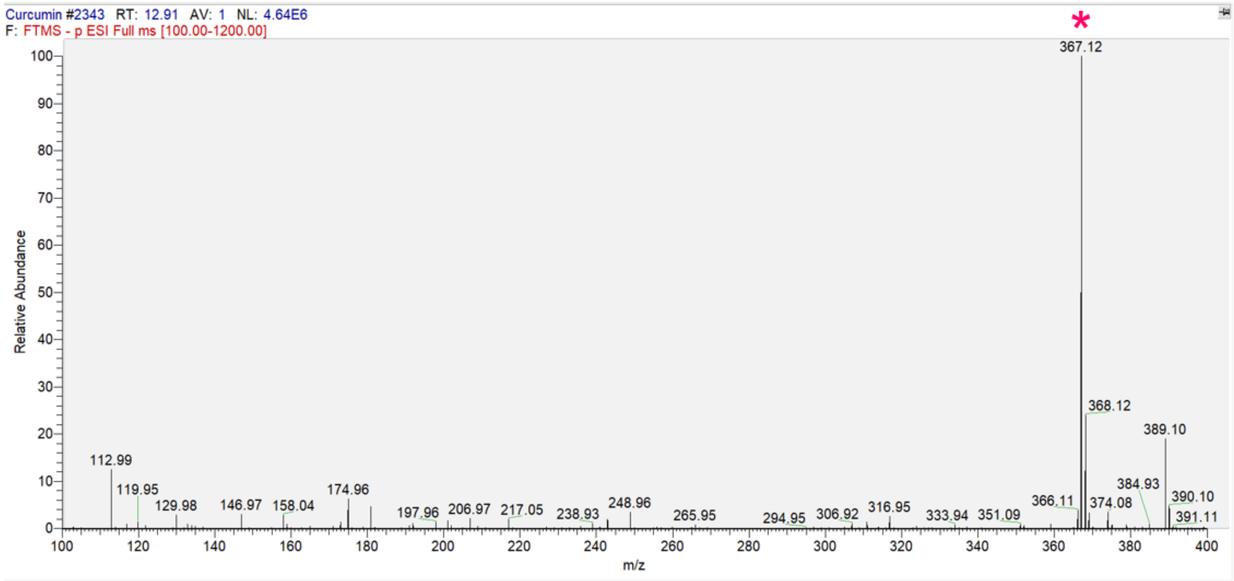

| Figure | Peak | Exact Mass | Observed $[M-H]^-$ | Compound |
|---|---|---|---|---|
| 4 & 5 | 4 | 290.15 | 289.14 | 6-DHG (standard) |

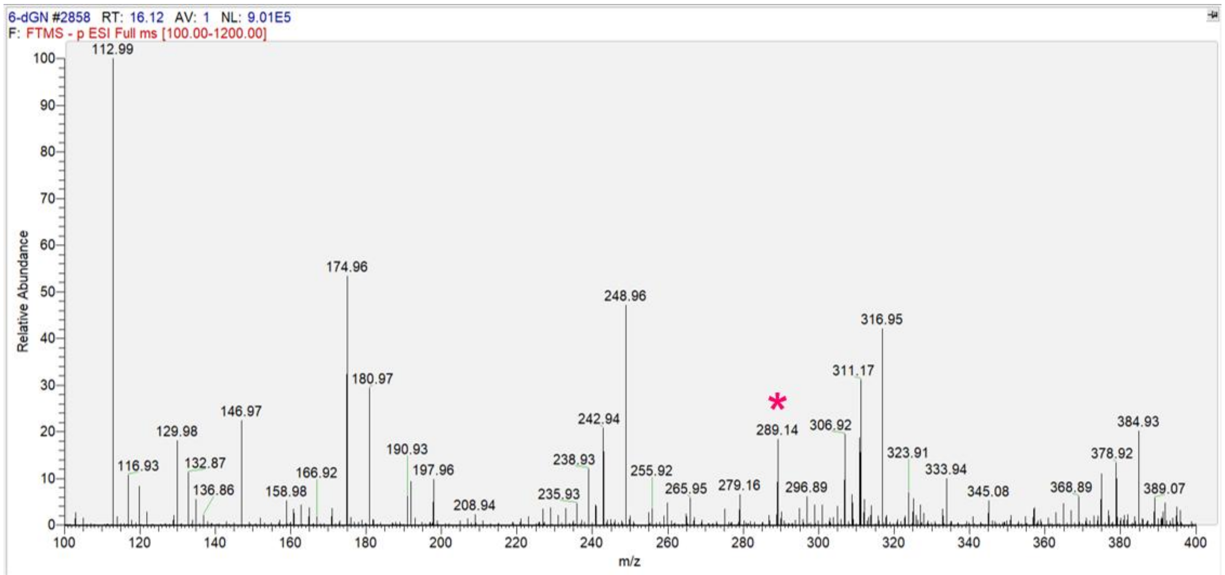

| Figure | Peak | Exact Mass | Observed $[M-H]^-$ | Compound |
|---|---|---|---|---|
| 4 | 3 | 368.13 | 367.12 | Curcumin |

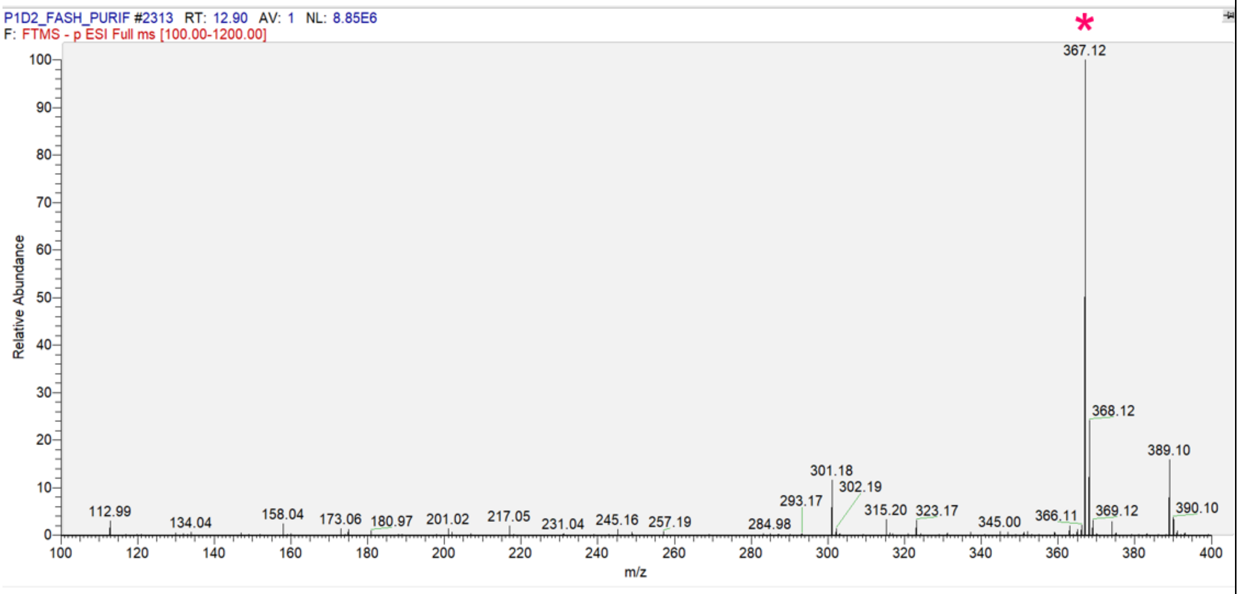

| Figure | Peak | Exact Mass | Observed $[M-H]^-$ | Compound |
|---|---|---|---|---|
| 4 | 4 | 290.15 | 289.14 | 6-DHG |

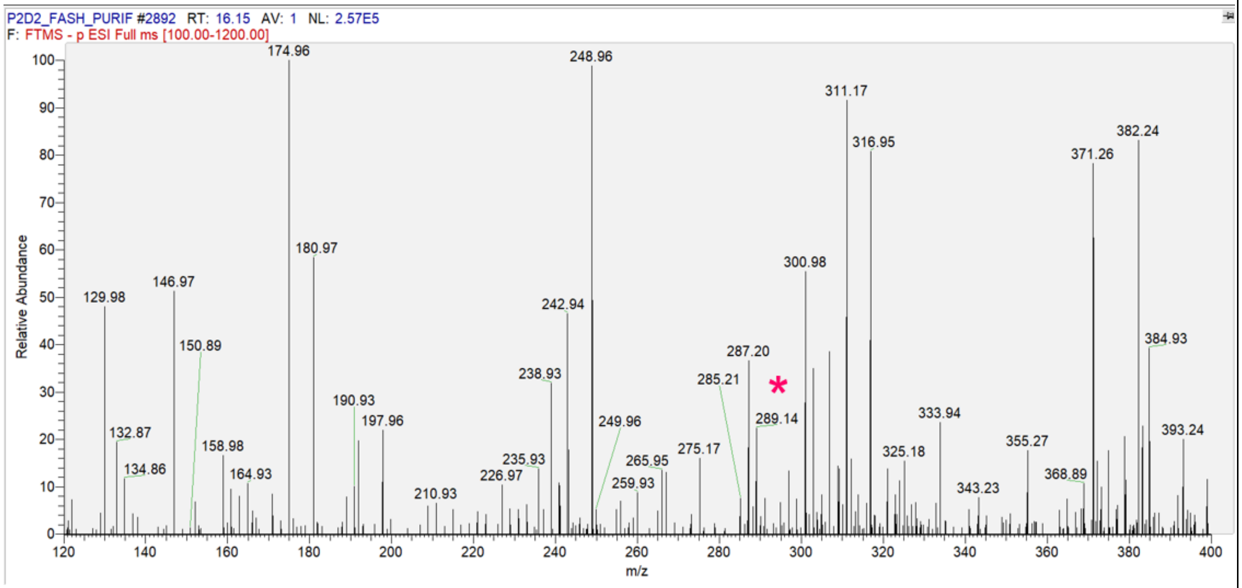

| Figure | Peak | Exact Mass | Observed [M-H]- | Compound |
|---|---|---|---|---|
| 5 | 2a (*Zo*PKS1) | 234.09 | 233.12 | Feruloylacetone |

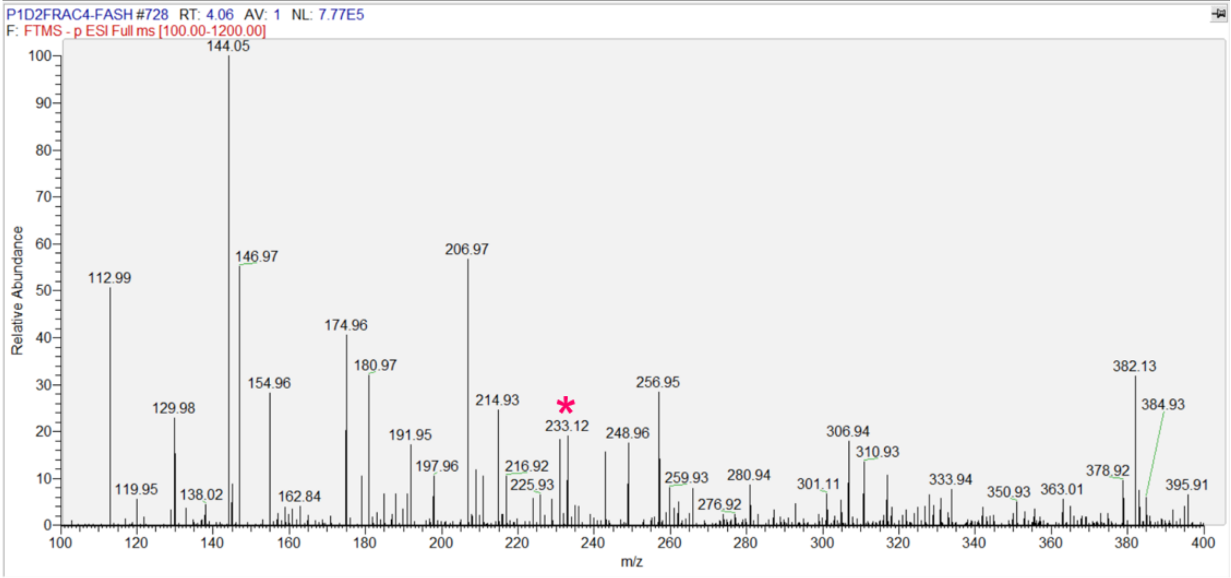

| Figure | Peak | Exact Mass | Observed [M-H]- | Compound |
|---|---|---|---|---|
| 5 | 2b (*Zo*PKS2) | 192.08 | 191.07 | Feruloylmethane (Dehydrozingerone) |

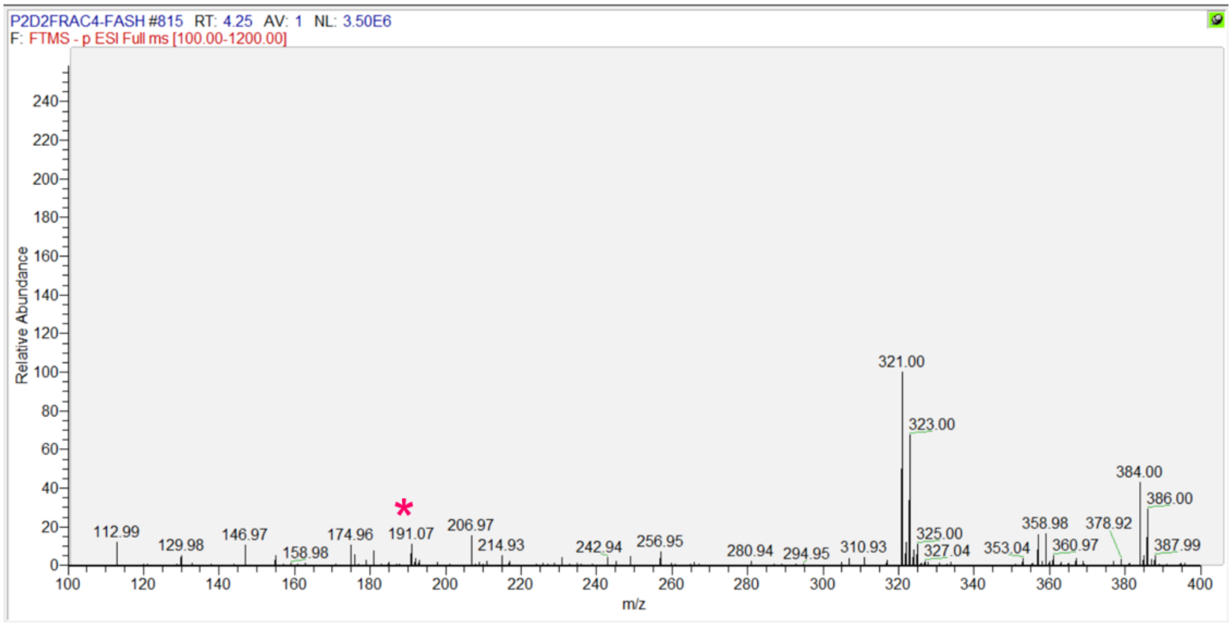

| Figure | Peak | Exact Mass | Observed $[M-H]^-$ | Compound |
|---|---|---|---|---|
| 6 | 3a | 308.10 | 307.10 | Bisdemethoxycurcumin |

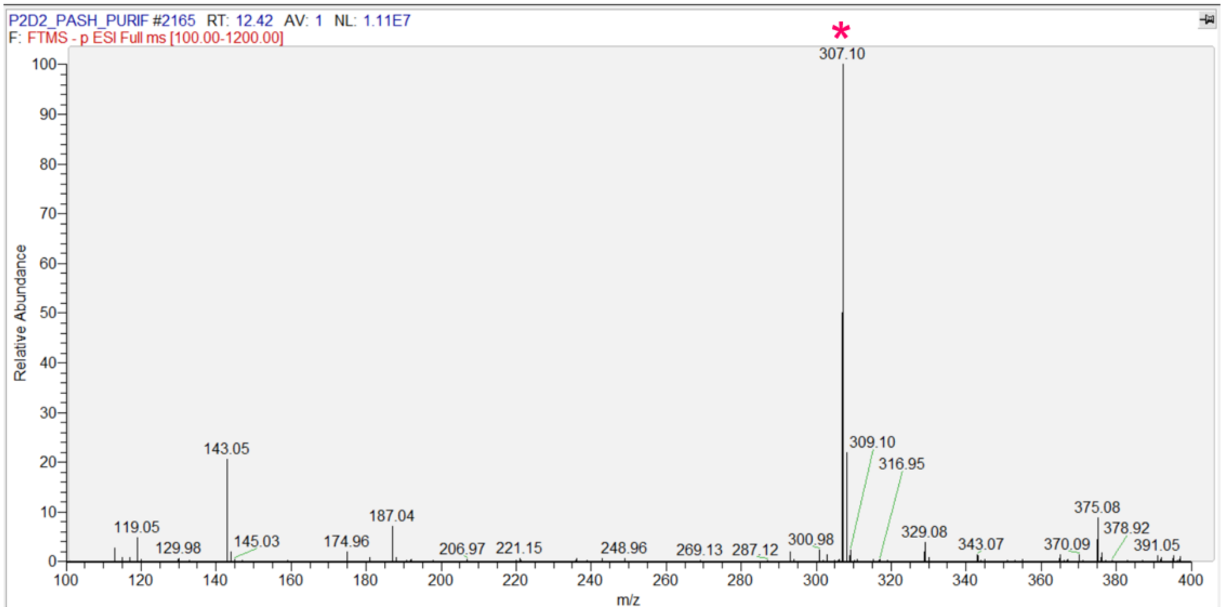

| Figure | Peak | Exact Mass | Observed $[M-H]^-$ | Compound |
|---|---|---|---|---|
| 6 | 3b | 276.12 | 276.16 | Dicinnamoylmethane (protonated) |

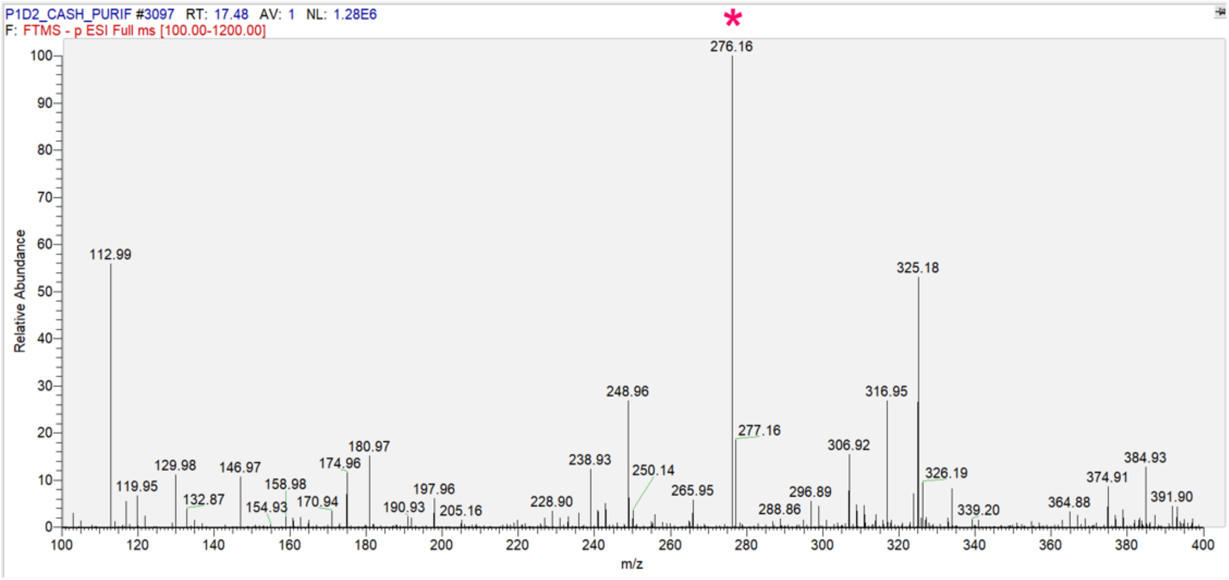

| Figure | Peak | Exact Mass | Observed [M-H]⁻ | Compound |
|---|---|---|---|---|
| 6 | 4a | 260.14 | 259.13 | *p*-Coumaroylhexanoylmethane |

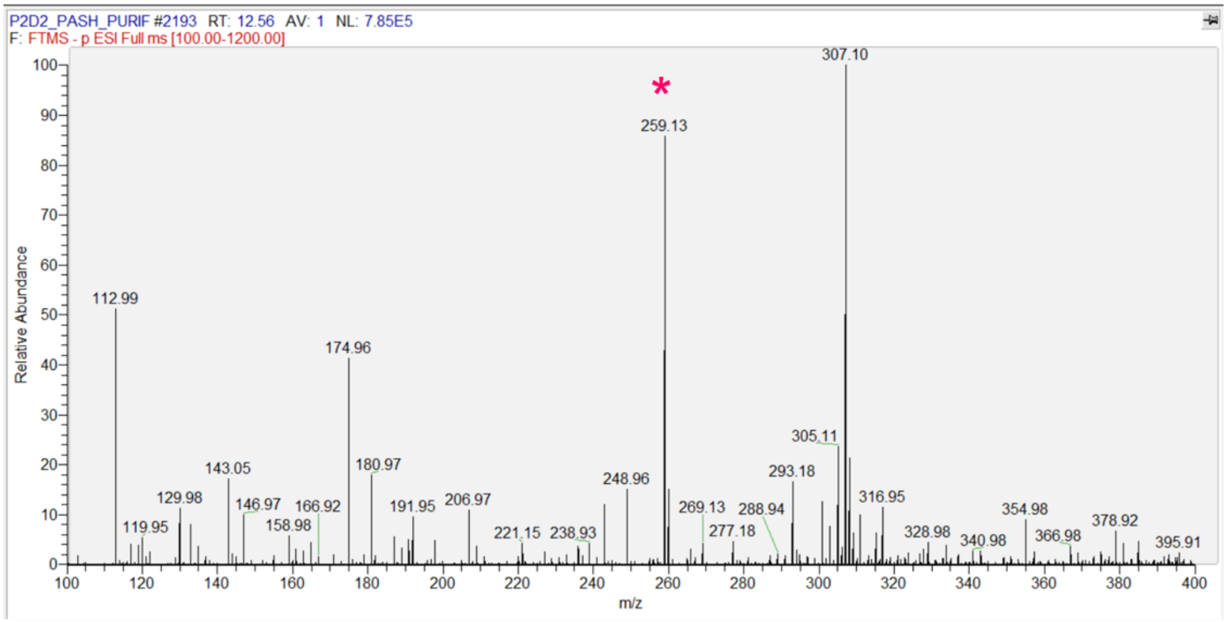

| Figure | Peak | Exact Mass | Observed [M-H]⁻ | Compound |
|---|---|---|---|---|
| 6 | 4c | 228.15 | 227.20 | Phenyldeca-1,4-dien-3-one |

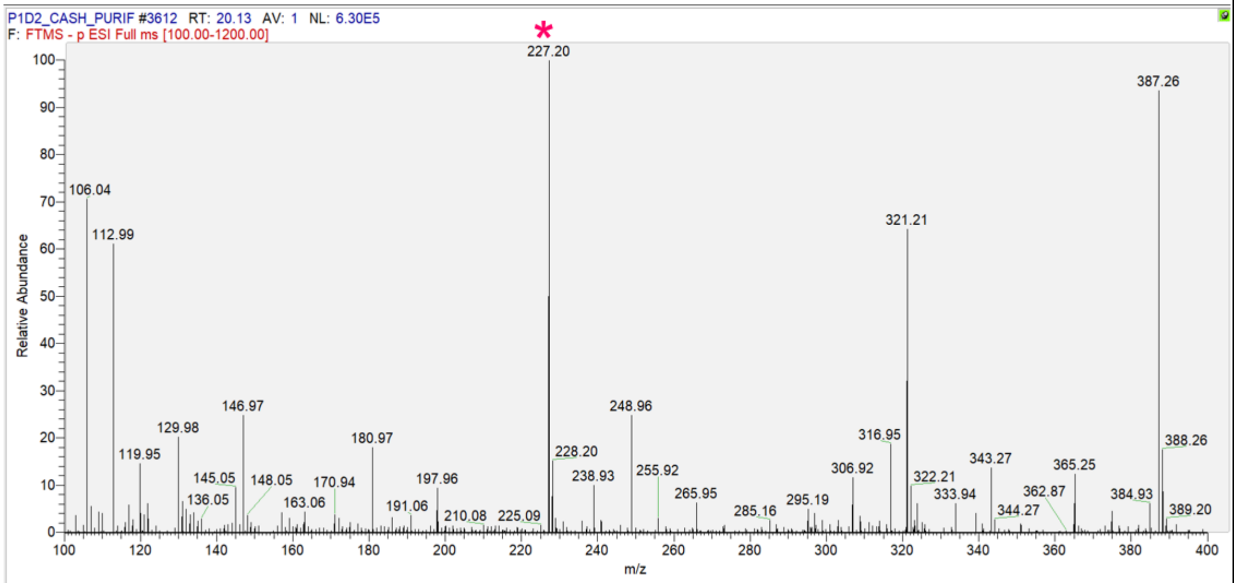

| Figure | Peak | Exact Mass | Observed [M-H]$^-$ | Compound |
|---|---|---|---|---|
| 6 | 6a | 206.06 | 205.05 | *p*-Coumaroyl-ß-ketoacid |

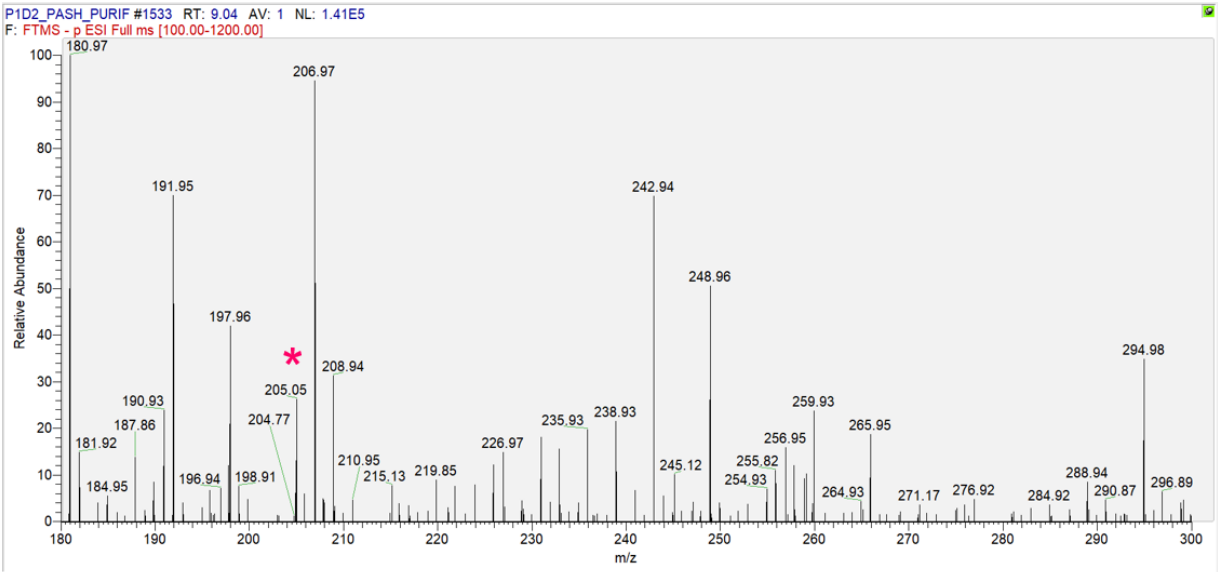

| Figure | Peak | Exact Mass | Observed [M-H]$^-$ | Compound |
|---|---|---|---|---|
| 6 | 6b | 162.07 | 161.04 | *p*-Hydroxybenzalacetone (*p*-Coumaroylmethane) |

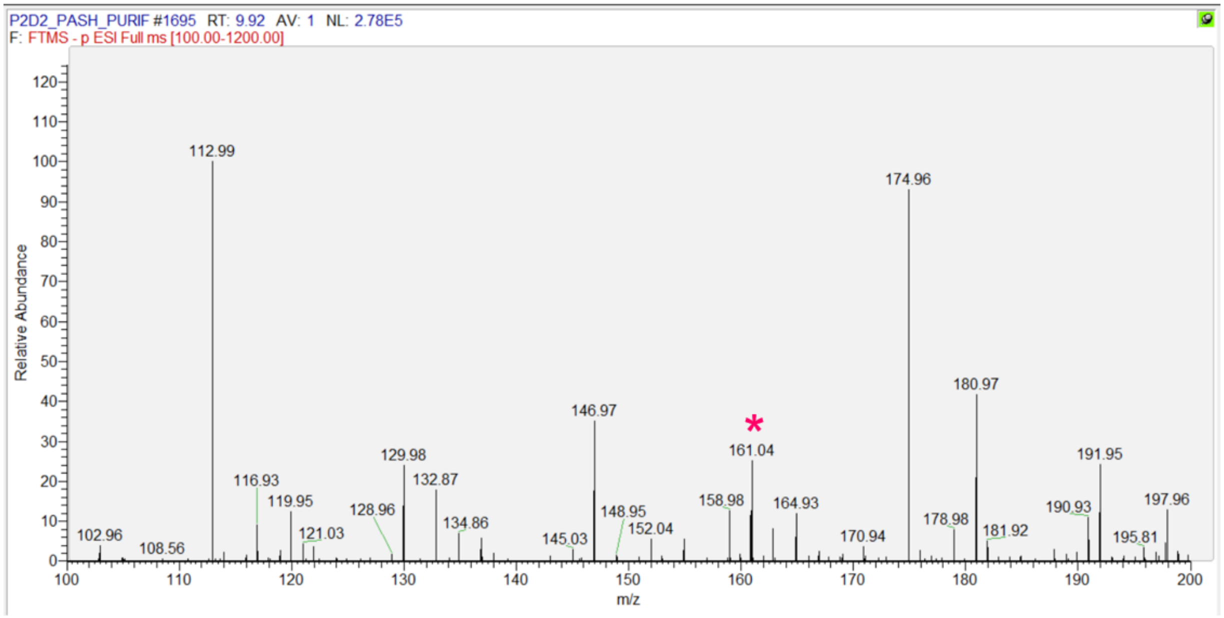

| Figure | Peak | Exact Mass | Observed [M-H]$^-$ | Compound |
|---|---|---|---|---|
| 6 | * | NA | 931.13, 841.16, 465.08, 419.08 | Unknown Peptide? |

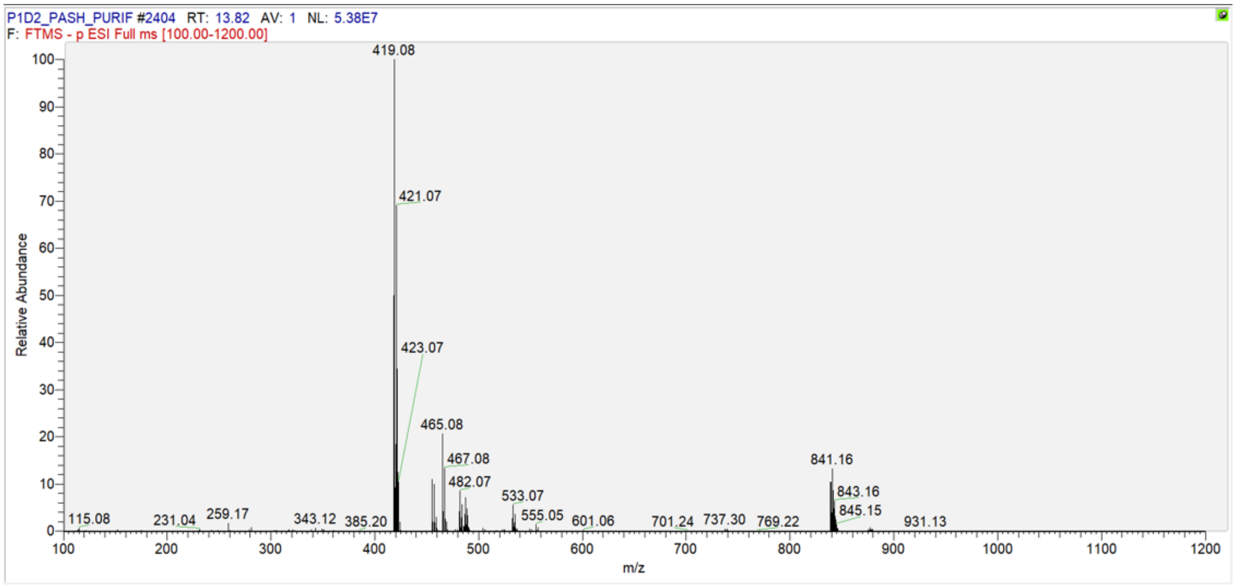

| Figure | Peak | Exact Mass | Observed [M-H]$^-$ | Compound |
|---|---|---|---|---|
| 6 | ? | NA | 213.96, 298.94, 174.96 | Unknown |

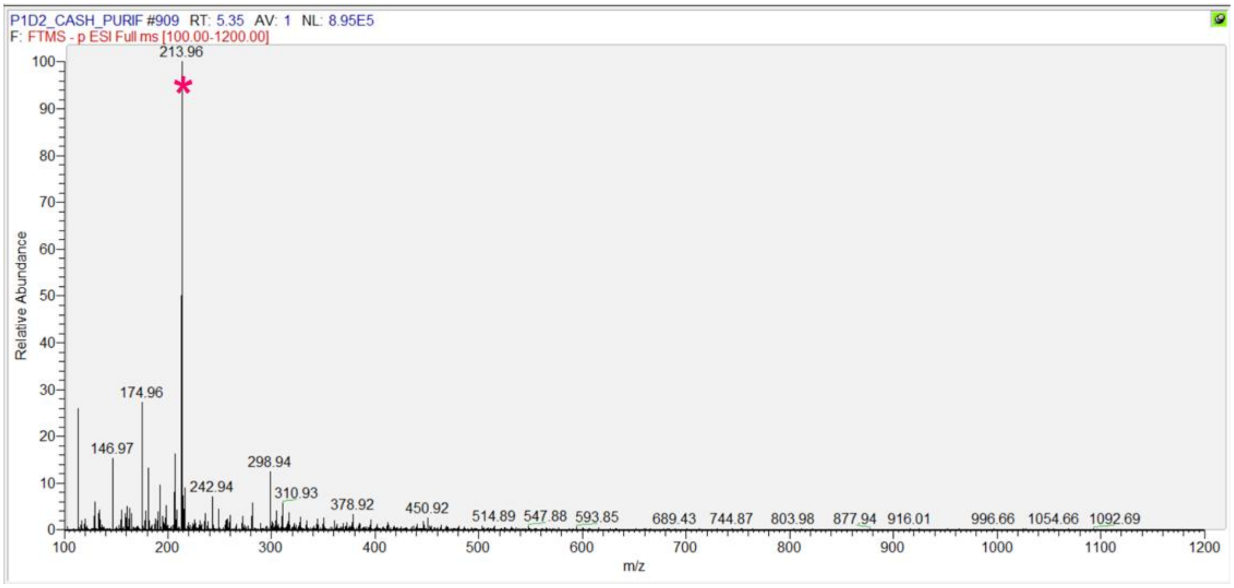

**Table S3: Plasmids and strains used in this study.**

| Plasmids | Description | Source |
|---|---|---|
| $pAC_{mod}$ - Empty | Constitutive $Lac_{mod}$ promoter, $Cm^R$ | [1] |
| $pAC_{mod}$-4CL1 | Constitutive expression vector for 4CL1, $Cm^R$ | [2] |
| $pAC_{mod}$-$His_6$-*At*4Cl1 | Derived from $pAC_{mod}$ : *At*4Cl1 with 6xHis-tag and thrombin cleavage site | This study |
| $pAC_{mod}$-$His_6$-*At*$ACS_{mut}$ | Derived from $pAC_{mod}$ : *At*$ACS_{T324G, V399A, W427G}$ with 6xHis-tag and thrombin cleavage site | This study |
| $pAC_{mod}$-$His_6$-*At*$ACS_{mut}$ + $His_6$-*At*4CL1 | Derived from $pAC_{mod}$-$His_6$- *At*$ACS_{T324G, V399A, W427G}$ : *At*4Cl1 cloned with $Lac_{mod}$ promoter, 6xHis-tag, and thrombin cleavage site cloned downstream with 160nt spacer between terminus and $Lac_{mod}$ promoter. | This study |
| pET28a-Empty | T7 promoter, $Km^R$ (6xHis-tag and thrombin site downstream of MCS in pET28a) | Invitrogen |
| pET28a-$His_6$-*Zo*PKS1 | Derived from pET28a: *Zo*PKS1 with His tag and thrombin cleavage site | This study |
| pET28a-$His_6$-*Zo*PKS2 | Derived from pET28a: *Zo*PKS2 with His tag and thrombin cleavage site | This study |
| pET28a-$His_6$-*Zo*DCS1 | Derived from pET28a: *Zo*DCS1 with His tag and thrombin cleavage site | This study |
| pET28a-$His_6$-*Zo*DCS2 | Derived from pET28a: *Zo*DCS2 with His tag and thrombin cleavage site | This study |
| pET28a-$His_6$-*Zo*PKS1 + $His_6$-*Zo*DCS1 | Derived from pET28a-$His_6$-*Zo*PKS1: ZoDCS1 with T7 promoter, 6xHis-tag, and thrombin cleavage site | This study |
| pET28a-$His_6$-*Zo*PKS1 + $His_6$-*Zo*DCS2 | Derived from pET28a-$His_6$-*Zo*PKS1: ZoDCS2 with T7 promoter, 6xHis-tag, and thrombin cleavage site | This study |
| pET28a-$His_6$-*Zo*PKS2 + $His_6$-*Zo*DCS1 | Derived from pET28a-$His_6$-*Zo*PKS2: *Zo*DCS1 with T7 promoter, 6xHis-tag, and thrombin cleavage site | This study |
| pET28a-$His_6$-*Zo*PKS2 + $His_6$-*Zo*DCS2 | Derived from pET28a-$His_6$-*Zo*PKS2: ZoDCS2 with T7 promoter, 6xHis-tag, and thrombin cleavage site | This study |
| **Strains** | **Description** | **Source** |
| *Escherichia coli* Top 10 | General cloning host | Invitrogen |
| *Escherichia coli* BL21(DE3) | Protein expression and bioconversion experiments | New England Biolabs |

**Table S4. Primers used for molecular cloning.**

| | | |
|---|---|---|
| | **Gene Insert Primer Sequences** | |
| ***Zo*PKS1 Gene Insert** | | |
| | Forward (5’-3’): | ccgcgcggcagcATGGGCAGCCTG |
| | Reverse (5’-3’): | cgagtgcggccgcttaCGGAATCGGCAGG |
| ***Zo*PKS2 Gene Insert** | | |
| | Forward (5’-3’): | ccgcgcggcagcATGGCGAGC |
| | Reverse (5’-3’): | gtgcggccgcttaCAGCGGCACG |
| ***Zo*DCS1 Gene Insert** | | |
| | Forward (5’-3’): | gcgcggcagcATGGCTGCTGTGATGGAAG |
| | Reverse (5’-3’): | gagtgcggccgcttaGTTAATCTGACGCGGCAC |
| ***Zo*DCS2 Gene Insert** | | |
| | Forward (5’-3’): | cgcgcggcagcATGGCGAGCACAGAAGCTTTC |
| | Reverse (5’-3’): | ctcgagtgcggccgcttaGTTAATCAGTTTAATCGGCAC |
| ***At*ACSmut Gene Insert** | | |
| | Forward (5’-3’): | gcgcggcagcATGGCGAGCGAAGAAAA |
| | Reverse (5’-3’): | gtgcggccgcttaCACATCCGCCAGC |
| **$P_{T7}$-RBS-$His_6$-*Zo*PKS1 Gene Insert** | | |
| | Forward (5’-3’): | caaggaatggtgcatgcaaggagacaaaaaacccctcaagacccgtttagaggcc |
| | Reverse (5’-3’): | tatagtgagtcgtattatggcgcccaacagtccccc |
| **$P_{T7}$-RBS-$His_6$-*Zo*PKS2 Gene Insert** | | |
| | Forward (5’-3’): | caaggaatggtgcatgcaaggagacaaaaaacccctcaagacccgtttagaggcc |
| | Reverse (5’-3’): | tatagtgagtcgtattatggcgcccaacagtccccc |
| **$Lac_{mod}$-RBS-$His_6$-*At*4Cl1 Gene Insert** | | |
| | Forward (5’-3’): | catcaccatcaccatcactgatTGAGTTAGCTCACTCATTAGGCACCCCAG |
| | Reverse (5’-3’): | accgcacagatgcgtaaggagaAAATACCGCAGCGG |
| | **Vector Backbone Primer Sequences** | |
| **pET28a-$His_6$ Backbone** | | |
| | Forward (5’-3’): | taagcggccgcactcgagcac |
| | Reverse (5’-3’): | gctgccgcgcggcaccag |
| **$pAC_{mod}$ Backbone** | | |
| | Forward (5’-3’): | atgcatccatggcggccgcc |
| | Reverse (5’-3’): | ATGGATCCAGATCTCCTCCTTCTACTAGACGC |
| **Bi-directional pET28a-$His_6$-ZoDCS Backbone** | | |
| | Forward (5’-3’): | taatacgactcactataggggaattgtgag |
| | Reverse (5’-3’): | tctccttgcatgcaccattccttg |
| **pACmod-$His_6$-*At*ACSmut Backbone** | | |
| | Forward (5’-3’): | tcttattaatcagataaaataggcggccgccatggatgcat |
| | Reverse (5’-3’): | gagtgagctaactcacattataatgtgagttagctcactcattaggcacccc |

**Table S5: Amino acid sequences of proteins used in this study.**

| Name | Amino Acid Sequence |
|---|---|
| His$_6$-Thrombin-*Zo*PKS1 | **M**GSSHHHHHHSSGLVPRGSMGSLQAMRRAKRAQGPATIMAVGTANPPNLYEQTSYPDFYFRVT<br>NSDDKHELKNKFRVICEKTRVKRRYLHLTEEILKQRPKLCSYMEPSFDDRQDIVVDEIPKLAKE<br>AAEKAIKEWGHPKSEITHLVFCSISGIDMPGADYRLAKLLGLPLSVNRLMLYSQACHMGAQML<br>RIAKDLAENNRGARVLAVSCEITVLSFRGPDAGDFEALACQAGFGDGAAAVVVGADPLPGVER<br>PIYEIAAAMQETVPESERAVGGHLREIGWTFHFFNQLPKLIAENIQSSLARAFKPLGITEWNDVF<br>WVAHPGNWGIMDAIETKLGLKQGKLATARHVFSEYGNMQSATVYFVMDEVRKRSAAEGRAT<br>TGEGLEWGVLFGFGPGLTIETVVLRSLPIP |
| His$_6$-Thrombin-*Zo*PKS2 | **M**GSSHHHHHHSSGLVPRGSMASINNVVVDAFPKPQRARGPATVMAIGTANPPNLYEQSSYPDF<br>YFRVTNSDHKPELKQKFRRLCERSMIKKRYMHLTEELLKEKPGMCSYMDTSFDERQDIVVEEV<br>PRLAKEAAVKAIKEWGRSKSEITHLVFCSTSGVDMPGADYRLANLLGLSSSVNRIMLYNQACHI<br>GAQTLRIAKDIAENNRTARVLVVACEVNTLIFRGPEERDFQSLAAQVAFGDGAAAVVVGADPVE<br>GVERPIFEIMAALPFTVPETQMAVGGQLKQIGLTFHFAHQLPGLIANNLETCLGEALKPLGISDW<br>NDVFWVAHPGNWGIMDAVEAKLGLEQGKLQSSRHVFSEFGNMMSATVLFVMDEVRKRAAAK<br>GAATTGDGLQWGVLCGFGPGLSIETLVLRSVPL |
| His$_6$-Thrombin-*Zo*DCS1 | **M**GSSHHHHHHSSGLVPRGSMAAVMEAFSRTPPADGAANVLAIGTANPSHFVDQMQYPEYYFRI<br>TDAEGKTELQQKFKRICEKSMIRKRHMCLTEEVLRENPCLCGYMTPSFDARQRIVVEEVPRLAK<br>EAADKAIKEWGHPVTDITHLVFCSAAGVDLPGADYSLLQLLGLPLHVRRVMLYNVGCHAGGT<br>ALRVAKDLAENNKGARVLVVCSELNVMFFRGPGDDHIENLIGQALFGDGAAAVIVGADTDETE<br>RPIYEVASATQVMLPESEEMVGGHLREIGLTFHLASRLPAVVGENIERCLESAFGVEAGDWNELF<br>WIVHPGGRAIIDQVEARVRLRPEKLAATRHVLREYGNMQSASVLFIMDEMRKLSAAEGCATTG<br>QGCQWGVLFGFGPGLTIETVVLRSVPRQIN |
| His$_6$-Thrombin-*Zo*DCS2 | **M**GSSHHHHHHSSGLVPRGSMASTEAFRRAPPADGPATVLAIGTANPSHFVDQMQYPDYYFRVT<br>NAEDKTELKQKFKRICEKSTIRKRHMCLTEEILKENPSLCAYMAPSFDARQGIVLEEVPRLAKEA<br>ADKAIKEWGRPVSDVTHLVFCSAAGVDLPGVDYRLIQLLGLPARVRRVMLYNVGCHAGGTAL<br>RVAKDLAENNKGARVLVVCSELNVMFFRGPDDHHFENLIGQALFGDGAAALIVGADPEEAERA<br>IYEVASATQVMLPESEEMVGGHLREIGLTFHLASKLPAVVGGNIERCLEAAFGPQAGVADWNEL<br>FWIVHPGGRAIIDQVEARAGLTAEKLAVTRHVLREYGNMQSASVLFIMDEMRKRSAAEGCATT<br>GQGCQWGVLFGFGPGLTVETVVLRSVPIKLIN |
| His$_6$-Thrombin-*At*4Cl1 | **M**HHHHHHSSGLVPRGSHMMGSGGSGVDMEEDYKMAPQEQAVSQVMEKQSNNNNSDVIFRS<br>KLPDIYIPNHLSLHDYIFQNISEFATKPCLINGPTGHVYTYSDVHVISRQIAANFHKLGVNQNDV<br>VMLLLPNCPEFVLSFLAASFRGATATAANPFFTPAEIAKQAKASNTKLIITEARYVDKIKPLQNDD<br>GVVIVCIDDNESVPIPEGCLRFTELTQSTTEASEVIDSVEISPDDVVALPYSSGTTGLPKGVMLTH<br>KGLVTSVAQQVDGENPNLYFHSDDVILCVLPMFHIYALNSIMLCGLRVGAAILIMPKFEINLLLE<br>LIQRCKVTVAPMVPPIVLAIAKSSETEKYDLSSIRVVKSGAAPLGKELEDAVNAKFPNAKLGQG<br>YGMTEAGPVLAMSLGFAKEPFPVKSGACGTVVRNAEMKIVDPDTGDSLSRNQPGEICIRGHQI<br>MKGYLNNPAATAETIDKDGWLHTGDIGLIDDDDELFIVDRLKELIKYKGFQVAPAELEALLIGHP<br>DITDVAVVAMKEEAAGEVPVAFVVKSKDSELSEDDVKQFVSKQVVFYKRINKVFFTESIPKAPS<br>GKILRKDLRAKLANGL |
| His$_6$-Thrombin-*At*ACS$_{mut}$ [T324G, V399A, W427G] | **M**GSSHHHHHHSSGLVPRGSMASEENDLVFPSKEFSGQALVSSPQQYMEMHKRSMDDPAAFWS<br>DIASEFYWKQKWGDQVFSENLDVRKGPISIEWFKGGITNICYNCLDKNVEAGLGDKTAIHWEG<br>NELGVDASLTYSELLQRVCQLANYLKDNGVKKGDAVVIYLPMLMELPIAMLACARIGAVHSVV<br>FAGFSADSLAQRIVDCKPNVILTCNAVKRGPKTINLKAIVDAALDQSSKDGVSVGICLTYDNSLA<br>TTRENTKWQNGRDVWWQDVISQYPTSCEVEWVDAEDPLFLLYTSGSTGKPKGVLHTTGGYMI<br>YTATTFKYAFDYKSTDVYWCTADCGWIGGHSYVTYGPMLNGATVVVFEGAPNYPDPGRCWDI<br>VDKYKVSIFYTAPTLVRSLMRDDDKFVTRHSRKSLRVLGSAGEPINPSAWRWFFNVVGDSRCPI<br>SDTWGQTETGGFMITPLPGAWPQKPGSATFPFFGVQPVIVDEKGNEIEGECSGYLCVKGSWPGA<br>FRTLFGDHERYETTYFKPFAGYYFSGDGCSRDKDGYYWLTGRVDDVINVSGHRIGTAEVESAL<br>VLHPQCAEAAVVGIEHEVKGQGIYAFVTLLEGVPYSEELRKSLVLMVRNQIGAFAAPDRIHWAP<br>GLPKTRSGKIMRRILRKIASRQLEELGDTSTLADPSVVDQLIALADV |

**Table S6: Nucleotide sequences of proteins used in this study.**

| Name | Nucleotide Sequence |
|---|---|
| $His_6$-Thrombin-*Zo*PKS1 | **ATG**GGCAGCAGCCATCATCATCATCATCACAGCAGCGGCCTGGTGCCGCGCGGCA<br>GCATGGGCAGCCTGCAAGCGATGCGTCGCGCGAAACGCGCGCAAGGCCCGGCGA<br>CCATTATGGCGGTGGGCACCGCGAACCCGCCGAACCTGTATGAACAGACGAGCTA<br>TCCGGATTTTTATTTTCGCGTGACCAACAGCGATGATAAACATGAACTGAAAAACA<br>AATTTCGCGTGATTTGCGAAAAAACCCGCGTGAAACGCCGCTATCTGCATCTGACC<br>GAAGAAATTCTGAAACAGCGCCCGAAACTGTGCAGCTATATGGAACCGAGCTTTG<br>ATGATCGCCAAGATATTGTGGTGGATGAAATTCCGAAACTGGCGAAAGAAGCGGC<br>GGAAAAAGCGATTAAAGAATGGGGCCATCCGAAAAGCGAAATTACCCATCTGGTG<br>TTTTGCAGCATTAGCGGCATTGATATGCCGGGCGCGGATTATCGCCTGGCGAAACT<br>GCTGGGCCTGCCGCTGAGCGTGAACCGCCTGATGCTGTATAGCCAAGCGTGCCATA<br>TGGGCGCGCAGATGCTGCGCATTGCGAAAGATCTGGCGGAAAACAACCGCGGCGC<br>GCGCGTGCTGGCGGTGAGCTGCGAAATTACCGTGCTGAGCTTTCGCGGCCCGGAT<br>GCGGGCGATTTCGAAGCGCTGGCCTGCCAAGCGGGCTTTGGCGACGGGGCGGCG<br>GCAGTAGTTGTGGGGGCGGATCCGCTGCCGGGCGTGGAACGCCCGATTTATGAAA<br>TTGCGGCGGCGATGCAAGAAACCGTGCCGGAAAGCGAACGCGCGGTGGGCGGCC<br>ATCTGCGCGAAATTGGCTGGACCTTTCATTTTTTTAATCAGCTGCCGAAACTGATTG<br>CGGAAAACATTCAGAGCAGCCTGGCGCGCGCGTTTAAACCGCTGGGCATTACCGA<br>ATGGAACGATGTGTTTTGGGTGGCGCATCCGGGCAACTGGGGCATTATGGATGCGA<br>TTGAAACCAAACTGGGCCTGAAACAAGGCAAACTGGCGACCGCGCGCCATGTGTT<br>TAGCGAATATGGCAACATGCAGAGCGCGACCGTGTATTTTGTGATGGATGAAGTGC<br>GCAAACGCAGCGCGGCGGAAGGCCGCGCGACCACCGGCGAAGGCCTGGAATGGG<br>GCGTGCTGTTTGGCTTTGGCCCGGGCCTGACCATTGAAACCGTGGTGCTGCGCAG<br>CCTGCCGATTCCGTAA |
| $His_6$-Thrombin-*Zo*PKS2 | **ATG**GGCAGCAGCCATCATCATCATCATCACAGCAGCGGCCTGGTGCCGCGCGGCA<br>GCATGGCGAGCATTAACAACGTAGTGGTGGATGCTTTCCCAAAACCGCAACGTGC<br>GCGCGGACCGGCGACAGTGATGGCGATTGGCACCGCGAACCCGCCGAACCTGTAT<br>GAACAGAGCAGCTATCCGGATTTTTATTTTCGCGTGACCAACAGCGATCATAAACC<br>GGAACTGAAACAGAAATTTCGCCGCCTGTGCGAACGCAGCATGATTAAAAAACGC<br>TATATGCATCTGACCGAAGAACTGCTGAAAGAAAAACCGGGCATGTGCAGCTATAT<br>GGATACGAGCTTTGATGAACGCCAAGATATTGTGGTGGAAGAAGTGCCGCGCCTG<br>GCGAAAGAAGCGGCGGTGAAAGCGATTAAAGAATGGGGCCGCAGCAAAAGCGAA<br>ATTACCCATCTGGTGTTTTGCAGCACGAGCGGCGTGGATATGCCGGGCGCGGATTA<br>TCGCCTGGCGAACCTGCTGGGCCTGAGCAGCAGCGTGAACCGCATTATGCTGTATA<br>ACCAAGCGTGCCATATTGGCGCGCAGACCCTGCGCATTGCGAAAGATATTGCGGAA<br>AACAACCGCACCGCGCGCGTGCTGGTGGTGGCGTGCGAAGTGAACACCCTGATTT<br>TTCGCGGCCCGGAAGAACGCGATTTTCAGAGCTTGGCGGCGCAAGTGGCGTTCGG<br>AGATGGAGCGGCGGCAGTGGTTGTAGGCGCGGATCCGGTGGAAGGCGTGGAACG<br>CCCGATTTTTGAAATTATGGCGGCGCTGCCGTTTACCGTGCCGGAAACGCAGATGG<br>CGGTGGGCGGTCAGCTGAAACAGATTGGCCTGACCTTTCATTTTGCGCATCAGCTG<br>CCGGGCCTGATTGCGAACAACCTGGAAACCTGCCTGGGCGAAGCGCTGAAACCG<br>CTGGGCATTAGCGATTGGAACGATGTGTTTTGGGTGGCGCATCCGGGCAACTGGG<br>GCATTATGGATGCGGTGGAAGCGAAACTGGGCCTGGAACAAGGCAAACTGCAGA<br>GCAGCCGCCATGTGTTTAGCGAATTTGGCAACATGATGAGCGCGACCGTGCTGTTT<br>GTGATGGATGAGGTGCGTAAGCGCGCGGCGGCGAAAGGCGCGGCAACCACCGGA<br>GACGGCTTACAGTGGGGAGTGCTGTGCGGCTTTGGCCCGGGCCTGAGCATTGAAA<br>CCCTGGTGCTGCGCAGCGTGCCGCTGTAA |
| $His_6$-Thrombin-*Zo*DCS1 | **ATG**GGCAGCAGCCATCATCATCATCATCACAGCAGCGGCCTGGTGCCGCGCGGCA<br>GCATGGCTGCTGTGATGGAAGCATTTTCACGTACCCCGCCGGCGGATGGCGCGGCC<br>AACGTGCTGGCGATTGGCACCGCGAACCCGAGCCATTTTGTGGATCAGATGCAGT<br>ATCCGGAATATTATTTTCGCATTACCGATGCGGAAGGCAAAACCGAACTGCAGCAG<br>AAATTTAAACGCATTTGCGAAAAAAGCATGATTCGCAAACGCCATATGTGCCTGAC<br>CGAAGAAGTGCTGCGTGAAAATCCGTGCCTGTGCGGCTATATGACCCCGAGCTTTG<br>ATGCGCGTCAGCGCATTGTGGTGGAAGAAGTGCCGCGCCTGGCGAAAGAAGCGG<br>CGGATAAAGCGATTAAAGAATGGGGCCATCCGGTGACCGATATTACCCATCTGGTG<br>TTTTGCAGCGCGGCGGGCGTGGATCTGCCGGGCGCGGATTATAGCCTGCTGCAGCT<br>GCTGGGCCTGCCGCTGCATGTGCGCCGCGTGATGCTGTATAACGTGGGCTGCCATG<br>CGGGCGGCACCGCGCTGCGCGTGGCGAAAGATCTGGCGGAAAACAACAAAGGCG<br>CGCGCGTGCTGGTGGTGTGCAGCGAACTGAACGTGATGTTTTTCGCGGCCCGGG |

| | |
|---|---|
| | CGATGATCATATTGAAAACCTGATTGGCCAAGCGCTGTTTGGCGATGGCGCGGCGG<br>CCGTGATTGTGGGCGCGGATACCGATGAAACCGAACGCCCGATTTATGAAGTGGCG<br>AGCGCGACCCAAGTGATGCTGCCGGAAAGCGAAGAAATGGTGGGCGGCCATCTG<br>CGCGAAATTGGCCTGACCTTTCATCTGGCGAGCCGCCTGCCGGCGGTGGTGGGCG<br>AAAACATTGAACGCTGCCTGGAAAGCGCGTTTGGCGTGGAAGCGGGCGATTGGA<br>ACGAACTGTTTTGGATTGTGCATCCGGGCGGCCGCGCGATTATTGATCAAGTGGAA<br>GCGCGCGTGCGCCTGCGCCCGGAAAAACTGGCGGCGACCCGCCATGTATTGCGCG<br>AATATGGCAACATGCAGAGCGCGAGCGTGCTGTTTATTATGGATGAAATGCGCAAA<br>CTGAGCGCGGCGGAAGGCTGCGCGACCACCGGCCAAGGCTGTCAGTGGGGCGTG<br>CTGTTTGGCTTTGGCCCGGGCCTGACCATTGAAACCGTGGTGCTGCGCAGCGTGC<br>CGCGTCAGATTAACTAA |
| His$_6$-Thrombin-*Zo*DCS2 | **ATG**GGCAGCAGCCATCATCATCATCATCACAGCAGCGGCCTGGTGCCGCGCGGCA<br>GCATGGCGAGCACAGAAGCTTTCCGCCGCGCGCCGCCGGCGGATGGCCCGGCAAC<br>CGTGCTGGCGATTGGCACCGCGAACCCGAGCCATTTTGTGGATCAGATGCAGTATC<br>CGGATTATTATTTTCGCGTGACCAACGCGGAAGATAAAACCGAACTGAAACAGAA<br>ATTTAAACGCATTTGCGAAAAAAGCACCATTCGCAAACGCCATATGTGCCTGACCG<br>AAGAAATTCTGAAAGAAAACCCGAGCCTGTGCGCGTATATGGCGCCGAGCTTTGA<br>TGCGCGCCAAGGCATTGTGCTGGAAGAAGTGCCGCGCCTGGCGAAAGAAGCGGC<br>GGATAAAGCGATTAAAGAATGGGGCCGCCCGGTGAGCGATGTGACCCATCTGGTG<br>TTTTGTAGCGCGGCGGGCGTGGATCTGCCGGGCGTGGATTATCGCCTGATTCAGCT<br>GCTGGGCCTGCCGGCGCGCGTGCGCCGCGTGATGCTGTATAACGTGGGCTGCCATG<br>CGGGCGGCACCGCGCTGCGCGTGGCGAAAGATCTGGCGGAAAACAACAAAGGCG<br>CGCGCGTGCTGGTGGTGTGCAGCGAACTGAACGTGATGTTTTTCGCGGCCCGGA<br>TGATCATCATTTTGAAAACCTGATTGGCCAAGCGCTGTTTGGCGATGGCGCGGCGG<br>CGCTGATTGTGGGCGCGGATCCGGAAGAAGCGGAACGCGCGATTTATGAAGTGGC<br>GAGCGCGACCCAAGTGATGCTGCCGGAAAGCGAAGAAATGGTGGGCGGCCATCT<br>GCGCGAAATTGGCCTGACCTTTCATCTGGCGAGCAAACTGCCGGCGGTGGTGGGC<br>GGCAACATTGAACGCTGCCTGGAAGCGGCGTTTGGCCCGCAAGCGGGCGTGGCG<br>GATTGGAACGAACTGTTTTGGATTGTGCATCCGGGCGGCCGCGCGATTATTGATCA<br>AGTGGAAGCGCGCGCGGGACTGACCGCGGAAAAACTGGCGGTGACCCGCCATGT<br>GCTGCGCGAATATGGCAACATGCAGAGCGCGAGCGTGCTGTTTATTATGGATGAAA<br>TGCGCAAACGTAGCGCGGCGGAAGGCTGCGCGACCACCGGCCAAGGCTGTCAGT<br>GGGGCGTGCTGTTTGGCTTTGGCCCGGGCCTGACCGTGGAAACCGTGGTGCTGCG<br>CAGCGTGCCGATTAAACTGATTAACTAA |
| His$_6$-Thrombin-*At*4Cl1 | **ATG**CATCACCACCATCACCATAGCTCTGGCCTGGTGCCGCGCGGCAGCCATATGAT<br>GGGTTCAGGGGGATCCGGTGTCGACATGGAGGAGGATTACAAAATGGCGCCACAA<br>GAACAAGCAGTTTCTCAGGTGATGGAGAAACAGAGCAACAACAACAACAGTGAC<br>GTCATTTTCCGATCAAAGTTACCGGATATTTACATCCCGAACCACCTATCTCTCCAC<br>GACTACATCTTCCAAAACATCTCCGAATTCGCCACTAAGCCTTGCCTAATCAACGG<br>ACCAACCGGCCACGTGTACACTTACTCCGACGTCCACGTCATCTCCCGCCAAATCG<br>CCGCCAATTTTCACAAACTCGGCGTTAACCAAAACGACGTCGTCATGCTCCTCCTC<br>CCAAACTGTCCCGAATTCGTCCTCTCTTTCCTCGCCGCCTCCTTCCGCGGCGCAAC<br>CGCCACCGCCGCAAACCCTTTCTTCACTCCGGCGGAGATAGCTAAACAAGCCAAA<br>GCCTCCAACACCAAACTCATAATCACCGAAGCTCGTTACGTCGACAAAATCAAAC<br>CACTTCAAAACGACGACGGAGTAGTCATCGTCTGCATCGACGACAACGAATCCGT<br>GCCAATCCCTGAAGGCTGCCTCCGCTTCACCGAGTTGACTCAGTCGACAACCGAG<br>GCATCAGAAGTCATCGACTCGGTGGAGATTTCACCGGACGACGTGGTGGCACTAC<br>CTTACTCCTCTGGCACGACGGGATTACCAAAAGGAGTGATGCTGACTCACAAGGG<br>ACTAGTCACGAGCGTTGCTCAGCAAGTCGACGGCGAGAACCCGAATCTTTATTTCC<br>ACAGCGATGACGTCATACTCTGTGTTTTGCCCATGTTTCATATCTACGCTTTGAACT<br>CGATCATGTTGTGTGGTCTTAGAGTTGGTGCGGCGATTCTGATAATGCCGAAGTTTG<br>AGATCAATCTGCTATTGGAGCTGATCCAGAGGTGTAAAGTGACGGTGGCTCCGATG<br>GTTCCGCCGATTGTGTTGGCCATTGCGAAGTCTTCGGAGACGGAGAAGTATGATTT<br>GAGCTCGATAAGAGTGGTGAAATCTGGTGCTGCTCCTCTTGGTAAAGAACTTGAA<br>GATGCCGTTAATGCCAAGTTTCCTAATGCCAAACTCGGTCAGGGATACGGAATGAC<br>GGAAGCAGGTCCAGTGCTAGCAATGTCGTTAGGTTTTGCAAAGGAACCTTTTCCG<br>GTTAAGTCAGGAGCTTGTGGTACTGTTGTAAGAAATGCTGAGATGAAAATAGTTGA<br>TCCAGACACCGGAGATTCTCTTTCGAGGAATCAACCCGGTGAGATTTGTATTCGTG<br>GTCACCAGATCATGAAAGGTTACCTCAACAATCCGGCAGCTACAGCAGAGACCAT<br>TGATAAAGACGGTTGGCTTCATACTGGAGATATTGGATTGATCGATGACGATGACG<br>AGCTTTTCATCGTTGATCGATTGAAAGAACTTATCAAGTATAAAGGTTTTCAGGTAG<br>CTCCGGCTGAGCTAGAGGCTTTGCTCATCGGTCATCCTGACATTACTGATGTTGCTG |

| | |
|---|---|
| | TTGTCGCAATGAAAGAAGAAGCAGCTGGTGAAGTTCCTGTTGCATTTGTGGTGAA<br>ATCGAAGGATTCGGAGTTATCAGAAGATGATGTGAAGCAATTCGTGTCGAAACAG<br>GTTGTGTTTTACAAGAGAATCAACAAAGTGTTCTTCACTGAATCCATTCCTAAAGC<br>TCCATCAGGGAAGATATTGAGGAAAGATCTGAGGGCAAAACTAGCAAATGGATTG<br>TGA |
| His$_6$-Thrombin-<br>*At*ACS$_{mut}$<br>[T324G,<br>V399A,<br>W427G] | ATGGGCAGCAGCCATCATCATCATCATCACAGCAGCGGCCTGGTGCCGCGCGGCA<br>GCATGGCGAGCGAAGAAAACGATCTGGTGTTTCCGAGCAAAGAATTTAGCGGCCA<br>AGCGCTGGTGAGCAGCCCGCAGCAGTATATGGAAATGCATAAACGCAGCATGGAT<br>GATCCGGCGGCGTTTTGGAGCGATATTGCGAGCGAATTTTATTGGAAACAGAAATG<br>GGGCGATCAAGTGTTTAGCGAAAACCTGGATGTGCGCAAAGGCCCGATTAGCATT<br>GAATGGTTTAAAGGCGGCATTACCAACATTTGCTATAACTGCCTGGATAAAAACGT<br>GGAAGCGGGCCTGGGCGATAAAACCGCGATTCATTGGGAAGGCAACGAACTGGG<br>CGTGGATGCGAGCCTGACCTATAGCGAACTGCTGCAGCGCGTGTGTCAGCTGGCG<br>AACTATCTGAAAGATAACGGCGTGAAAAAAGGCGATGCGGTGGTGATTTATCTGCC<br>GATGCTGATGGAACTGCCGATTGCGATGCTGGCGTGCGCGCGCATTGGCGCGGTGC<br>ATAGCGTGGTGTTTGCGGGCTTTAGCGCGGATAGCCTGGCGCAGCGCATTGTGGAT<br>TGCAAACCGAACGTGATTCTGACCTGCAACGCGGTGAAACGCGGCCCGAAAACC<br>ATTAACCTGAAAGCGATTGTGGATGCGGCGCTGGATCAGAGCAGCAAAGATGGCG<br>TGAGCGTGGGCATTTGCCTGACCTATGATAACAGCCTGGCGACCACCCGCGAAAA<br>CACCAAATGGCAGAACGGCCGCGATGTGTGGTGGCAAGATGTGATTAGTCAGTATC<br>CGACGAGCTGCGAAGTGGAATGGGTGGATGCGGAAGATCCGCTGTTTCTGCTGTA<br>TACGAGCGGCAGCACCGGCAAACCGAAAGGCGTGCTGCATACCACCGGCGGCTAT<br>ATGATTTATACCGCGACCACCTTTAAATATGCGTTTGATTATAAAGCACCGATGTGT<br>ATTGGTGCACCGCGGATTGCGGCTGGATTGGCGGCCATAGCTATGTGACCTATGGC<br>CCGATGCTGAACGGCGCGACCGTGGTGGTGTTTGAAGGCGCGCCGAACTATCCGG<br>ATCCGGGCCGCTGCTGGGATATTGTGGATAAATATAAAGTGAGCATTTTTTATACCG<br>CGCCGACCCTGGTGCGCAGCCTGATGCGCGATGATGATAAATTTGTGACCCGCCAT<br>AGCCGTAAAAGCCTGCGCGTGCTGGGCAGCGCGGGCGAACCGATTAACCCGAGC<br>GCGTGGCGCTGGTTTTTTAACGTGGTGGGCGATAGCCGCTGCCCGATTAGCGATAC<br>CTGGGGTCAGACCGAAACCGGCGGCTTTATGATTACCCCGCTGCCGGGCGCGTGG<br>CCGCAGAAACCGGGCAGCGCGACCTTTCCGTTTTTTGGCGTGCAGCCGGTGATTG<br>TGGATGAAAAAGGCAACGAAATTGAAGGCGAATGCAGCGGCTATCTGTGCGTGAA<br>AGGCAGCTGGCCGGGCGCGTTTCGCACCCTGTTTGGCGATCATGAACGCTATGAA<br>ACCACCTATTTTAAACCGTTTGCGGGCTATTATTTTAGCGGCGATGGCTGCAGCCGC<br>GATAAAGATGGCTATTATTGGCTGACCGGCCGCGTGGATGATGTGATTAACGTGAG<br>CGGCCATCGCATTGGCACCGCGGAAGTGGAAAGCGCGCTGGTGCTGCATCCGCAG<br>TGCGCGGAAGCGGCGGTGGTGGGCATTGAACATGAAGTGAAAGGCCAAGGCATTT<br>ATGCGTTTGTGACCCTGCTGGAAGGCGTGCCGTATAGCGAAGAACTGCGTAAAAG<br>CCTGGTGCTGATGGTGCGCAATCAGATTGGCGCGTTTGCGGCGCCGGATCGCATTC<br>ATTGGGCGCCGGGCCTGCCGAAAACCCGCAGCGGCAAAATTATGCGCCGCATTCT<br>GCGCAAAATTGCGAGCCGTCAGCTGGAAGAACTGGGCGATACGAGCACCCTGGC<br>GGATCCGAGCGTGGTGGATCAGCTGATTGCGCTGGCGGATGTGTAA |
